# The entorhinal spatial map integrates visual identity information of landmarks

**DOI:** 10.1101/2025.03.13.643169

**Authors:** Garret Wang, Farid Shahid, Taylor Malone, Jean Tyan, Kyle Cekada, Yi Gu

## Abstract

The location and identity of landmarks provide spatial and nonspatial information of an environment, respectively, to guide spatial navigation. While the medial entorhinal cortex (MEC) is essential for representing spatial information, it is unclear whether it also encodes landmark identity. Here, we conducted two-photon calcium imaging of the MEC when mice navigated in multiple virtual environments, and discovered a general ability of the MEC to encode landmark identity through cue cells, which responded to individual landmarks during virtual navigation. Cue cells represented landmark identity by exhibiting more distinct activity patterns between visually disparate landmarks than identical ones. The representation was modulated by the spatial shift of cue cell activity relative to landmark location. Moreover, the identity encoding by the same cue cell population changed between different environments but was maintained within the same environment despite increased experience. In contrast, landmark location encoding by cue cells was regulated by experience, suggesting different mechanisms underlying the encodings of landmark identity and location. Finally, compared to cue cells, grid cells weakly encoded landmark identity but more robustly encoded landmark location. Thus, the MEC integrates both spatial and nonspatial information during navigation, but potentially through different circuit mechanisms.

## Introduction

Landmarks convey important information for spatial navigation in an environment^1^. Landmark location (“where”) provides spatial information, whereas landmark identity (“what”), such as whether two landmarks look similar or different, provides nonspatial information. However, it is unclear how spatial and nonspatial information of landmarks is integrated in the same cognitive map. Classical models propose that neocortical inputs carrying spatial and nonspatial information separately arrive at the medial (MEC) and lateral entorhinal cortices (LEC), respectively. Both regions project to the hippocampus, which combines the two information types into a complete cognitive map^2–4^. Consistent with the models, MEC dysfunction or lesion led to deficits in spatial navigation and memory^5–9^ but not nonspatial memory^6^. The MEC is also abundant with spatially-modulated cells, such as grid cells with triangular firing pattern in an open arena^10^, and cells representing animals’ head direction ^11^ and environmental borders^12^.

However, some lesion studies suggest that the MEC is also involved in item recognition in spatial environments^13,14^ and it was active at similar levels in spatial and nonspatial tasks based on its immediate early gene expression^15^. In environments with comparable spatial information but spatial cues in visual and auditory modalities, the MEC exhibited different cognitive maps^16^. MEC activity also differentiated reward cues with different identity ^17^. Therefore, nonspatial landmark identity could potentially be encoded by the MEC. Along with this idea, the MEC contains object-vector cells and cue cells, which specifically respond to objects/visual landmarks during navigation in real^18^ and virtual environments^19^, respectively. However, these cells are thought to provide vectorial representation of an animal’s location relative to landmarks, because they are active at consistent distances and directions from all salient objects, regardless of their identity^18,19^. Thus, whether MEC neurons compute landmark identity during spatial navigation remains unanswered.

To directly investigate the MEC encoding of landmark identity during navigation, we measured MEC calcium dynamics, which approximated its neural activity, as mice navigated in multiple virtual reality (VR) linear tracks containing visual landmarks with identical and disparate appearances, which corresponded to the same and different identity, respectively. We hypothesized that if MEC neurons encode landmark identity, they should exhibit a larger activity difference between disparate than identical landmarks (Fig. 1A, a). Otherwise, the differences between these landmarks should be comparable (Fig. 1A, b). Since the activity difference could be confounded by the encoding of landmark distance and location, we eliminated the effect of landmark distance by specifically comparing the activity difference between identical and disparate landmark pairs at matched distances. We further eliminated the effect of landmark location by repeating the comparison across multiple tracks with many landmark pairs at various locations. Thus, we uncovered the encoding of landmark identity by cue cells, which exhibited greater activity difference between disparate landmarks than identical ones. The encoding was modulated by the spatial shift of cue cell activity relative to landmark location and by track identity, but was not influence by increased track experience. However, the encoding of landmark location by cue cells was regulated by experience. In contrast to cue cells, grid cells weakly represented landmark identity but more robustly encoded landmark location. Therefore, during navigation, the MEC encodes not only spatial, but also nonspatial, information, potentially through different circuit mechanisms.

**Figure 1.**
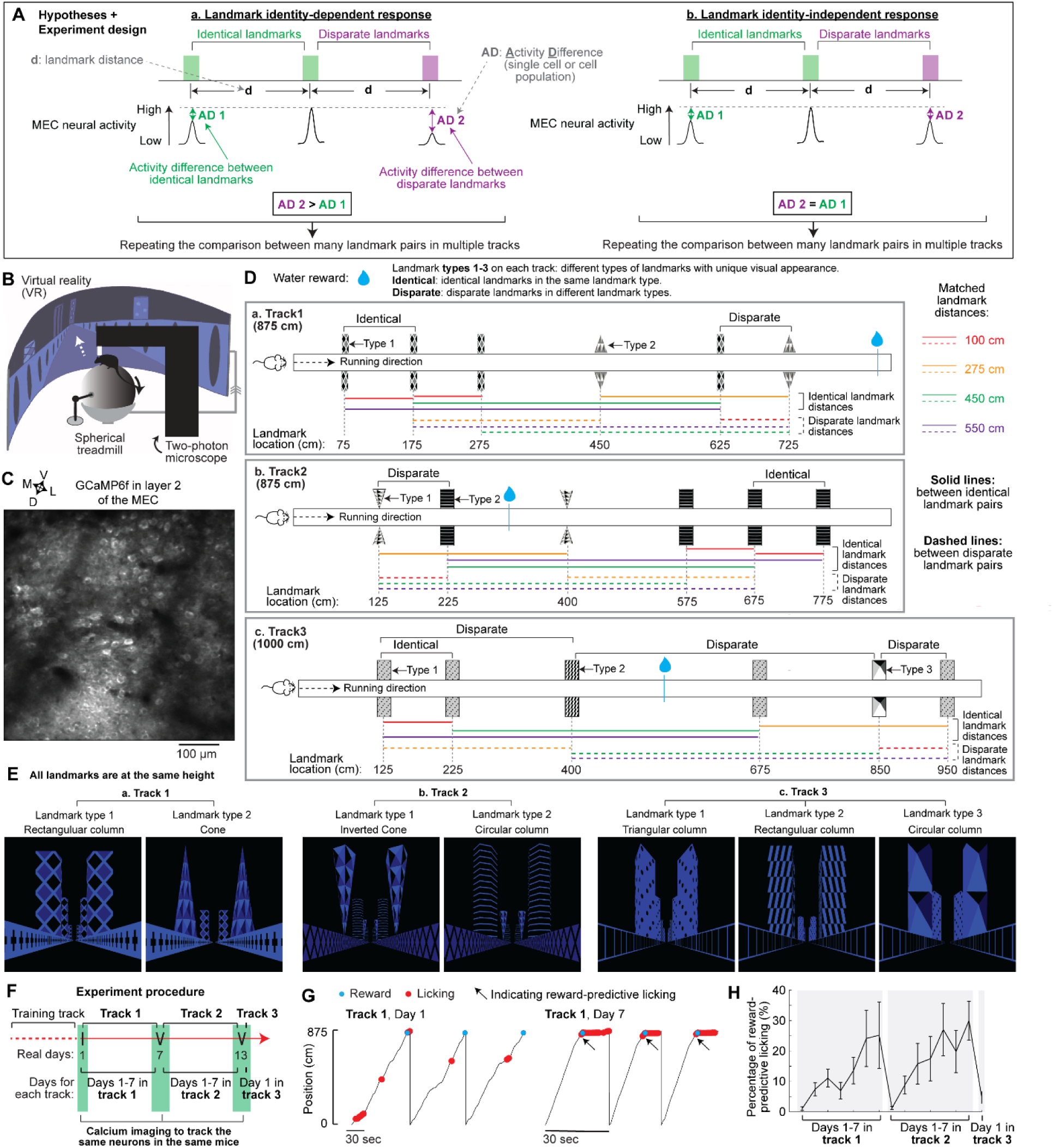
Experiment design. **A.** Hypothesis about the encoding of landmark identity. **B.** Experimental setup. **C.** Example two-photon FOV showing excitatory neurons in layer 2 of the MEC expressing GCaMP6f. V: ventral; D: dorsal; M: medial; L: lateral. **D.** The design of tracks 1-3. **E.** Example views of tracks 1-3 from a mouse’s perspective, showing different types of landmarks. **F.** Experimental procedure. Note that on real days 7 and 13, the mouse performed in two sessions, which are considered as day 7 of track 1 and day 1 of track 2 (day 7) or day 7 of track 2 and day 1 of track 3 (day 13). **G.** Behavioral examples of a mouse on days 1 and 7 in track 1. Black arrows indicate the occurrence of reward-predictive licking before the reward. **H.** The percentage of reward-predictive licking in tracks 1-3. Gray blocks highlight the days in the same track. The percentage was calculated as the percentage of licking events right before the reward location among those in the whole track, excluding the licking for reward consumption. Error bars represent mean ± SEM. Same for all figures.

## Results

### Experiment design for investigating the encoding of landmark identity

We conducted cellular-resolution two-photon calcium imaging of the MEC in head-fixed mice while they unidirectionally navigated linear VR tracks for multiple runs via teleportation at the end of the tracks^20^ (Fig. 1B). GP5.3 transgenic mice were used to enable the imaging of stably expressed fluorescent calcium indicator GCaMP6f^21^ in MEC layer 2 excitatory neurons, which are comprised of many functional cell types for navigation, such as grid cells and cue cells^19,22^ (Fig. 1C).

As proposed in Fig. 1A, we designed three VR tracks with visual landmarks symmetrically arranged on both sides of each track (Fig. 1D). Track 1 was 875 cm long and contained two sets of landmarks, which had identical appearance (i.e., shapes and surface pattern) within the same set (“identical landmarks”) and distinct appearance across sets (“disparate landmarks”) (Fig. 1D and E, a). The landmarks were repeated at multiple locations so that four matched distances (100, 275, 450, and 550 cm) were created for both the identical and disparate pairs. The four distances were also preserved for different sets of identical and disparate landmarks at distinct locations in tracks 2 (875 cm) and 3 (1000 cm) (Fig. 1D and E, b and c). Tracks 1-3 also had different wall patterns and reward locations. Thus, tracks 1-3 together enabled our study on the general ability of the MEC to encode landmark identity (identical versus disparate landmarks), independent of landmark distance, location, and other track features.

After the initial acclimation to VR navigation in a training track, water-restricted GP5.3 mice explored tracks 1-3, where they received a water reward upon reaching a specific location in each track. The mice were exposed to tracks 1 and 2 for seven days per track and to track 3 for one day (Fig. 1F). During the seven days in tracks 1 and 2, the mice gradually improved their specificity to predictively lick right before the reward, indicating their learning of the tracks^23,24^ (Fig. 1G and H). Calcium imaging was conducted on days 1 and 7 in tracks 1 and 2 and on day 1 in track 3. The same layer 2 excitatory neurons in the MEC of each mouse were tracked across the five sessions.

Overall, the above setting allowed us to investigate the MEC encoding of landmark identity in multiple tracks (tracks 1-3) and during learning of the same track (tracks 1 and 2).

### Encoding of landmark identity by cue cells at the individual-cell level

We first asked whether MEC neural response differentiated identical and disparate landmarks on day 1 in tracks 1-3. We focused on cue cells as they have highly specific activity toward landmarks. Cue cells were identified in each track based on cue scores, which quantified their landmark-dependent activation^19^. Around 20% of active cells were cue cells (track 1: 16.5 ± 1.9 %; track 2: 22.0 ± 2.0 %; track 3: 22.1 ± 2.1%). As previously reported^19^, each cue cell exhibited a characteristic “spatial shift”, reflecting the distance and direction (before, at, or after) of cue cell activity relative to landmarks (Fig. 2A). The activity and the “landmark template”, which represented the pattern of landmark arrangement, were best matched after shifting the template according to the spatial shift. Sorted activities of cue cells by their shifts formed consistent sequences around individual landmarks^19^ (Fig. 2B).

**Figure 2.**
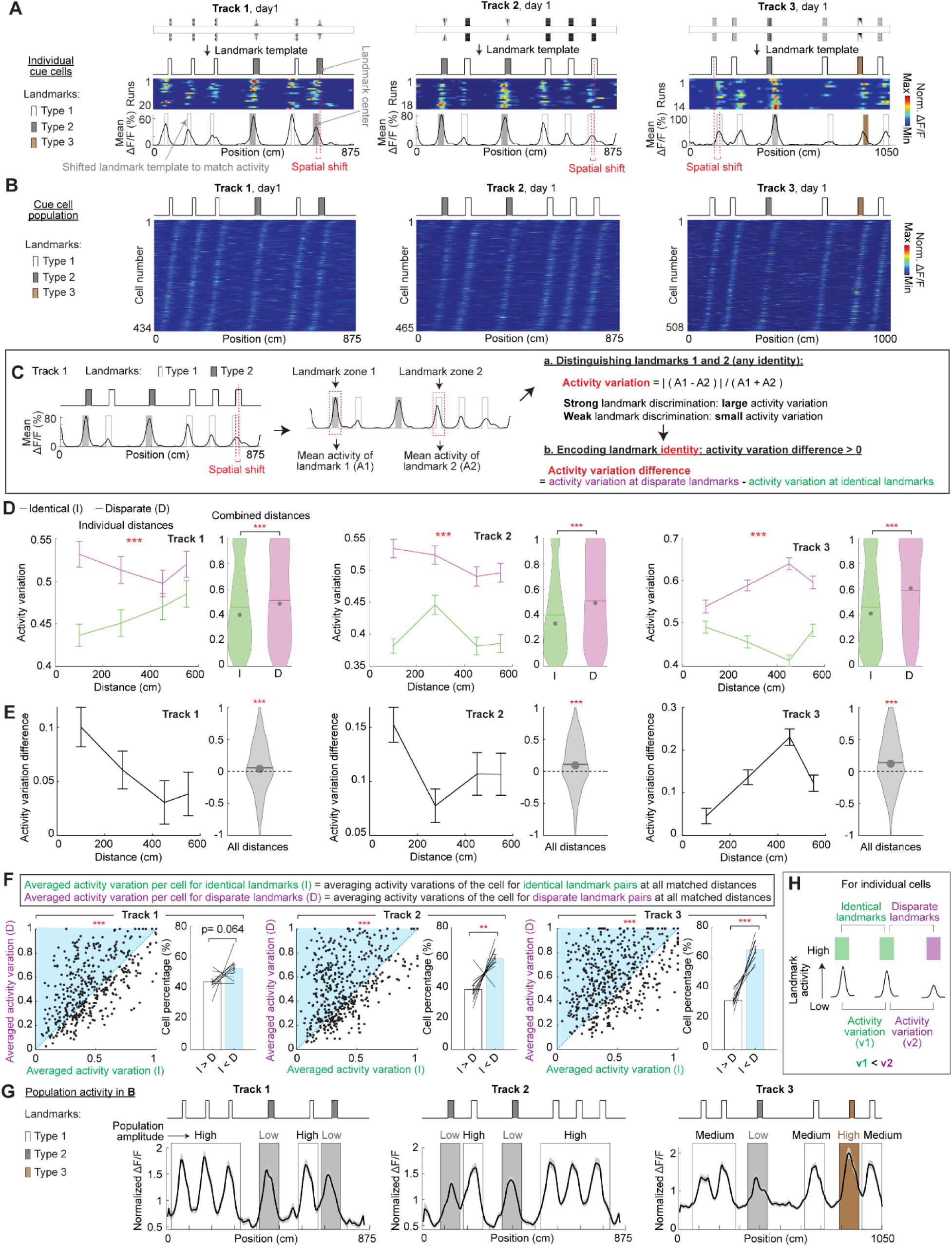
Landmark identity encoding at the individual-cell level. **A.** Examples of cue cells in individual tracks. For each cell: from top to bottom: real track; simplified landmark template with different landmark types; run-by-run calcium dynamics; mean ΔF/F (spatially binned) along the track. The landmark template is aligned to the mean ΔF/F under a spatial shift (the distance between red dashed lines). **B.** Cue cells in individual tracks. Cue cell activities were sorted by their spatial shifts relative to landmark template. The activity of each cell is normalized by its mean. **C.** The calculations of “activity variation” and “activity variation difference”. **D.** Activity variation for identical (I, green) and disparate landmark pairs (D, magenta) in tracks 1 to 3 (left to right) at individual (left) and combined matched distances (right). The asterisks in left panels indicate the results of two-way ANOVA for I and D curves. The asterisks in the right panels indicate the results of two-tailed t tests. Same below. **E.** Activity variation difference in the three tracks at individual (left) and combined distances (right). The asterisks in right panels indicate a comparison with zero. Same below. **F.** For each track: left: “averaged activity variations” for identical landmark pairs (I) versus those for disparate pairs (D) in individual cells. Blue and white zones include cells with larger variations for disparate and identical landmark pairs, respectively. Right: the percentage of cells per FOV with larger activity variations for identical (white) or disparate landmark pairs (blue). **G.** The mean (black curve) and standard errors (gray shade) of all cell activity in each track in B. The activity for each landmark is highlighted by the box with the landmark color and its relative amplitude is indicated above. **H.** A summary for activity variation of individual cells at identical and disparate landmark pairs. *p ≤ 0.05, **p ≤ 0.01, ***p ≤ 0.001, n.s. p > 0.05. Significant p values are in Supplementary Table 1. In violin plots, the dot and horizontal bar represent the median and mean, respectively. Same for all figures.

We examined the encoding of landmark identity at the individual-cell level by evaluating activity variation of each cue cell between landmark pairs (Fig. 2C). We aligned the activity of a cell with landmark template based on its spatial shift and identified its activity zone around each landmark (“landmark zone”). Its “landmark activity” was the mean activity in each landmark zone, and its “activity variation” between a landmark pair was the absolute difference over the sum of its “landmark activities” at the two landmarks (Fig. 2C, a). Large and small activity variations indicate stronger and weaker landmark discrimination, respectively. If a cue cell encodes landmark identity, it should exhibit larger activity variation between disparate than identical landmarks. Consequently, its “activity variation difference”, which is its activity variation between disparate landmarks subtracted by that between identical ones at matched distances, is above zero (Fig. 2C, b) Across the three tracks, activity variation of cue cells (grouping cells from all mice) was larger between disparate than identical landmarks at both individual and combined distances (Fig. 2D), suggesting that the activity variation encodes landmark identity. Additionally, there was no consistent relationship between the variation and landmark distance for either identical or disparate pairs (Fig. 2D, the trends of magenta and green curves), indicating that the variation does not represent landmark distance. As expected, activity variation difference was above zero (Fig. 2E, right for each track) and did not consistently change with landmark distance (Fig. 2E, left curve for each track), suggesting that landmark identity encoding is also distance-independent. The above results remained true after grouping the cells by imaging field of view (FOV) (Fig. S1A and B) and by mouse (Fig. S1C and D), confirming that the results were not dominated by cells in particular FOVs or mice, despite the variation in cue cell numbers (Fig. S1E and F).

We further evaluated landmark identity encoding in each cue cell by comparing its “averaged activity variations” for identical and disparate landmark pairs across all matched distances. The averaged variations for the two types of landmark pairs were positively correlated (Fig. 2F, left for each track), but more cells showed greater variations for disparate ones (Fig. 2F, right for each track). Therefore, while cue cells have a consistent ability to distinguish landmarks in general, more cells better discriminated disparate landmarks. Finally, activity variation differences between disparate and identical landmarks could also be directly visualized in the population activity of cue cells, which exhibited more distinct and similar activity amplitudes between disparate and identical landmarks in each track, respectively (Fig. 2G).

Therefore, individual cue cells better discriminate disparate than identical landmarks by exhibiting greater activity variation between the disparate ones (Fig. 2H).

### Encoding of landmark identity by cue cells at the population level

We next examined landmark identity encoding by cue cells at the population level (Fig. 3A). We hypothesized that relative activity levels between simultaneously recorded cue cells serve as a population code for each landmark, and the code is more different between disparate than identical landmarks. To quantify the code, we randomly picked a fixed number of cue cells (a “cue cell group”) in a FOV and ranked their “landmark activities” at each landmark. The similarity of the ranks at different landmarks was quantified by their correlation (“rank correlation”, Fig. 3A, a). Higher and lower correlations reflected similar and different codes for a pair of landmarks, corresponding to weaker and stronger landmark discrimination, respectively. If cue cells encode landmark identity, they should exhibit lower rank correlation between disparate than identical landmarks. Thus, their “rank correlation difference”, which was the rank correlation between disparate landmarks subtracted by that between identical ones at matched distances, should be below zero (Fig. 3A, b). For each FOV, the number of unique cue cell groups equaled the number of cells in the FOV (see Methods), enabling comparable statistical power of this calculation and that for activity variation (Fig. 2C).

**Figure 3.**
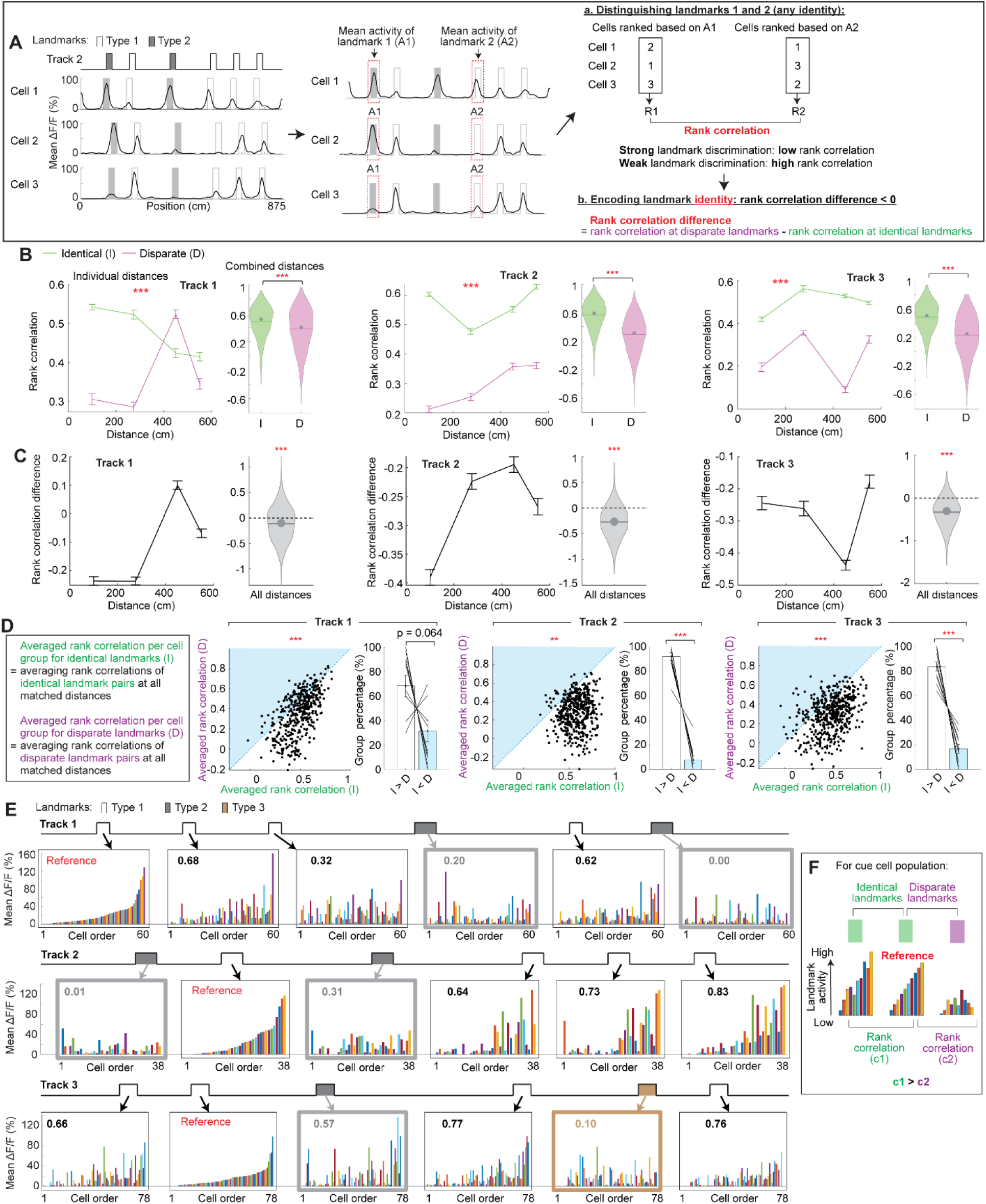
Landmark identity encoding at the population level. **A.** The calculations of “rank correlation” and “rank correlation difference”. **B.** Rank correlation for identical (I, green) and disparate landmark pairs (D, magenta) for tracks 1 to 3 (left to right). **C.** Rank correlation difference in the three tracks. Each cue cell group contained 15 cells. **D.** For each track: left: “averaged rank correlations” for identical landmark pairs (I) versus those for disparate pairs (D) for individual cell groups. Blue and white zones include cell groups with larger rank correlations for disparate and identical landmark pairs, respectively. Right: the percentage of cell groups per FOV with larger rank correlations for identical (white) or disparate landmark pairs (blue). **E.** Visualization of the activity of simultaneously imaged cue cells in the same FOV at individual landmarks. One FOV example is shown for each track. Cue cells were ranked ascendingly based on their “landmark activity” at the “reference” landmark and their activity at other landmarks were plotted in the same ranking. Rank correlation for these cells between each landmark and the reference, is shown on the top left of each landmark box. A higher rank correlation corresponds to more consistent trend of activity ascent at a landmark compared to the reference, indicating that the relative activities of the cells between the two landmarks are more similar. **F.** A summary of rank correlation of cue cell population at identical and disparate landmark pairs.

While grouping all cell groups (15 cells per group), rank correlations in tracks 1-3 were always lower between disparate than identical landmarks at individual and combined distances (Fig. 3B), suggesting that the population code represents landmark identity. There was no consistent association between rank correlation and landmark distance (Fig. 3B, the trends of magenta and green curves), indicating that the code does not represent landmark distance. Accordingly, rank correlation differences were below zero and were not consistently regulated by landmark distances (Fig. 3C). These results were largely preserved when grouping the cell groups by FOV (Fig. S2A and B) and by mouse (Fig. S2C and D). The rank correlation difference was also significantly below zero regardless of the number of cells per group (Fig. S2E). Therefore, the population code of cue cells better discriminates disparate landmarks.

For individual cue cell groups, their “averaged rank correlations” for identical and disparate landmark pairs across matched distances were positively correlated and more groups showed lower correlations for disparate landmarks (Fig. 3D), indicating the consistent ability of cue cell groups to discriminate both types of landmark pairs, but a stronger ability towards the disparate ones. Moreover, the difference in rank correlations for identical and disparate landmark pairs could be directly revealed by the relative activity of all simultaneously imaged cue cells in a FOV (Fig. 3E). We ranked cue cells in an ascending order based on their landmark activities at a reference landmark and arranged their activities at other landmarks in the same ranking. The ascending trend was less apparent at disparate than identical landmarks, reflecting the larger difference in relative activity levels of cue cells between disparate landmarks.

Thus, population code of cue cells distinguishes identical and disparate landmarks (Fig. 3F).

### Landmark identity is modulated by spatial shifts of cue cells

The spatial shift of cue cell activity potentially provides vectorial information about an animal’s spatial location relative to landmarks^19^. To understand how this spatial encoding interacts with landmark identity encoding, we investigated the identity encoding by cue cells at various spatial shifts (Fig. 4A). Spatial shifts of cue cells were largely limited to 50 cm before and after landmarks (Fig. 4B), as previously reported^19^. Besides a small fraction of cue cells active at landmarks (“at-cue cells” with zero spatial shifts), most cue cells were active at certain distances away from landmarks and there were more cells active before than after landmarks (“before-cue” and “after-cue cells” with positive and negative spatial shifts, respectively) (Fig. 4C).

**Figure 4.**
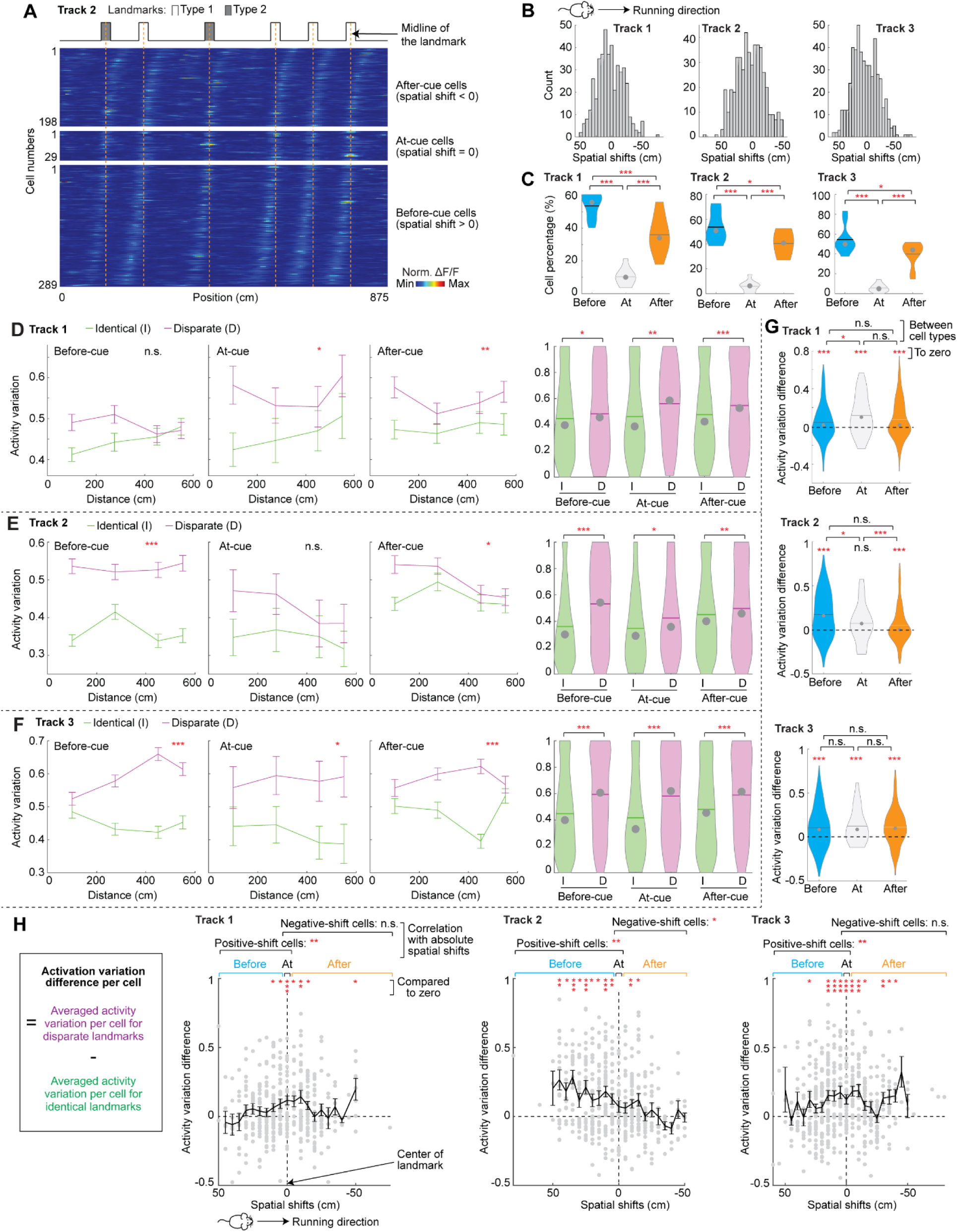
Landmark identity encoding of cue cells with various spatial shifts. **A.** Activity of before-cue, after-cue, and at-cue cells in track 2. The activity of each cell is normalized by its mean and all activities are sorted by their spatial shifts. The red vertical lines indicate the centers of individual landmarks. **B.** Spatial shifts of cue cells in each track. **C.** The percentages of before-cue (Before), at-cue cells (At), and after-cue (After) in tracks 1-3. **D.** Track 1: activity variation of before-cue, at-cue, and after-cue cells at individual and combined distances (the last panel). **E** and **F.** Similar to D but for tracks 2 and 3. **G.** Activity variation differences of before-cue, at-cue, and after-cue cells compared with zero and between these cell types. **H.** For each track: activity variation differences of individual cue cells (gray dots) as a function of their spatial shifts. The dashed horizontal line indicates zero difference. The black curve represents the mean and standard deviations of the difference at spatial shifts with at least 3 cells. The asterisks next to “positive-shift” or “negative-shift” cells indicate a significant correlation between the differences and the absolute spatial shifts.

We first asked whether landmark identity encoding was regulated by the direction of cue cell activity (i.e., before, at, or after landmarks). In all three tracks, before-cue, at-cue, and after-cue cells exhibited higher activity variations between disparate than identical landmarks (Fig. 4D-F) and activity variation differences above zero (Fig. 4G, “to zero”), indicating the general encoding of landmark identity regardless of cue cell activity direction. However, the relative levels of activity variation differences among these cell types varied across different tracks (Fig. 4G, “between cell types”), suggesting a track-specific modulation of landmark identity encoding by the activity direction.

We next examined how landmark identity encoding changes with the distance between cue cell activity and landmarks (i.e., the absolute spatial shifts). We calculated the activity variation difference of each cell by subtracting its averaged activity variation for identical landmarks from that of disparate ones and plotted the differences of individual cells as a function of their spatial shift (Fig. 4H). For each track, we further grouped cue cells into “positive-shift cells” (at-cue and before-cue cells) and “negative-shift cells” (at-cue and after-cue cells), and separately calculated the correlation between their activity variation differences and absolute spatial shifts. We observed significant correlations for positive-shift cells in all tracks, but the trends of correlation varied: the cells in tracks 1 and 3 showed increased activity variation differences with reduced spatial shifts, whereas the cells in track 2 showed the opposite trend. In contrast, significant correlations were not always observed in negative-shift cells. The difference in the correlation significance for positive-shift and negative-shift cells was not due to the fewer number of negative-shift cells (Fig. S3A). Additionally, the track-specific relationship between activity variation difference and spatial shift was also true for individual mice (Fig. S3B). Thus, while landmark identity is modulated by the distance between cue cell activity and landmarks, more consistent modulation occurs when a mouse is approaching (when positive-shift cells are active) than moving away from landmarks (when negative-shift cells are active). The trend of the modulation is track-specific.

Collectively, these results reveal both general and track-specific interactions between landmark identity encoding and spatial shifts of cue cells. Landmark identity is encoded by cue cells active in all directions (before, at, and after landmarks). The encoding was more consistently modulated by the distance of cue cell activity from landmarks when a mouse is approaching landmarks. However, the detailed modulations by direction and distance are track-specific.

### The encoding of landmark identity by cue cells changes in different environments

Give the above track-specific modulation of landmark identity encoding by spatial shifts of cue cells, we further asked whether landmark identity encoding by the same cue cell population changes across different tracks. We focused on common cue cells in each pair of tracks (tracks 1 and 2, 2 and 3, or 1 and 3) (Fig. 5A). These cells showed weakly correlated spatial shifts across tracks (in two out of three track pairs) (Fig. 5A and B), as previously reported^19^, but exhibited uncorrelated cue scores (Fig. 5C).

**Figure 5.**
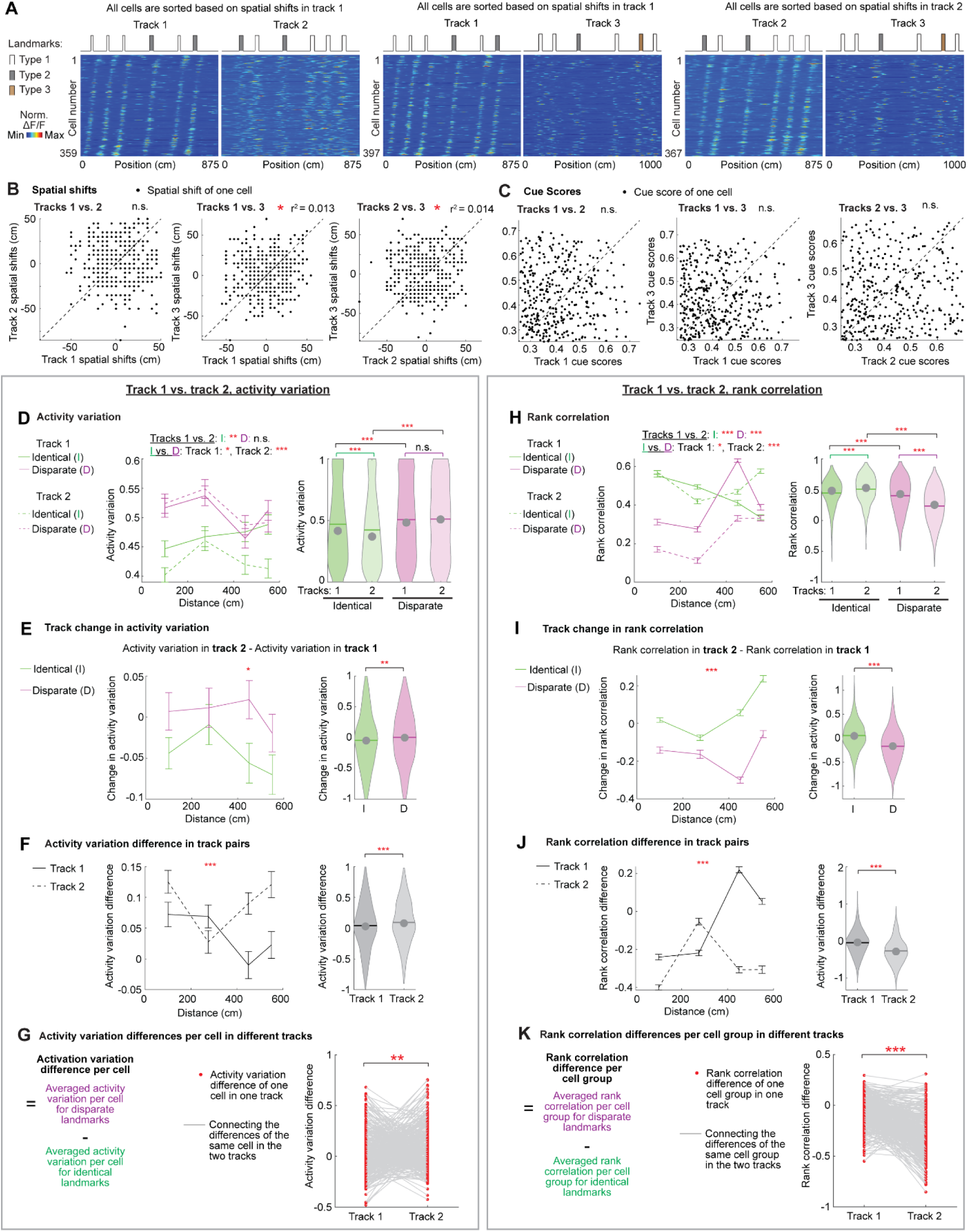
Landmark identity encoding by cue cells in different tracks. **A.** Common cue cells on individual track pairs. For tracks 1 and 2, the activity of each cell is normalized by its mean and all activities were sorted by their spatial shifts to landmark templates in track 1. Same for other track pairs. **B.** The correlation between spatial shifts of individual common cue cells in different tracks. The asterisks indicate a significant correlation. R^2^ represents the fit of the data to the diagonal line (dashed line), which indicates equal shifts in the two tracks. Same in Figure 6. **C.** Similar to B but for cue scores. **D.** Activity variation of common cue cells in tracks 1 (solid line) versus 2 (dashed line) for identical (I, green) and disparate landmark pairs (D, magenta). The statistics on the left represent the comparison of two curves using two-day ANOVA: “Tracks 1 vs 2 I”: identical landmarks on tracks 1 versus 2. “Tracks 1 vs 2 D”: disparate landmarks on tracks 1 versus 2. “I vs D track 1”: identical versus disparate landmarks in track 1. “I vs D track 2” identical versus disparate landmarks in track 2. The statistics on the right represent the results of two-tailed t-test. **E.** “Track changes” in activity variation for identical (green) and disparate landmark pairs (magenta) in tracks 1 and 2. **F.** The comparison of activity variation difference in tracks 1 and 2. **G.** Activity variation differences of individual common cells in tracks 1 and 2. **H-K.** Similar to D-G but for rank correlations in tracks 1 and 2. Each cue cell group contained 15 cells.

We first compared the activity variation of common cue cells. Across tracks 1 and 2, the cells changed their activity variations at identical landmarks (Fig. 5D, the two green groups), but not at disparate ones (Fig. 5D, the two magenta groups). The different degrees of change for the two landmark types (Fig. 5E) led to different activity variation differences between the two tracks (Fig. 5F). Activity variation differences of individual cue cells also significantly altered across the two tracks (Fig. 5G). The same results were also obtained for other track pairs (Fig. S4).

Similarly, cue cell groups changed their rank correlations at identical and disparate landmarks in different tracks (Fig. 5H and S5A). The different degrees of change for the two landmark types (Fig. 5I and S5B) resulted in altered rank correlation differences across each track pair (Fig. 5J and S5C). Rank correlation differences of individual cell groups also changed between tracks (Fig. 5K and S5D) and the change was largely consistent for the groups with various numbers of cells (Fig. S5E).

In summary, the ability of cue cells to encode landmark identity varies in different environments.

### The encoding of landmark location but not identity is modulated by experience

We next investigated whether landmark identity encoding by cue cells is affected by increased experience in the same environment during spatial learning. We focused on tracks 1 and 2, which allowed the comparison of cue cell activity on day 1 (with less experience) and day 7 (with more experience) in the same track. Common cue cells on the two days exhibited well-correlated spatial shifts (Fig. 6A and B) and cue scores (Fig. 6C), in contrast with the weakly correlated spatial shifts (Fig. 5A and B) and uncorrelated cue scores on different tracks (Fig. 5C). Thus, cue cells better preserve their landmark-associated activation within the same track than between different tracks.

**Figure 6.**
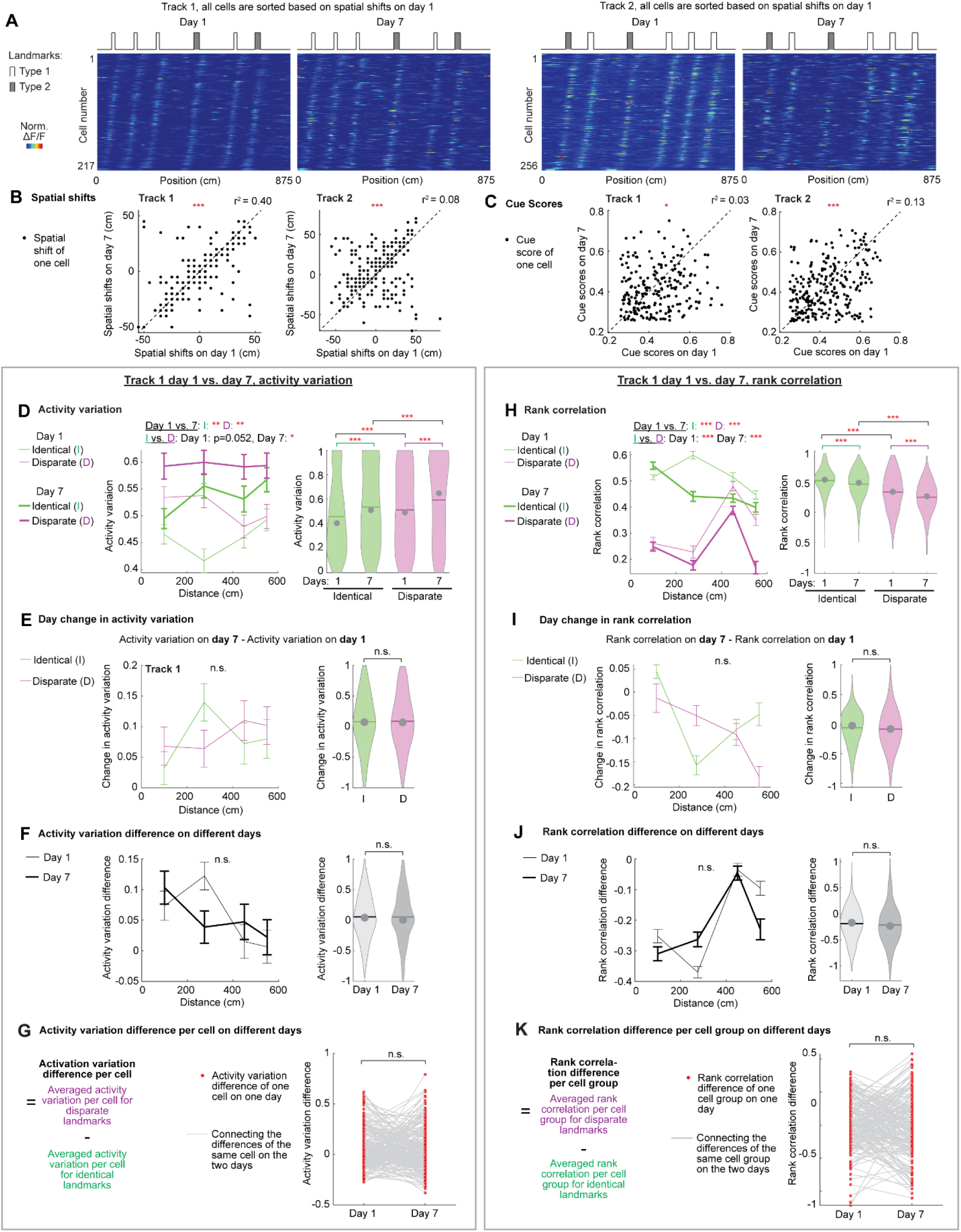
Landmark identity encoding by cue cells on different days in the same track. **A.** Activity of common cue cells on days 1 and 7 in tracks 1 (left) and 2 (right). The activity of each cell is normalized by its mean and all activities were sorted by their spatial shifts on day 1. **B.** The correlation between spatial shifts of individual common cue cells on days 1 and 7 in tracks 1 and 2. **C.** Similar to B but for cue scores. **D.** Activity variation on days 1 (thin line) and 7 (thick line) for identical (green) and disparate landmark pairs (magenta) in tracks 1. The statistics on the left represent the comparison of two curves using two-way ANOVA: “Days 1 vs 7 I”: identical landmarks on days 1 versus 2. “Days 1 vs 7 D”: disparate landmarks on days 1 versus 2. “I vs D day 1”: identical versus disparate landmarks on day 1. “I vs D day 7”: identical versus disparate landmarks on day 7. The statistics on the right represent the results of two-tailed t-test. **E.** Learning changes between days 1 and 7 in activity variation for identical and disparate landmark pairs in track 1. **F.** The comparison of activity variation difference on days 1 and 7 in track 1. **G.** Comparison of activity variation differences of individual common cells on days 1 and 7 of track 1. **H-K.** Similar to D-G but for rank correlation on days 1 and 7 in track 1.

We observed that activity variation of common cue cells in track 1 increased from days 1 to 7 for both identical and disparate landmarks (Fig. 6D), suggesting an improved ability to differentiate landmarks after increased experience. The increase was consistently observed in before-cue cells but not in after-cue cells, indicating a specific improvement in prospective landmark encoding (Fig. S6A). Interestingly, the degree of improvement was similar for identical and disparate landmarks (Fig. 6E and S6B), and consequently, activity variation differences on the two days were comparable (Fig. 6F and S6C). Moreover, the activity variation differences of individual cells on the two days also showed no overall change (Fig. 6G). The same results are also obtained for track 2 (Fig. S6 and S7A-D).

Similarly, cue cell groups (Fig. 6H and Fig. S7E), especially the before-cue cell groups (Fig. S8A), decreased their rank correlation from days 1 to 7 by the same extent for identical and disparate landmarks (Fig. 6I, S7F, and S8B), leading to similar rank correlation differences on the two days (Fig. 6J, Fig. S7G and S8C). For individual cue cell groups, their overall rank correlation differences on the two days were comparable (Fig. 6K and S7H) regardless of the number of cells per group (Fig. S7I).

Overall, with increased environmental experience, cue cells maintain the same level of landmark identity encoding, contrasting with the changed encoding in different tracks (Fig. 5, S4 and S5). Therefore, landmark identity encoding by cue cells is environment-specific. Notably, increased experience leads to improved ability of cue cells to distinguish landmarks regardless of their identity and distance, suggesting a better encoding of landmark location. Thus, the encodings of landmark identity and location by cue cells are differentially regulated by experience, suggesting different circuit mechanism underlying these encodings.

### The encodings of landmark identity and location are differentially supported by cue cell and grid cells

While the above results revealed the integration of spatial and nonspatial information in cue cell activity, we further asked whether this integration also occurs to grid cells, which are widely believed to encode spatial information during navigation^10^. We classified grid cells on days 1 and 7 in tracks 1 and 2^22,23,25–27^. The grid cells conjunctive with cue cells^19^ were removed to minimize the effect of cue-cell-like activity on grid cell response. Indeed, the remaining grid cells did not show cue-specific activity (Fig. 7A).

**Figure 7.**
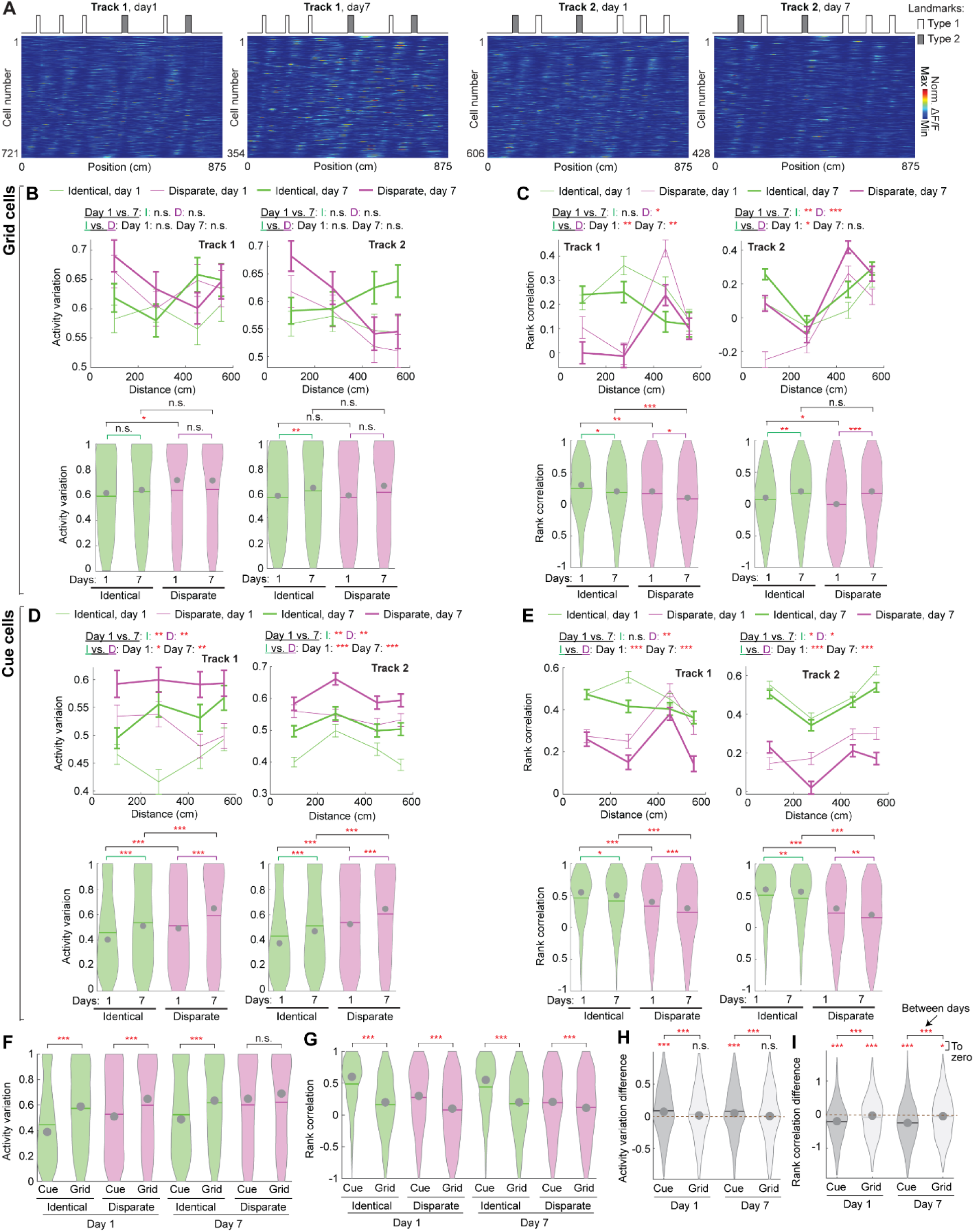
Landmark identity encoding by spatial activity of grid and cue cells. **A.** Grid cells on days 1 and 7 in tracks 1 and 2. The activity of each cell is normalized by its mean and all activities were sorted by their spatial shifts to landmark templates. **B.** Activity variation of grid cells for identical (green) and disparate landmarks (magenta) on days 1 (thin lines) and 7 (thick lines). **C.** Similar to B but for rank correlations. Each cue cell group contained 5 cells due to the small number of common grid cells on days 1 and 7 in each FOV. **D and E**. Similar to B and C but for cue cells. The panels in D are identical to those in Fig. 6D and Fig. S7A and are shown here just for the direct comparison with those for grid cells. The panels in E are different from those in Fig. 6H and Fig. S7E, which used 15 cue cells per group. All results in E used 5 cells per group to match the calculations of grid cells, which had a lower number of common cells per FOV. **F and G.** The comparison between activity variation (F) and rank correlation (G) of grid and cue cells. **H and I.** The comparison between activity variation difference (H) and rank correlation difference (I) of grid and cue cells.

We first evaluated activity variation and rank correlation of spatial activity of common grid cells on days 1 and 7 in tracks 1 or 2. Like cue cells, their “landmark zones” were identified after shifting their activity patterns to best match the landmark template. Grid cells showed largely comparable activity variation for disparate and identical landmarks (Fig. 7B) and a slight trend of lower rank correlation for disparate ones (Fig. 7C). These activity parameters were not consistently regulated by landmark distance or experience. In comparison to cue cells (Fig. 7D and E), grid cells showed greater activity variation (Fig. 7F) and lower rank correlation (Fig. 7G) between both identical and disparate landmarks, reflecting their better ability to differentiate landmarks, regardless of their identity. Since these parameters were not consistently regulated by landmark distance, they likely reflect a stronger encoding of landmark location by grid cells than cue cells. However, the differences in these parameters between identical and disparate landmarks were smaller in grid cells (Fig. 7H and I), suggesting their weaker encoding of landmark identity. These conclusions also held when comparing spatial activity of grid and cue cells on day 1 in track 3 (Fig. S9A-H). Overall, spatial activity of grid cells exhibits stronger encoding of landmark location but weaker encoding of landmark identity than cue cells.

However, since the above analyses were designed for landmark-specific activity of cue cells, they may not properly reveal landmark identity encoding by grid cells. Therefore, we investigated whether landmark identity can be decoded from temporal activity of grid cells (Fig. 8A). The decoding was based on a population of 40 simultaneously imaged grid cells randomly drawn from a FOV. For a landmark pair, we generated population activity vectors of the cells in even runs across individual spatial bins of the zone around each landmark. We then asked whether, in odd runs, the activity at individual time points in the zone of a landmark better matched the population vectors in the zone of the same or different landmark, corresponding to correct and incorrect landmark decoding, respectively. The percentage of correct decoding combining all temporal activity around the two landmarks was defined as “decoding accuracy” (Fig. 8A, a). If grid cell activity decodes landmark identity, the decoding accuracy should be higher at disparate than identical landmarks, leading to positive “decoding accuracy difference”, which was the accuracy at disparate landmarks subtracted by that at identical ones (Fig. 8A, b).

**Figure 8.**
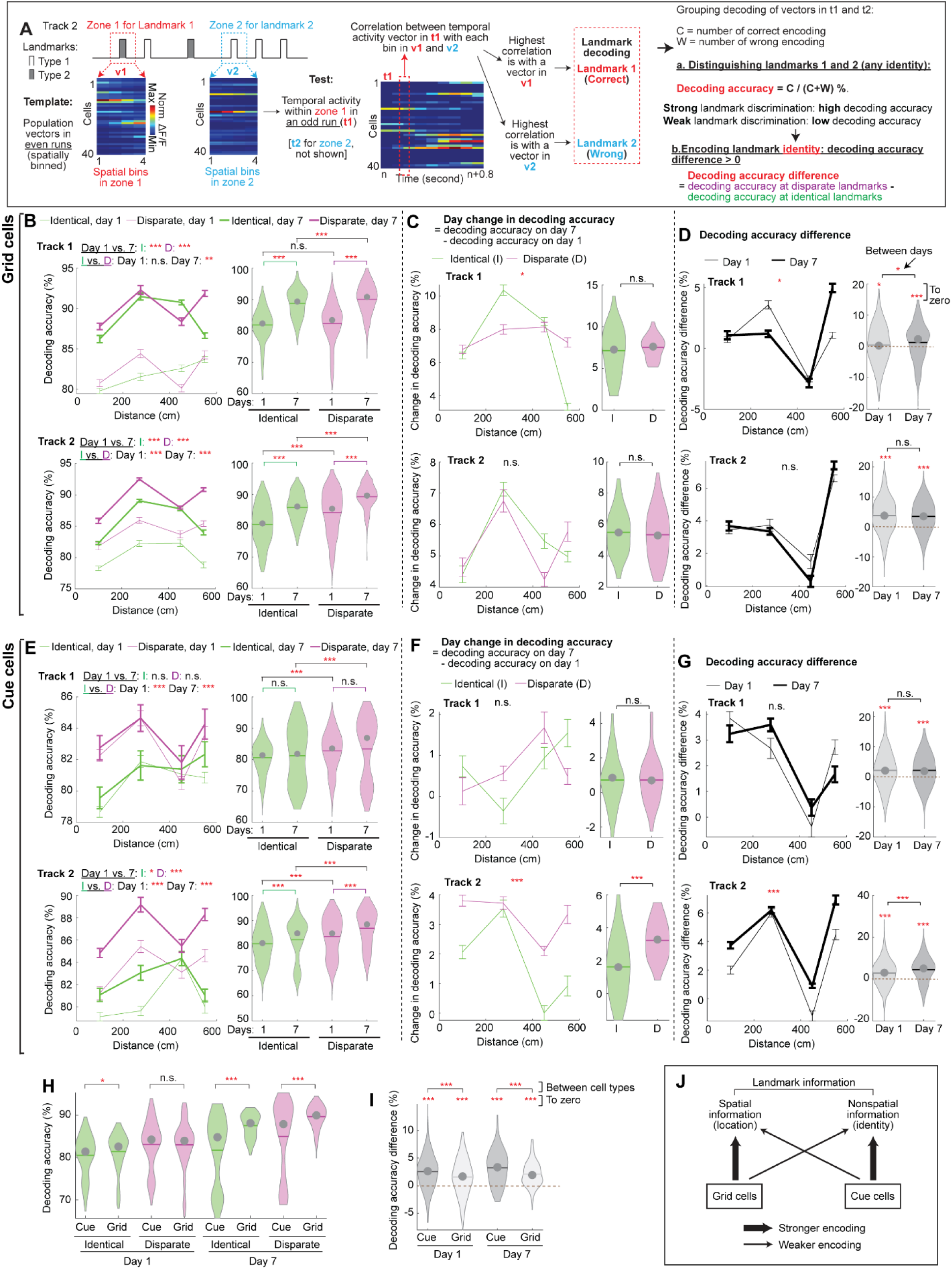
Encoding of landmark identity by temporal activity of grid and cue cells. **A.** The calculation of landmark decoding. The zone of each landmark was determined as 50 cm before and after the center of the landmark. **B.** Grid cells: the accuracy of landmark decoding before (thin lines) and after learning (thick lines) for identical (green) and disparate landmark pairs (magenta) in tracks 1 and 2. **C.** Grid cells: the comparisons between the learning change in decoding accuracy for identical (green) and disparate landmark pairs (magenta). **D.** Grid cells: the comparisons between the decoding accuracy difference on days 1 (thin lines) and 7 (thick lines). **E-G.** Similar to B-D but for cue cells. **H.** The comparison between decoding accuracy of grid and cue cells. **I.** The comparison between decoding accuracy differences of grid and cue cells. **J.** A summary of the encoding of landmark location and identity by grid and cue cells. Thick and thin lines indicate strong and weak encodings, respectively.

In tracks 1 and 2, grid cells showed a high decoding accuracy for identical landmarks regardless of their distances (Fig. 8B, >80%), suggesting that the activity represents landmark location. Additionally, there was a trend of higher accuracy in decoding disparate than identical landmarks, revealing grid cell’s ability to discriminate landmark identity. The decoding accuracy for identical and disparate landmark pairs similarly improved from day 1 to day 7 (Fig. 8C), and therefore, there was no consistent regulation of decoding accuracy difference by track experience (Fig. 8D). Thus, increased experience leads to improved representation of landmark locations but not identity by temporal activity of grid cells.

On the other hand, while the efficient representations of landmark location and identity were also observed in temporal activity of cue cells (Fig. 8E), these representations were not consistently regulated by experience in tracks 1 and 2 (Fig. 8E-G).

Comparing the decoding results between grid and cue cells, we found that similar to spatial activity (Fig. 7F-I), temporal activity of grid cells showed higher decoding accuracy for both identical and disparate landmark pairs, especially after learning (Fig. 8H), indicating their stronger ability to encode landmark location. In contrast, cue cells showed higher decoding accuracy difference between these landmark pairs (Fig. 8I), implying their more robust representation of landmark identity. These conclusions were also confirmed by the temporal activity of grid and cue cells on day 1 in track 3 (Fig. S9I-L).

In summary, while grid cells and cue cells represent both landmark location and identity, grid cells preferentially represent landmark location, whereas cue cells better represent landmark identity. Therefore, the representations of landmark location and identity are preferentially supported by different functional cell types (Fig. 8J).

## Discussion

Here, we demonstrate a novel function of the MEC to encode landmark identity, a function that is previously thought to be conducted by the LEC, the hippocampus, and other association cortices^28–33^. This ability is supported by spatial activity of cue cells, which respond to individual landmarks during virtual navigation^19^. Cue cells encoded landmark identity by responding more differentially to disparate than identical landmarks. The encoding was conducted by cue cells active before, at, and after landmarks, but the relative encoding levels were modulated by the activity locations of individual cue cells in a track-specific manner. The same cue cell population altered their landmark identity encoding in different tracks but maintained the encoding in the same environment, despite increased environmental experience. In contrast, increased experience led to improved prospective encoding of landmark locations by cue cells. Finally, grid cell activity also represented landmark identity, but to a much lesser extent than cue cells. However, grid cells more robustly encoded landmark locations. These results highlight the MEC’s capacity to encode landmark identity so that spatial and nonspatial information of an environment can be integrated in the MEC cognitive map before being delivered to the hippocampus. Furthermore, the encodings of landmark location and identity are independently regulated by experience and are differentially conducted by cue cells and grid cells, suggesting separate circuit mechanisms for these encodings.

The representation of landmark identity by MEC cue cells exhibits both uniqueness and commonality compared to other brain regions, such as the hippocampus, the LEC, and other association cortices^28–32^. First, object selectivity in the other regions is mostly achieved by the cells specifically active at a particular subset of objects^28–32^. However, cue cells respond to all landmarks^19^ but vary their activity at the individual ones. A similar response was also identified in landmark vector cells in the hippocampal CA1^33^. Second, object-specific activity in the LEC and many association cortices largely appears when animals reach the object location^28–31^, whereas cue cells can be active when the mice are away from individual objects at certain distances and orientations. This feature is also observed in object vector cells in the MEC^18^, landmark vector cells in the hippocampus^32^, boundary vector cells in the subiculum^34^, and some object-responsive cells in the LEC^28^. This allows cue cells to not only provide a vectorial representation of animal’s location relative to landmarks, but also represent landmark identity across a broad area around landmarks, potentially supporting precise mapping of animal’s position during real-time navigation in environments with complex landmark features. Lastly, the encoding of specific objects can be regulated by either sensory stimuli or cognition, as some object-responsive cells were only active when the objects were present (sensory-stimuli-driven)^28,30–32^, whereas others in the LEC, the hippocampus, and the anterior cingulate cortex encode the previous existence of objects at specific places (cognition-driven)^28,30,32,35^. In terms of cue cells, a previous study found that their landmark response disappeared when the landmark was removed^19^, indicating that the response does not encode the previous existence of landmarks. Here, we revealed that the dependency of landmark identity encoding on cue cell activity location relative to landmarks was environment-specific. The identity encoding by the same cue cell population was also environment-specific and was not regulated by increased experience.

These results suggest that landmark identity encoding by cue cells is driven by sensory features of an environment.

In addition to cue cells, we also discovered the encoding of landmark identity by grid cells. This finding is surprising because grid cells are widely believed to primarily encode spatial information during navigation^10^. Previous studies also found that grid cells exhibited object-selective activity when the objects were associated with reward^17^ and grid cell activity remapped in environments with comparable spatial information delivered by cues in different sensory modalities^16^. These results suggest that grid cells not only present spatial information, but also incorporate nonspatial information of an environment.

Furthermore, our study implies that landmark identity and location are both encoded in the MEC but through different mechanisms. First, with increased experience in the same environment, cue cells maintained the same landmark identity encoding, but had improved encoding of landmark location, suggesting that the encoding of landmark identity and location by cue cells is driven by sensory stimuli and cognition, respectively. The separate regulations of the encoding of landmark identity and location by experience were also observed in grid cells. Second, although both grid and cue cells encoded landmark identity and locations, cue cells exhibited stronger encoding of landmark identity, whereas grid cells more robustly encoded landmark location. This cell-type-specific encoding of landmark identity and location further suggests different circuit mechanisms supporting the two types of encoding.

Our study indicates that the MEC encodes both spatial and nonspatial information during navigation. The two types of information are also jointly represented by the LEC based on several anatomical^36^, behavioral^37–39^, and neural recording studies^17,28,29^. On the other hand, previous literature has demonstrated the general activities of the MEC and the LEC for spatial and nonspatial information, respectively^10–12,28,29,40^. Collectively, we propose that both the MEC and the LEC integrate spatial and nonspatial information before delivering it to the hippocampus. Meanwhile, each region also has its own preference towards different types of information. This configuration allows not only a strong output of the full cognitive map from multiple regions (the MEC, the LEC, and the hippocampus), but also dedicated computation of specific information in local circuits (the MEC and the LEC), supporting robust and accurate navigation.

## Resource availability

### Lead contact

Further information and requests for resources and reagents should be directed to the lead contact, Dr. Yi Gu.

### Materials availability

This study did not generate new unique materials.

### Data and code availability

Data and code will be available upon request to the lead contact, Dr. Yi Gu.

## Acknowledgements

We thank all colleagues in the Gu laboratory for supporting the work. We thank Dr. Lorna Role for constructive comments on the manuscript.

This work was supported by the NIH/NINDS Intramural Research Program (to YG).

## Author contributions

Conceptualization: GW, YG

Methodology: GW, JT, TM, YG

Investigation: GW, JT, TM, KC, YG

Visualization: GW, TM, YG

Funding acquisition: YG Supervision: YG

Writing – original draft: YG

Writing – review & editing: GW, JT, TM, KC, YG

---

**Figure S1.**
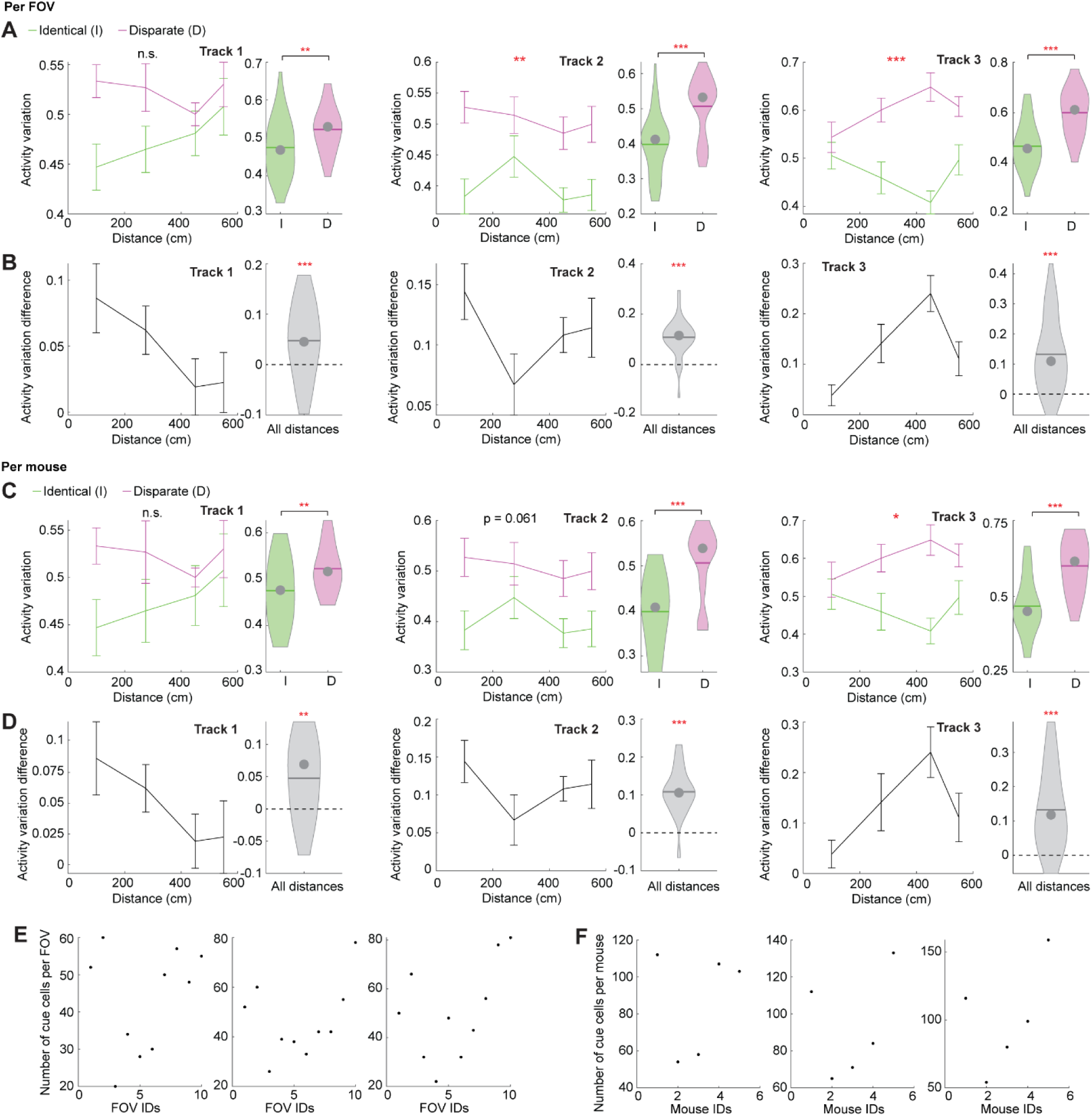
Activity variation per FOV and per mouse. **A.** Per FOV: activity variation for identical (I, green) and disparate landmark pairs (D, magenta) in tracks 1 to 3 (left to right). These data were generated by grouping those in Fig. 2D by FOV. **A.** Per FOV: activity variation difference in the three tracks. **C-D.** Similar to A and B but per mouse. **E.** Number of cue cells per FOV. **F.** Number of cue cells per mouse. *p ≤ 0.05, **p ≤ 0.01, ***p ≤ 0.001, n.s. p > 0.05. Significant p values are in Supplementary Table 1. Error bars represent mean ± SEM. In violin plots, the dot and horizontal bar represent the median and mean, respectively. Same for all figures.

**Figure S2.**
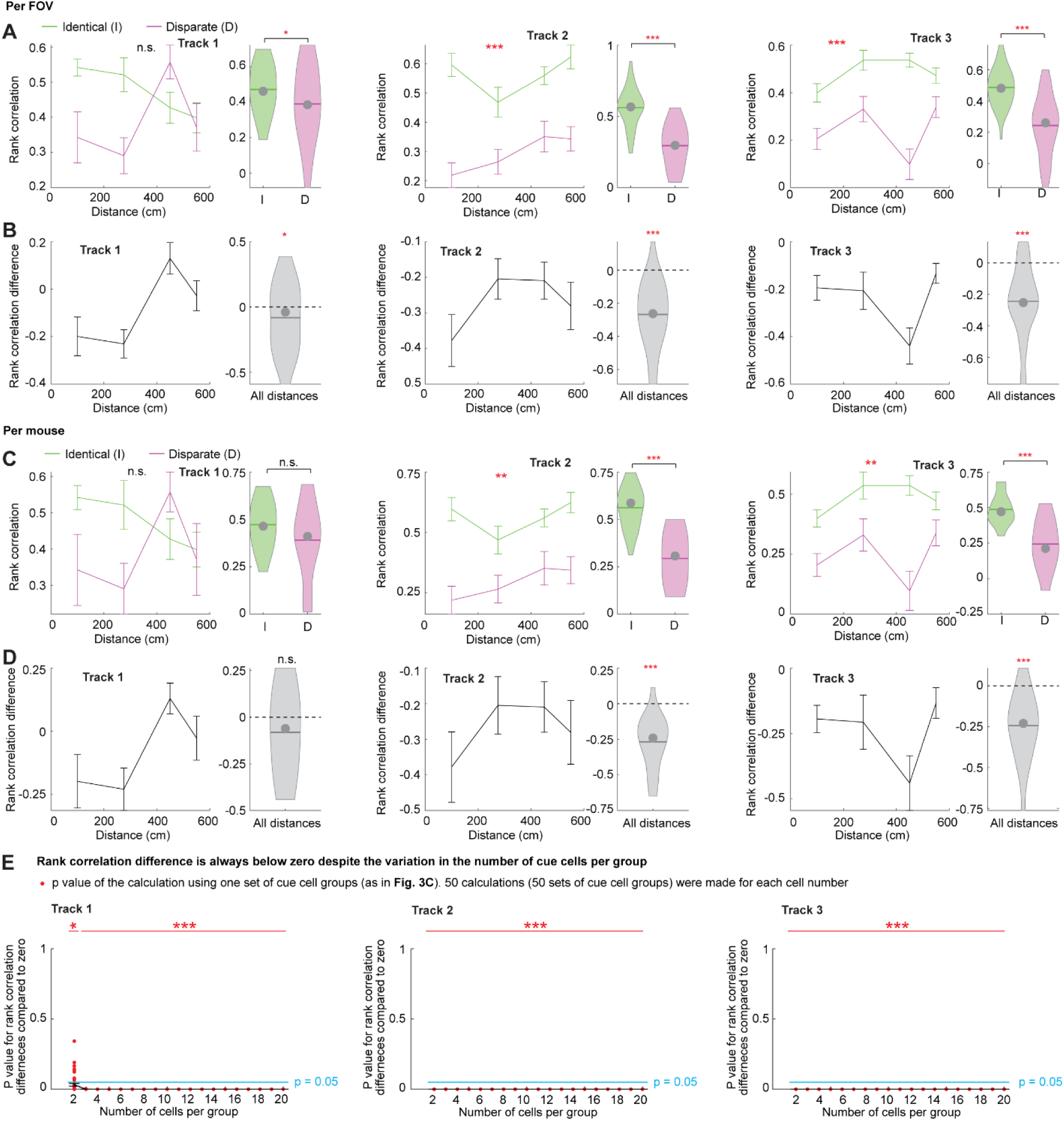
Rank correlation per FOV, per mouse, and with cue cell groups in difference sizes. **A.** Per FOV: rank correlation for identical (I, green) and disparate landmark pairs (D, magenta) in tracks 1 to 3 (left to right). These data were generated by grouping those in Fig. 3B by FOV. **B.** Per FOV: rank correlation difference in the three tracks. **C-D.** Similar to A and B but per mouse. **E.** The comparison between zero and rank correlation differences calculated using cue cell groups of different number of cells (group size). Each red dot indicates the p value for the comparison between zero and rank correlation difference of a set of cue cell groups, as the p value in the right panel of Fig. 3C for each track. Since each calculation was based on cue cell groups randomly chosen from individual FOVs, 50 calculations were made based on 50 sets of cue cell groups for each group size. The individual p values (red dots) and their mean and standard errors (black curve) are plotted and are compared with a significant p value 0.05 (blue line). The asterisks on top indicate the comparison between the 50 p values and 0.05 for each group size.

**Figure S3.**
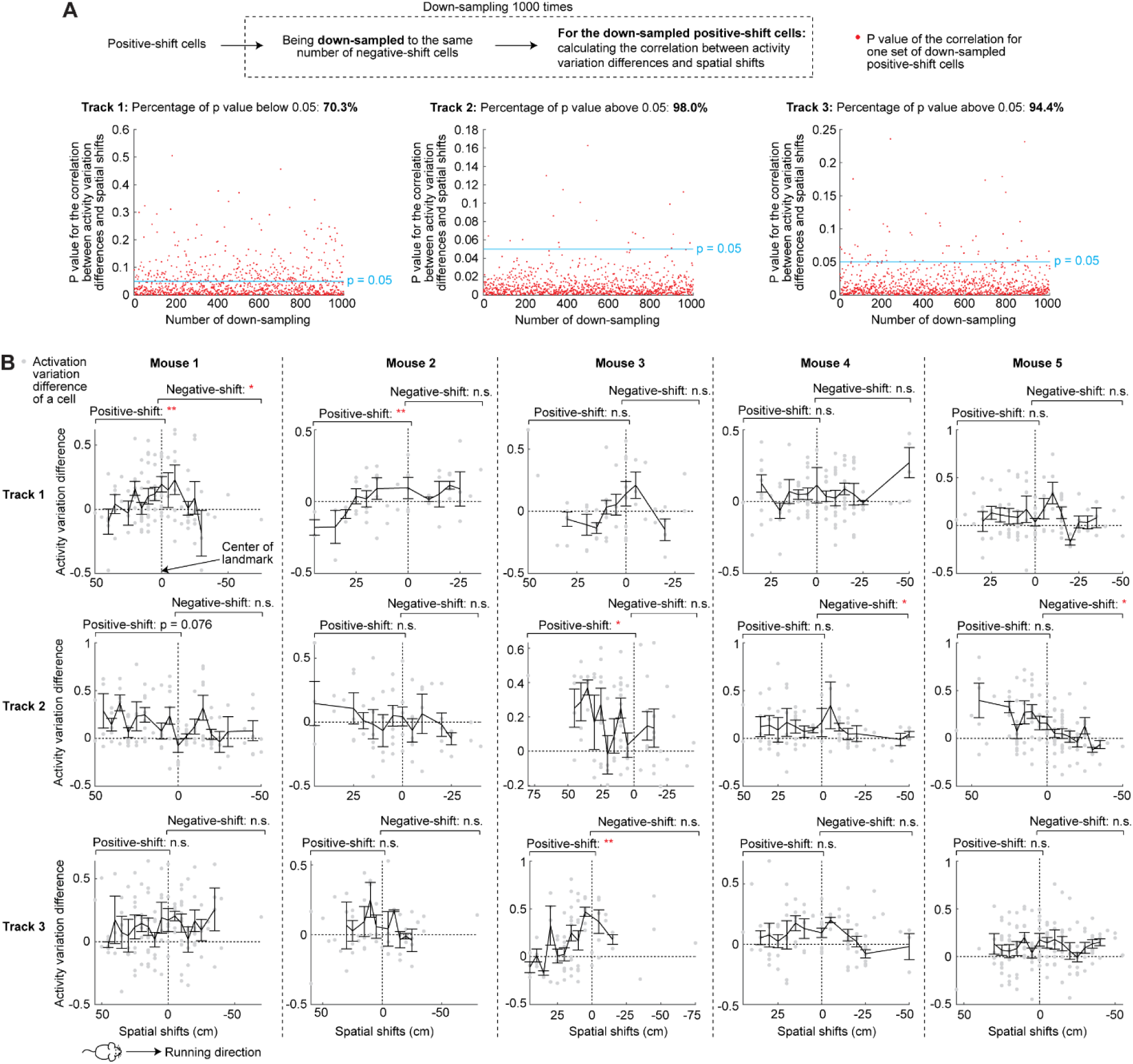
Activity variation difference of cue cells with different spatial shifts. **A.** The significant correlation between activity variation differences and absolute spatial shifts of positive-shift cells is unlikely due to the larger number of positive-shift cells than negative-shift cells. Each red dot represents the p value of the correlation for one calculation, during which positive-shift cells were down-sampled to the same number of negative-shift cells and the correlation between the activity variation differences and absolute spatial shifts of down-sampled positive-shift cells were calculated. 1000 calculations were done for each track and all the p values were compared with the significant p value 0.05 (blue line). The percentage of p values below 0.05 is indicated in the title. For all three tracks, the majority of p values are below 0.05. **B.** For individual mice: activity variation differences of individual cue cells (gray dots) as a function of their spatial shifts.

**Figure S4.**
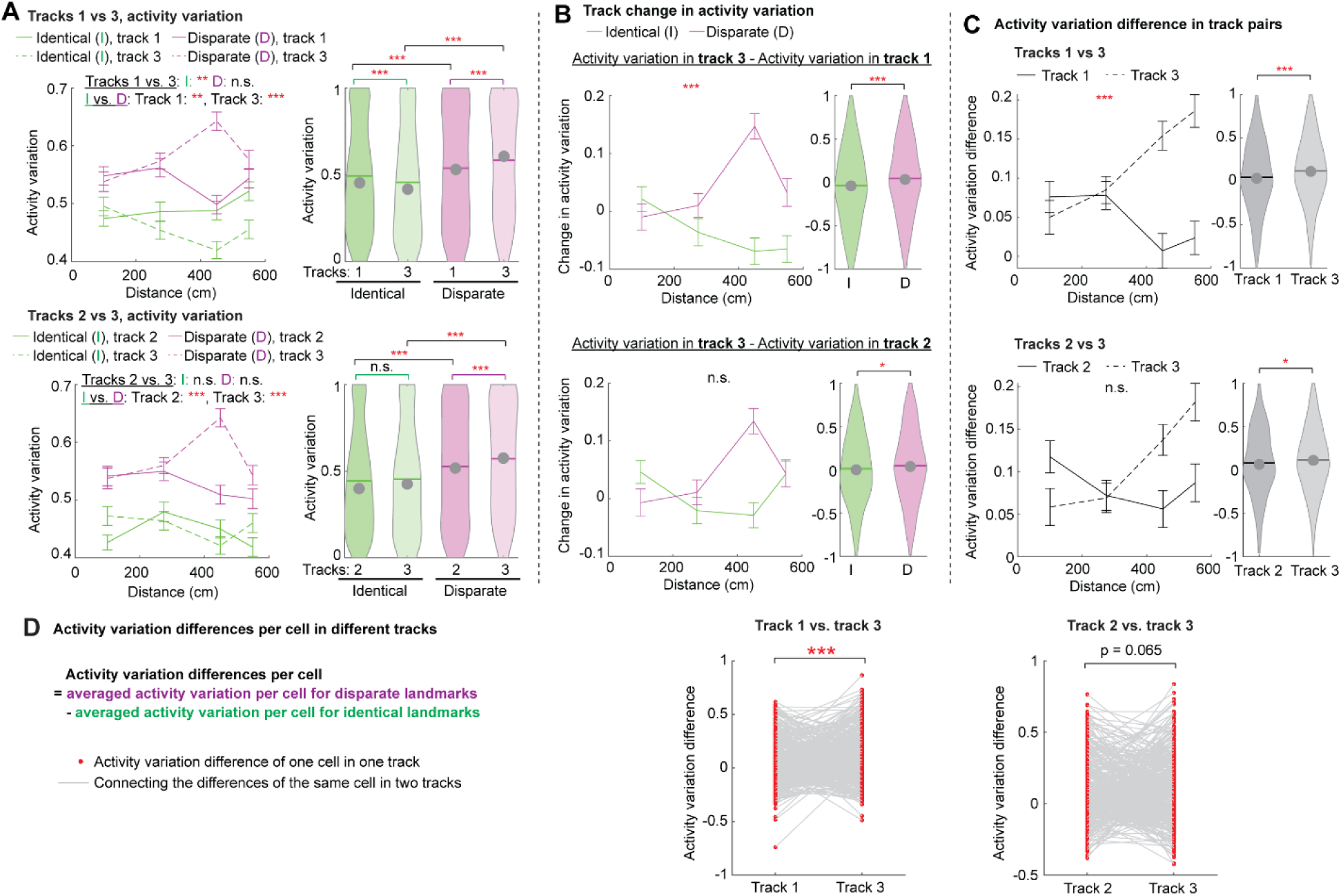
Activity variation of common cue cells in different tracks. **A-C**. Similar to Fig. 5D-F: activity variation, track change in the variation, and activity variation difference for tracks 1 versus 3 (top) and tracks 2 versus 3 (bottom). **D.** Activity variation differences of individual common cells in tracks 1 versus 3 and tracks 2 versus 3.

**Figure S5.**
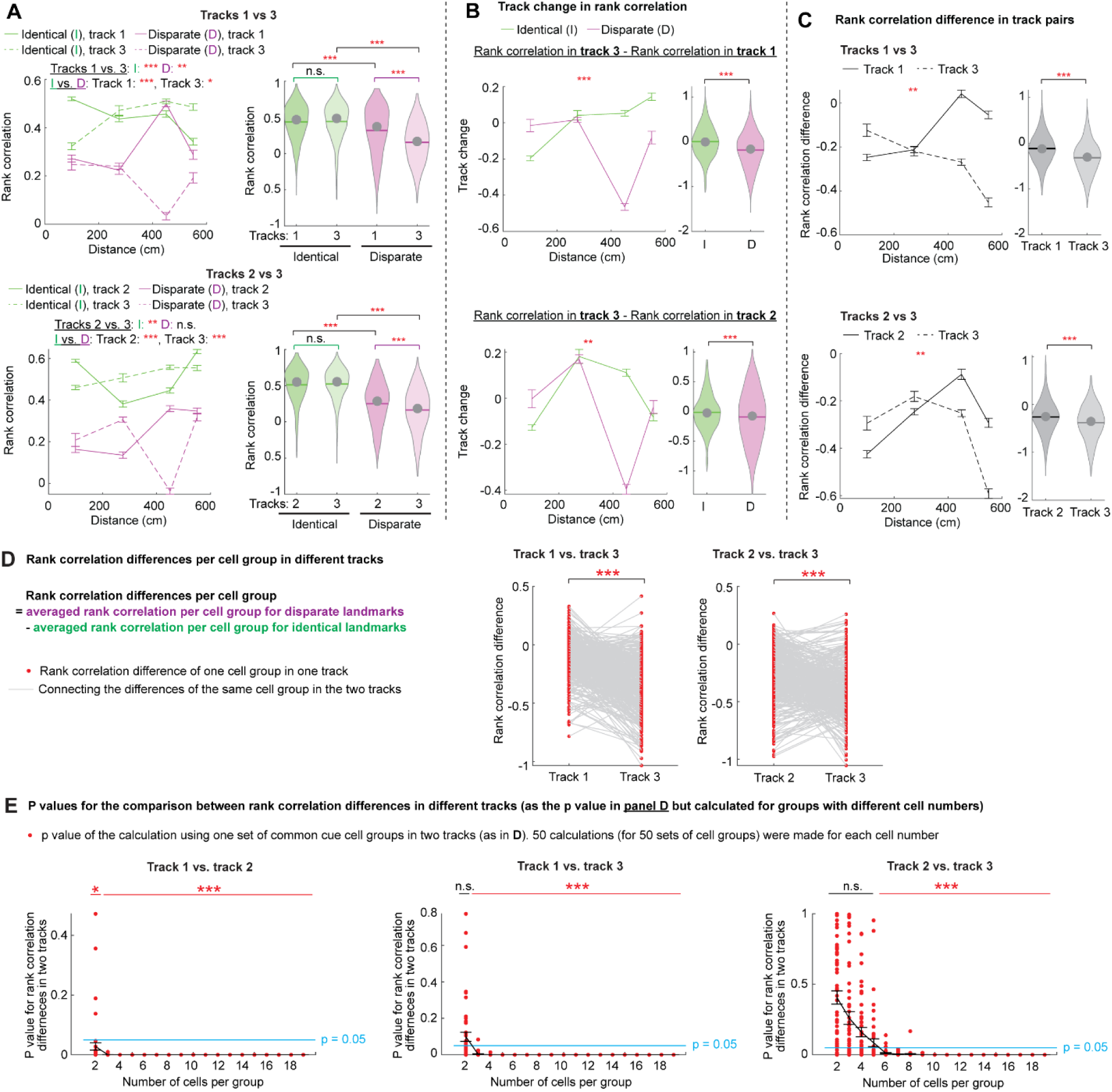
Rank correlation of common cue cell groups in different tracks. **A-D.** Similar to Fig. S4 but for rank correlation of cue cell groups. **E.** P values for the comparison between rank correlation differences of the same group of common cells in different tracks when varying the size of cell groups. Each red dot is the p value for one calculation using one set of common cell groups, as in D. Since the cell groups were randomly sampled within each FOV, 50 calculations were made using 50 sets of common cell groups for each size of cell group. The individual p values (red dots) and their mean and standard errors (black curve) are plotted and are compared with a significant p value 0.05 (blue line). The asterisks or n.s. on top indicate the comparison between the 50 p values and 0.05 for each group size.

**Figure S6.**
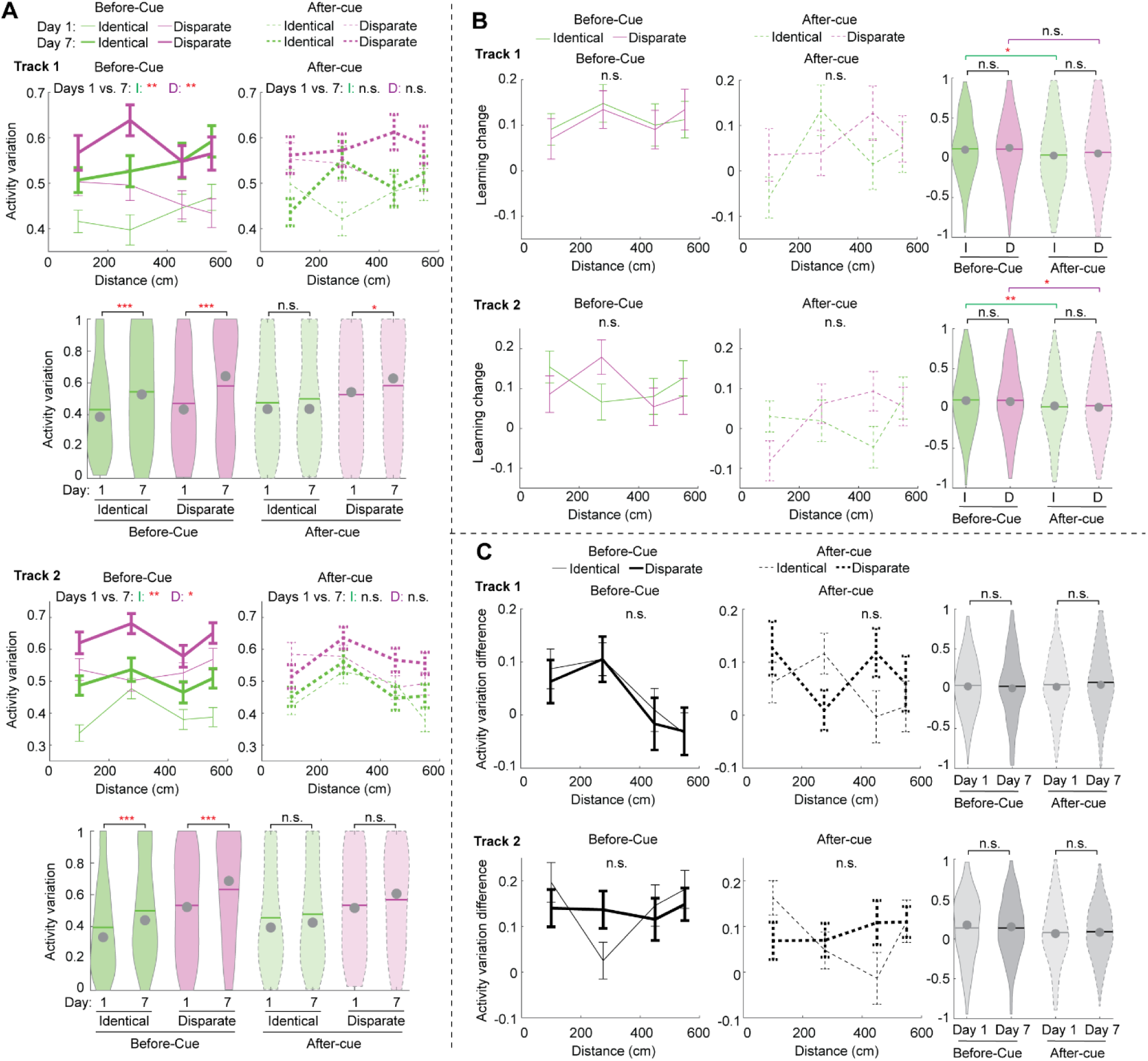
Activity variation of before-cue and after-cue cells during learning. **A.** Activity variation of before-cue (solid lines, left) and after-cue cells (dashed lines, right) for identical (green) and disparate landmark pairs (magenta) on days 1 (thin lines) and 7 (thick lines) in tracks 1 and 2. **B.** The comparison between the learning change in activity variation for before-cue (solid lines, left) and after-cue cells (dashed lines, middle) at identical (green) and disparate landmark pairs (magenta) in tracks 1 and 2. **C.** The comparison between the activity variation difference for before-cue (solid lines, left) and after-cue cells (dashed lines, middle) on days 1 (thin lines) and 7 (thick lines).

**Figure S7.**
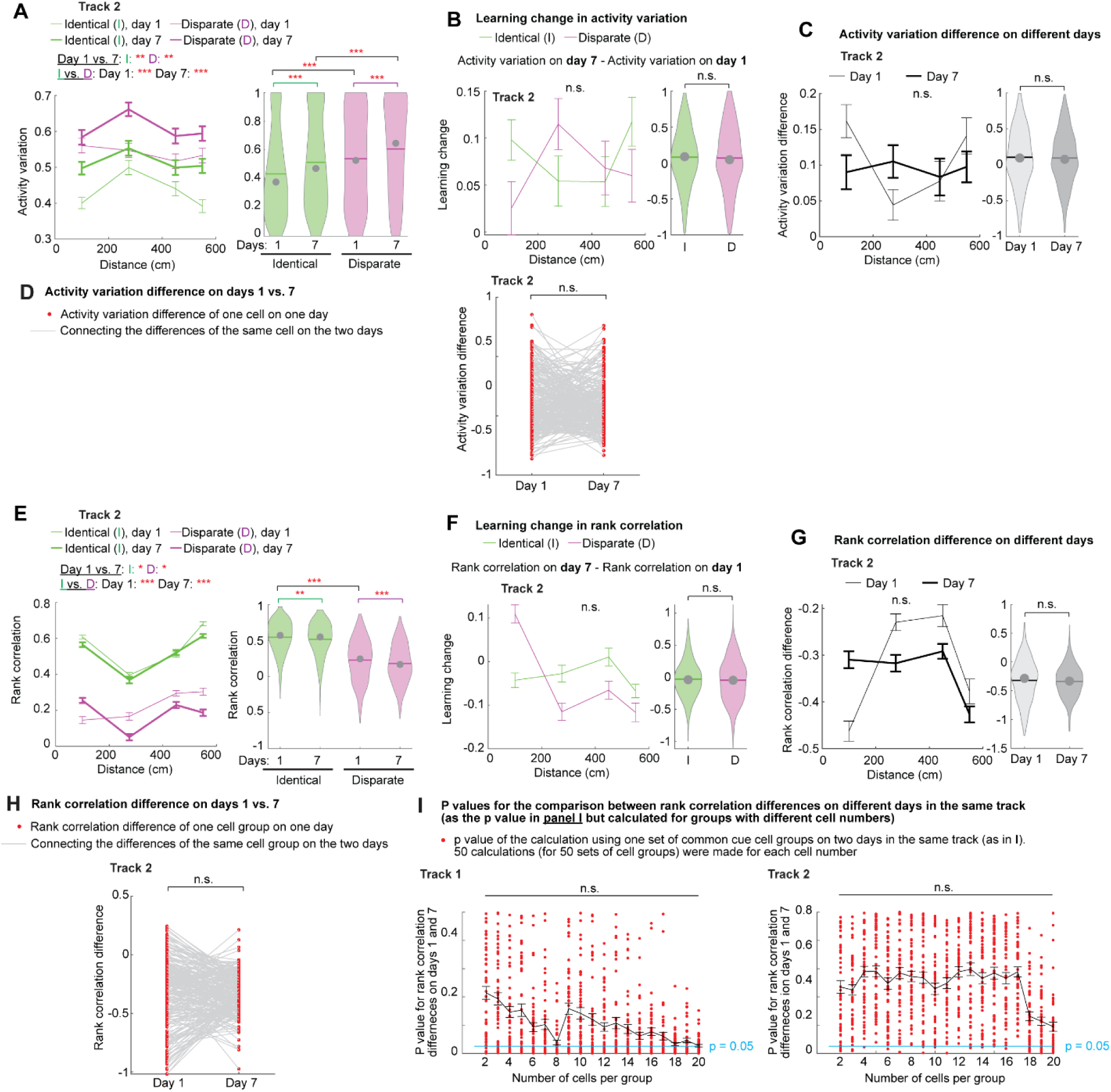
Common cue cells and cell groups on different days in the same tracks. **A-C.** Similar to Fig. 6D-F but for activity variation, the learning change of the variation, and activity variation difference on days 1 and 7 in track 2. **D.** Comparison of activity variation differences of individual common cells on days 1 and 7 of track 2. **E-H.** Similar to A-D but for rank correlation of common cell groups. **I.** P values for the comparison between rank correlation differences of the same group of common cells on days 1 and 7 of the same track when varying the size of cell groups. Each red dot is the p value for one calculation using one set of common cell groups, as in I. Since the cell groups were randomly sampled within each FOV, 50 calculations were made using 50 sets of common cell groups for each size of cell group. The individual p values (red dots) and their mean and standard errors (black curve) are plotted and are compared with a significant p value 0.05 (blue line). The n.s. on top indicate the comparison between the 50 p values and 0.05 for each group size.

**Figure S8.**
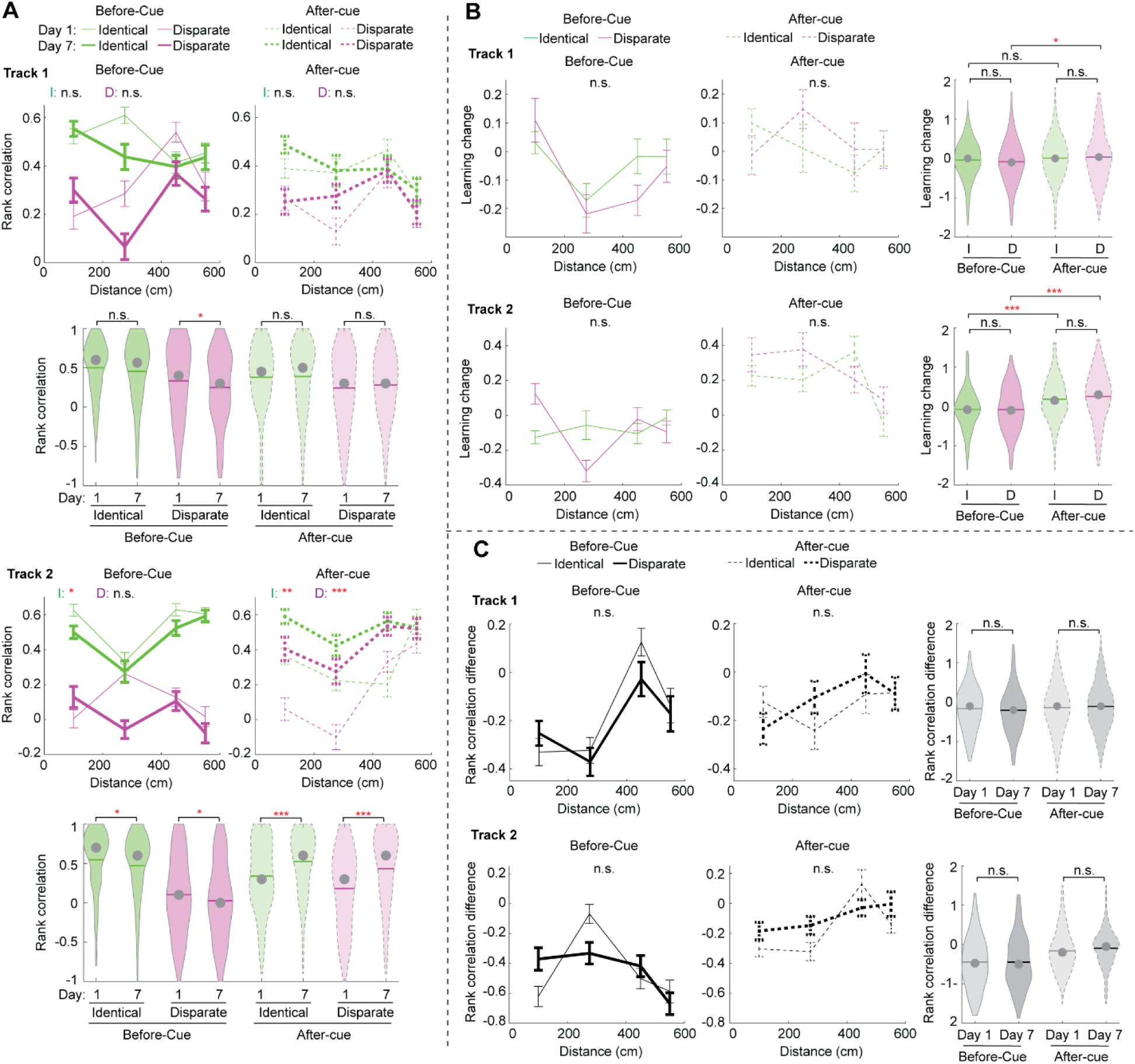
Rank correlation of before-cue and after-cue cells during learning. **A.** Rank correlations of before-cue (solid lines, left) and after-cue cells (dashed lines, right) for identical (green) and disparate landmark pairs (magenta) on days 1 (thin lines) and 7 (thick lines) in tracks 1 and 2. **B.** The comparison between the learning change in rank correlation for before-cue (solid lines, left) and after-cue cells (dashed lines, middle) at identical (green) and disparate landmark pairs (magenta) in tracks 1 and 2. **C.** The comparison between the rank correlation difference for before-cue (solid lines, left) and after-cue cells (dashed lines, middle) on days 1 (thin lines) and 7 (thick lines).

**Figure S9.**
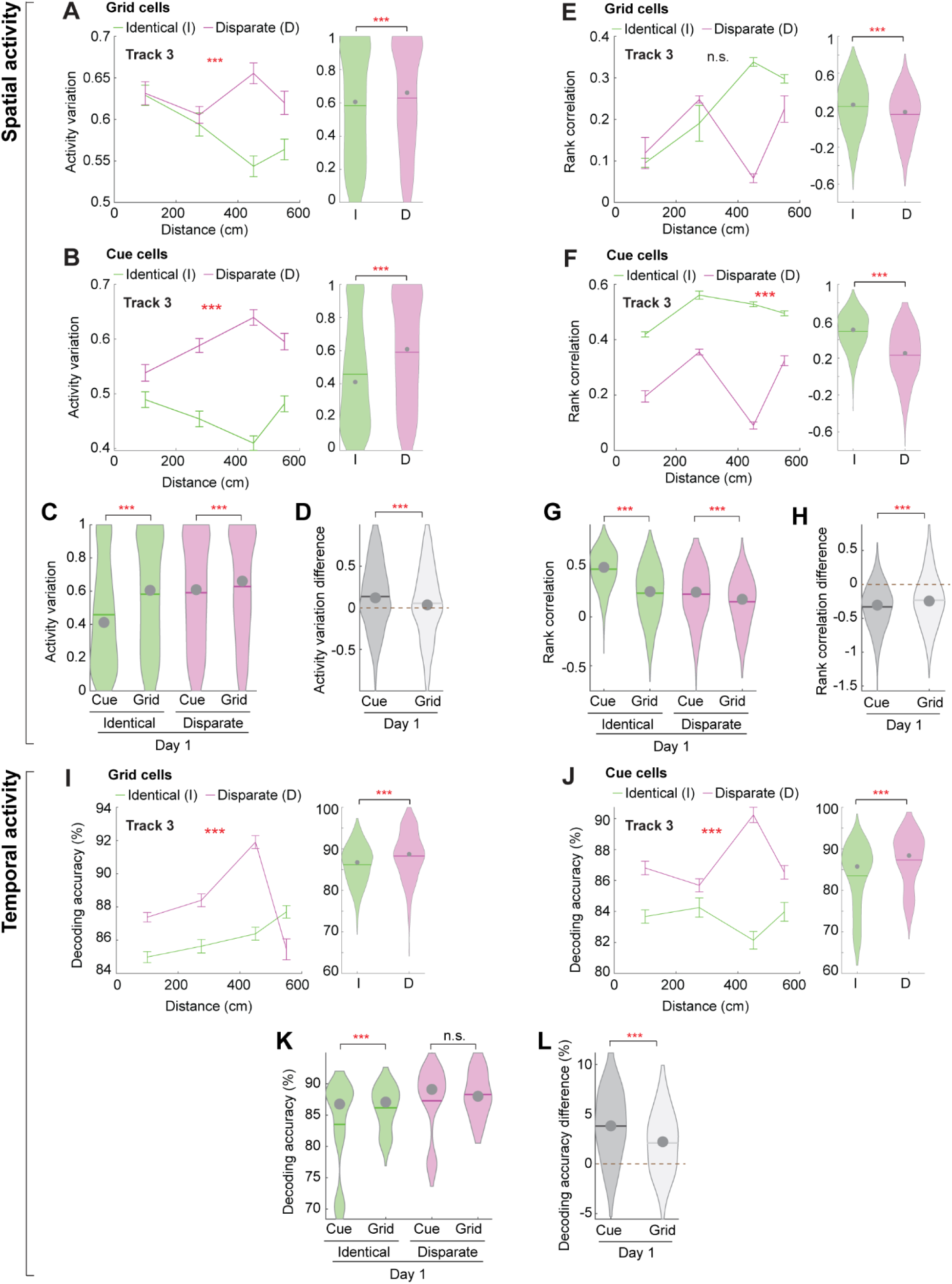
The comparison of grid and cue cells based on their activity on day 1 in track 3. **A.** Activity variation of grid cells for identical (green) and disparate landmarks (magenta). **B.** Similar to A but for cue cells, identical to the plots in Fig. 2D. **C.** Comparison between activity variation of grid and cue cells. **D.** Comparison between activity variation difference of grid and cue cells. **E-H.** Similar to A-D but for rank correlation and its difference. Each cell group contained 15 cells. F is identical to the plots in Fig. 2D. **I.** Decoding accuracy by temporal activity of grid cells between identical (green) and disparate landmarks (magenta). **J.** Similar to I but for cue cells. **K and L.** Similar to C and D but for decoding accuracy and its difference.

**Supplementary Table 1:**
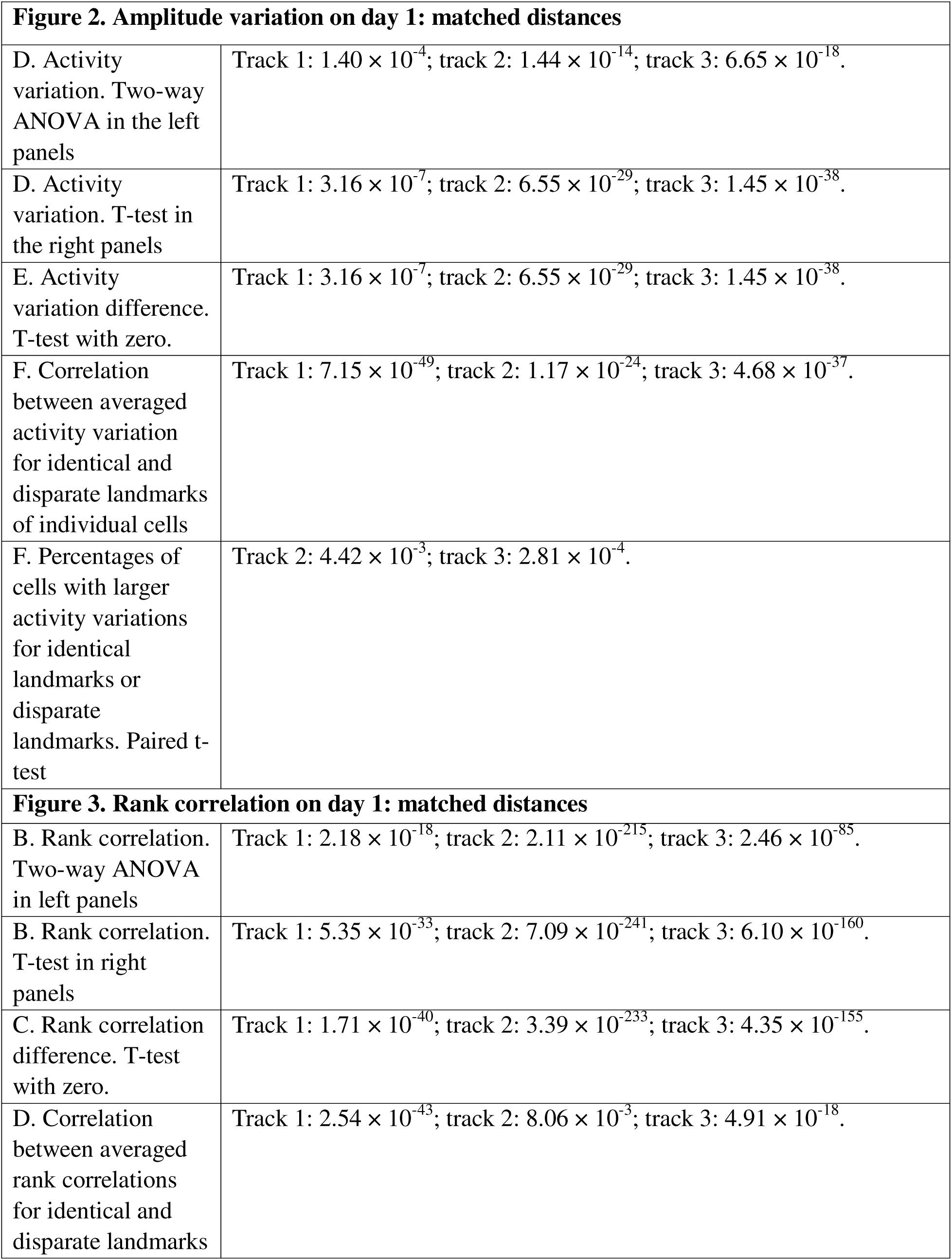
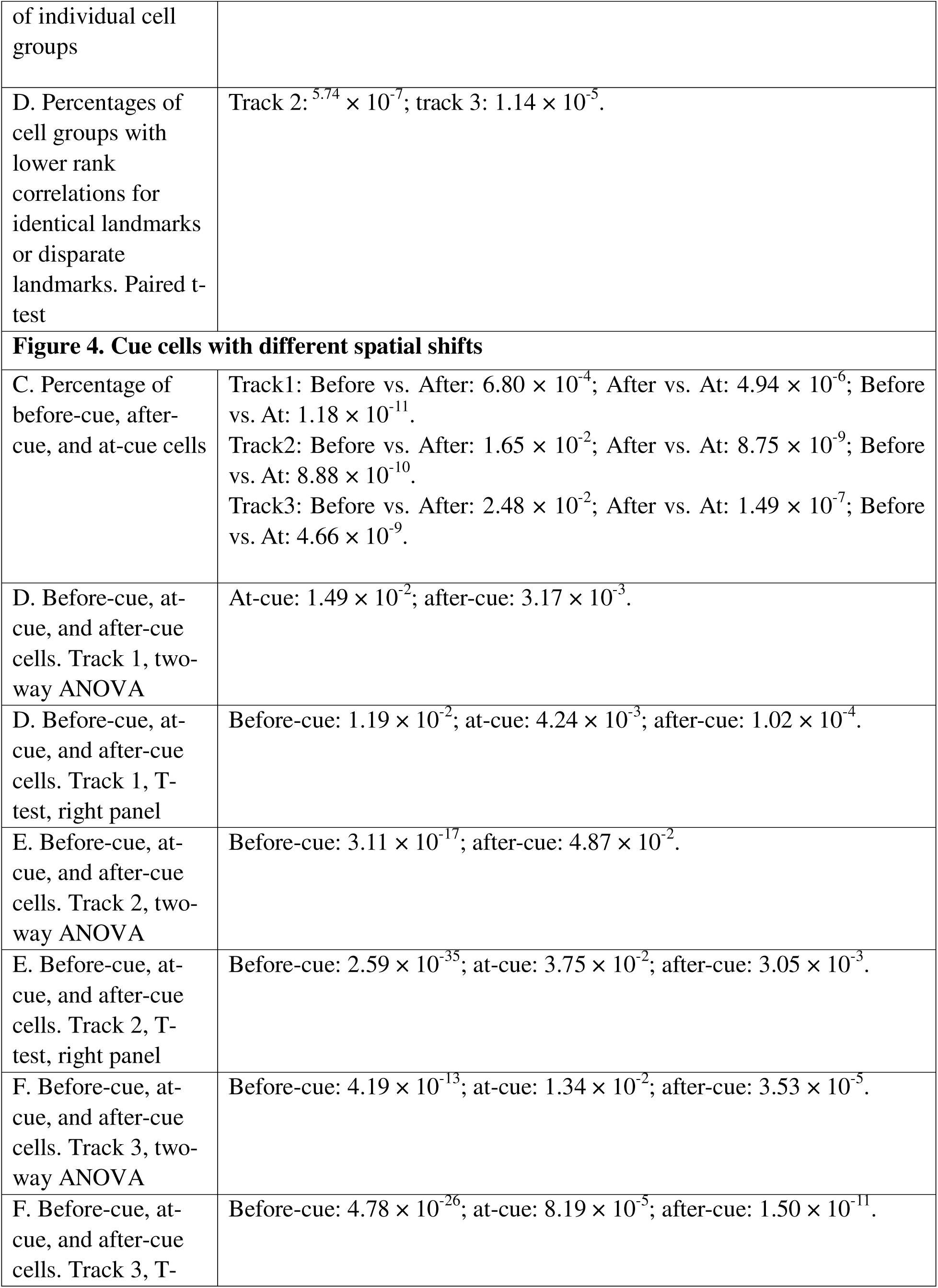
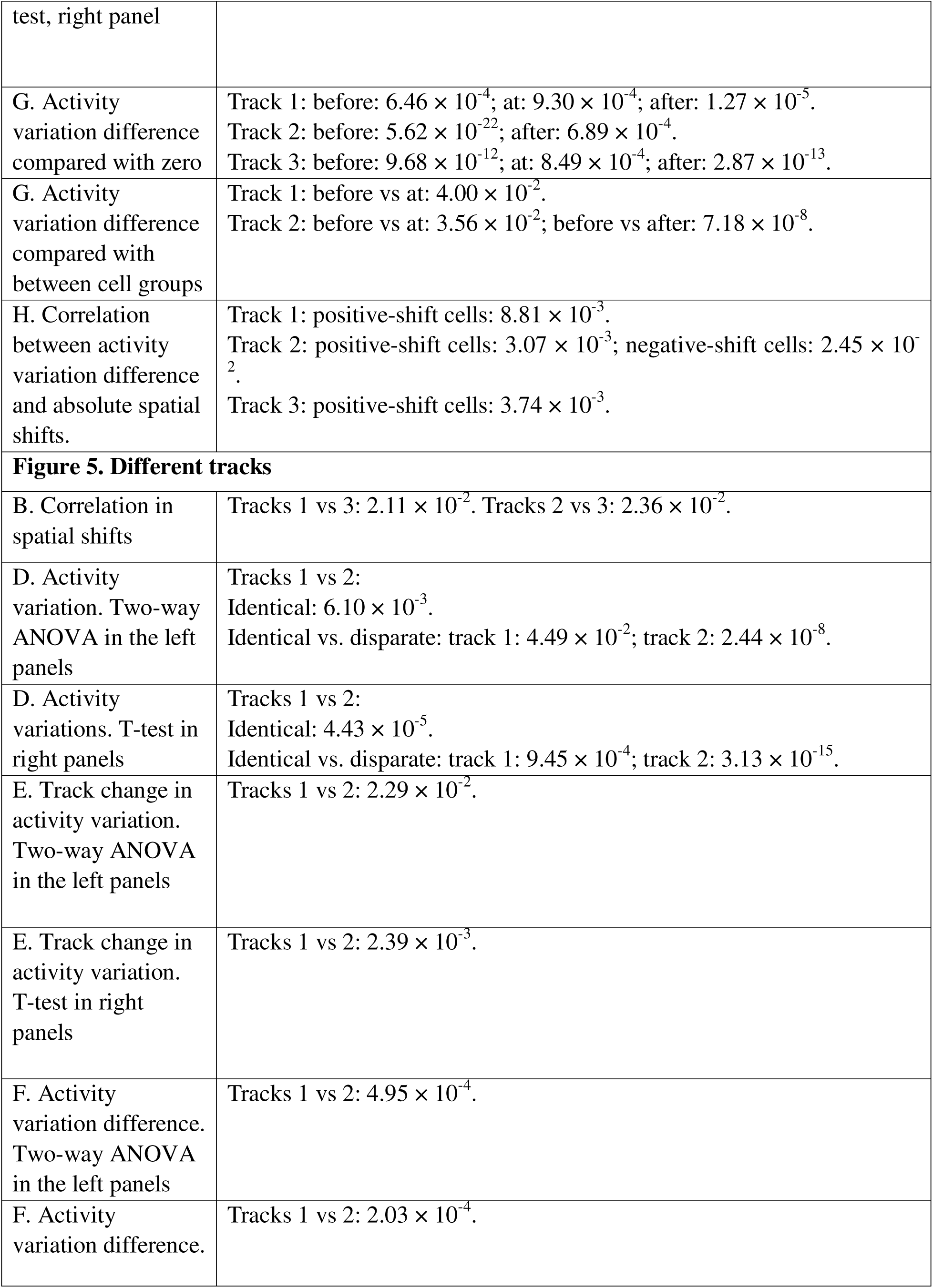
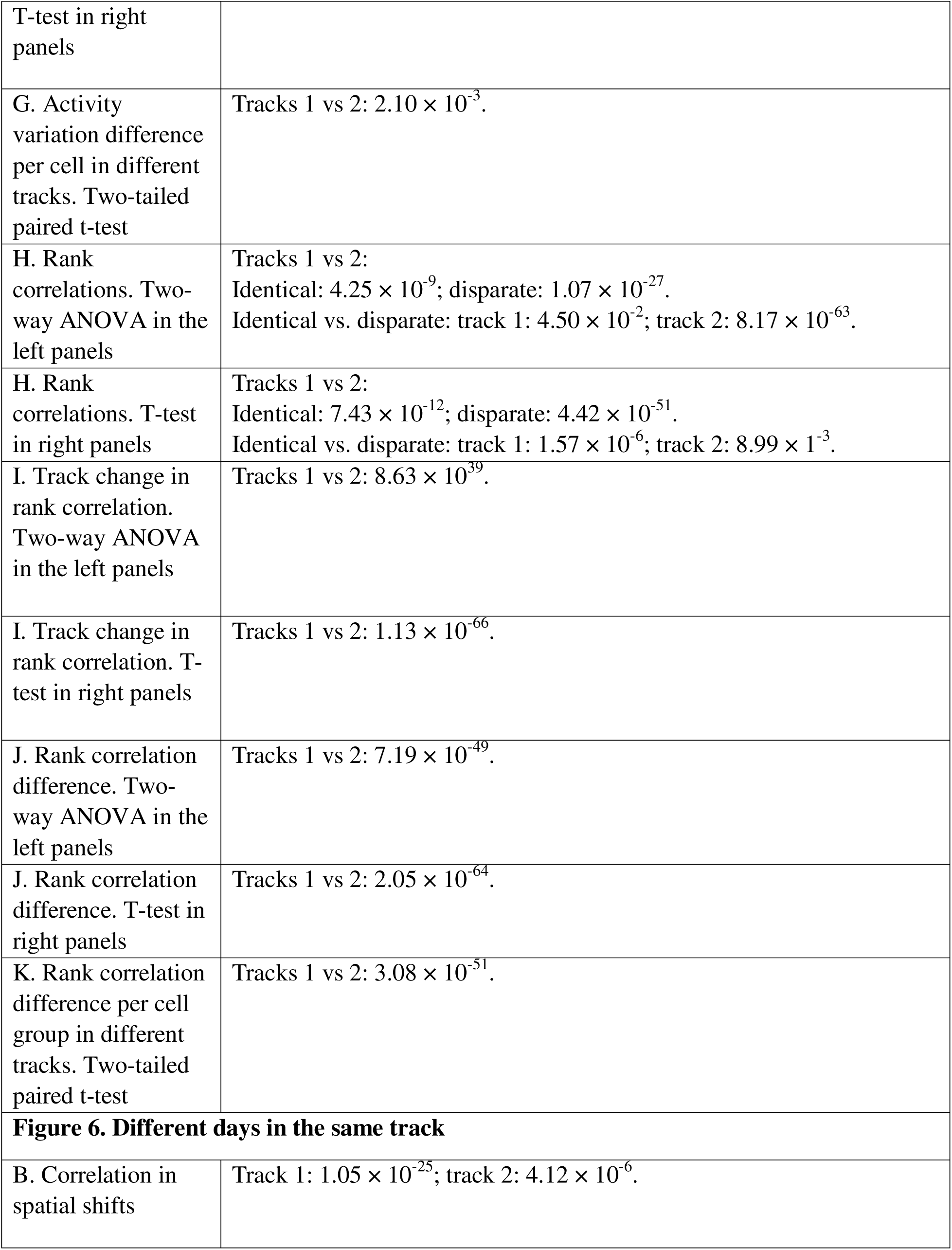
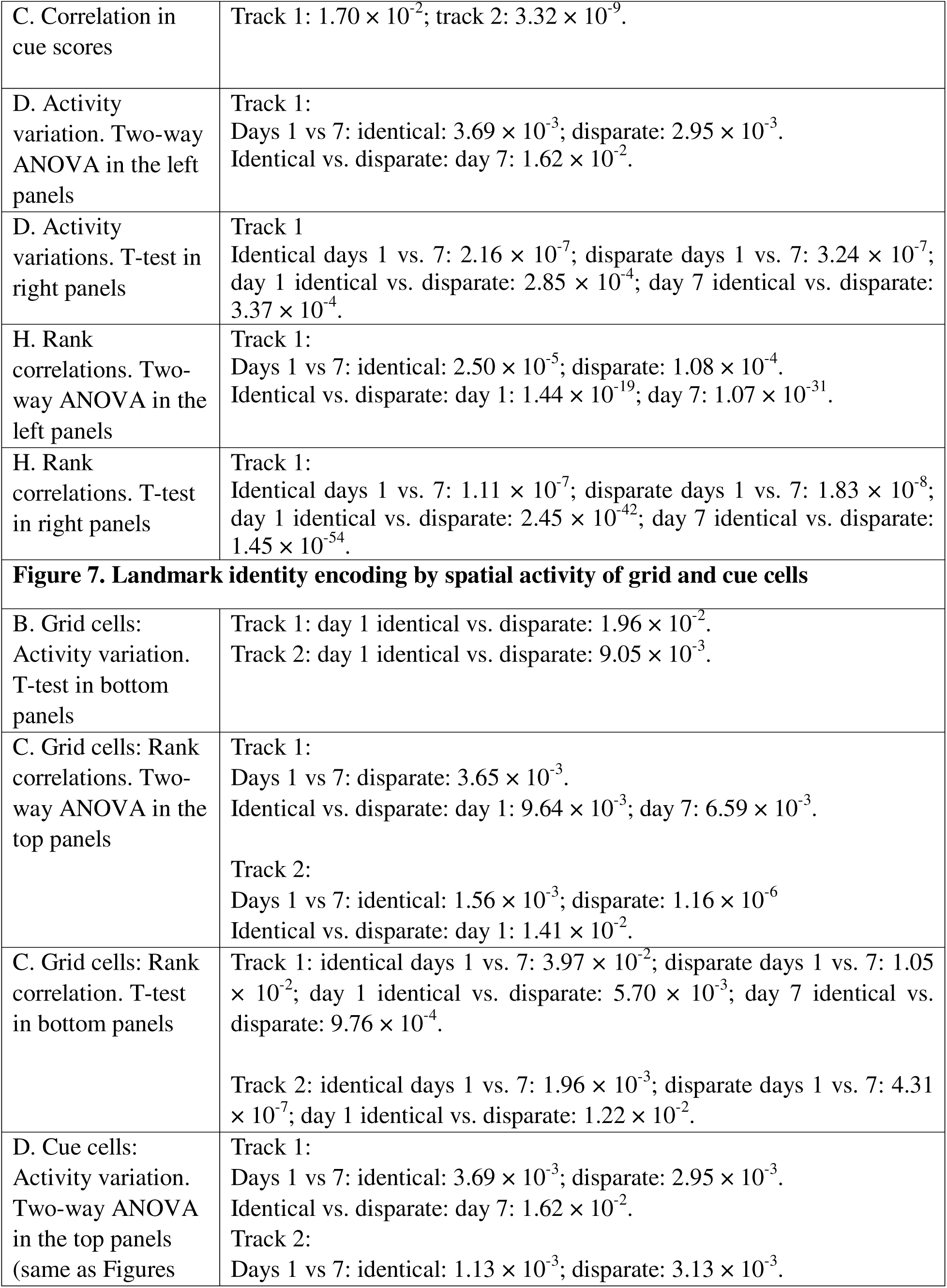
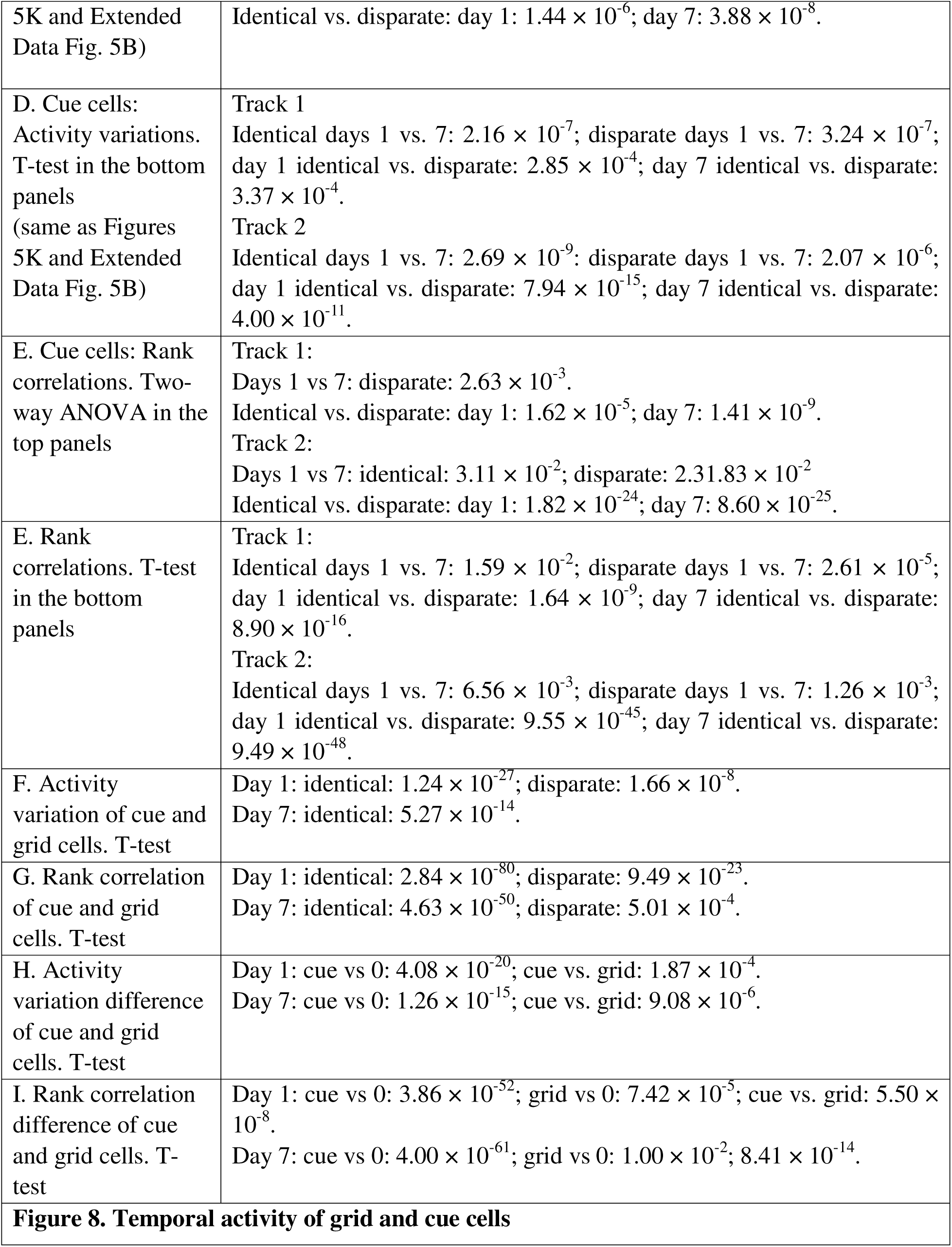
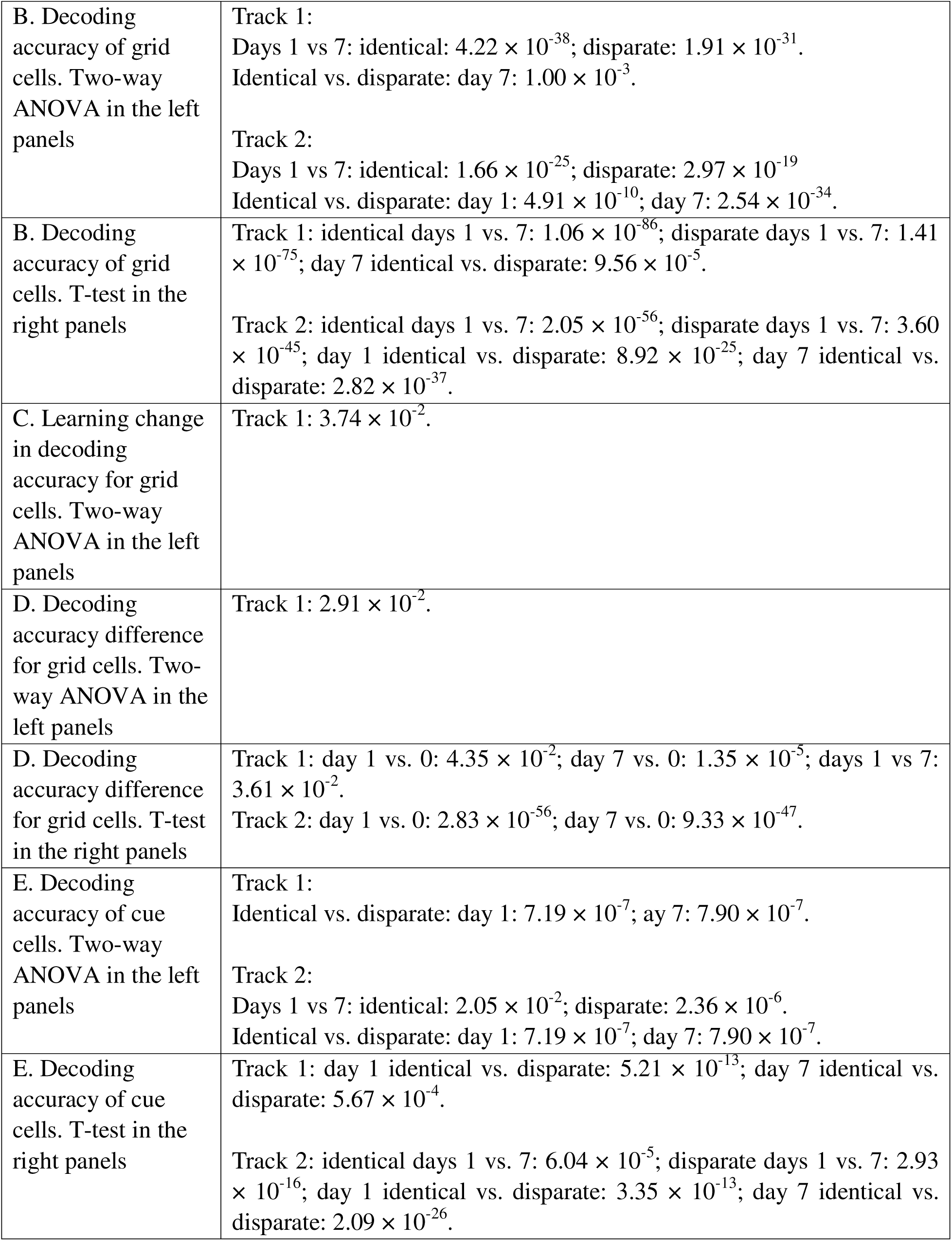
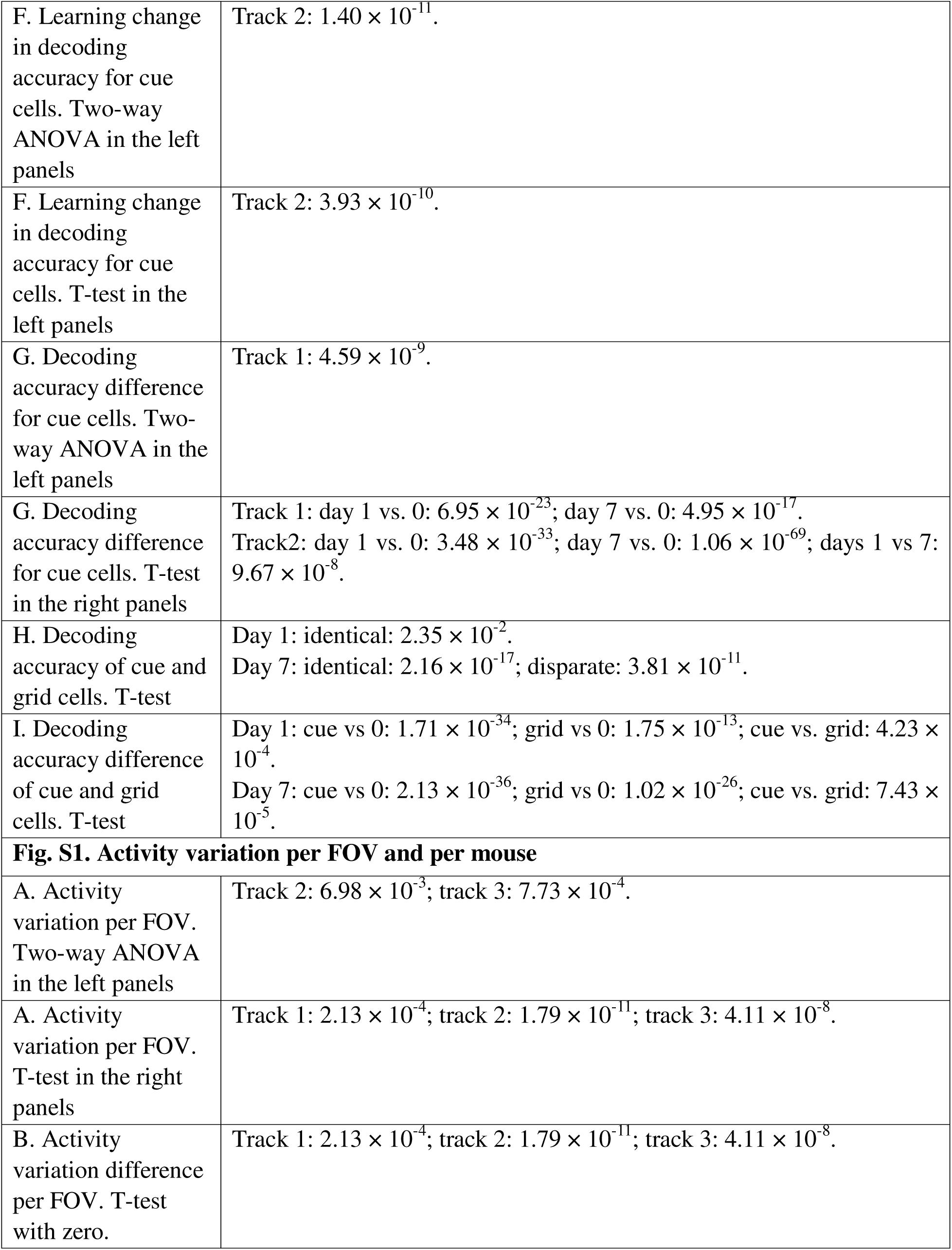
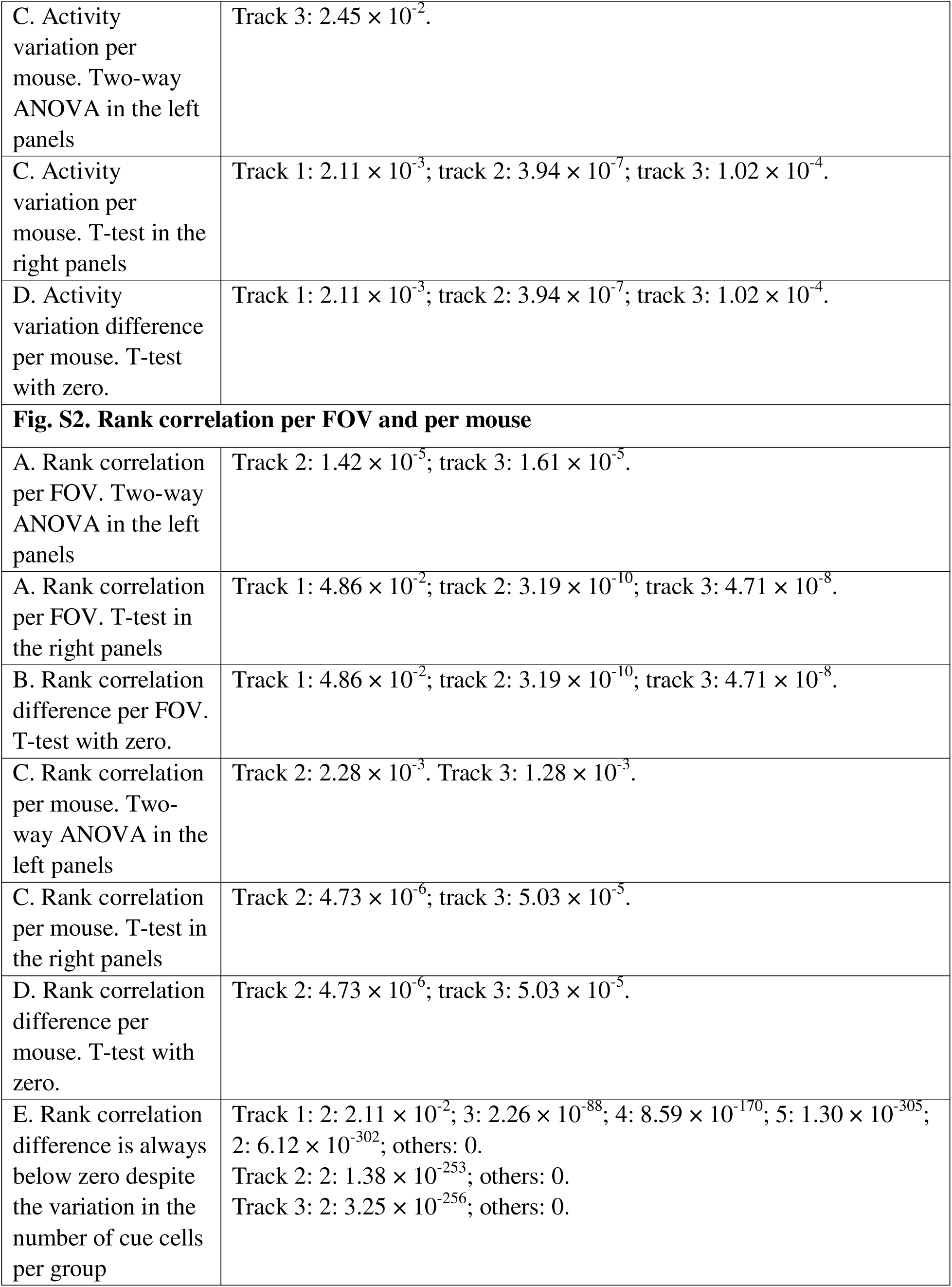
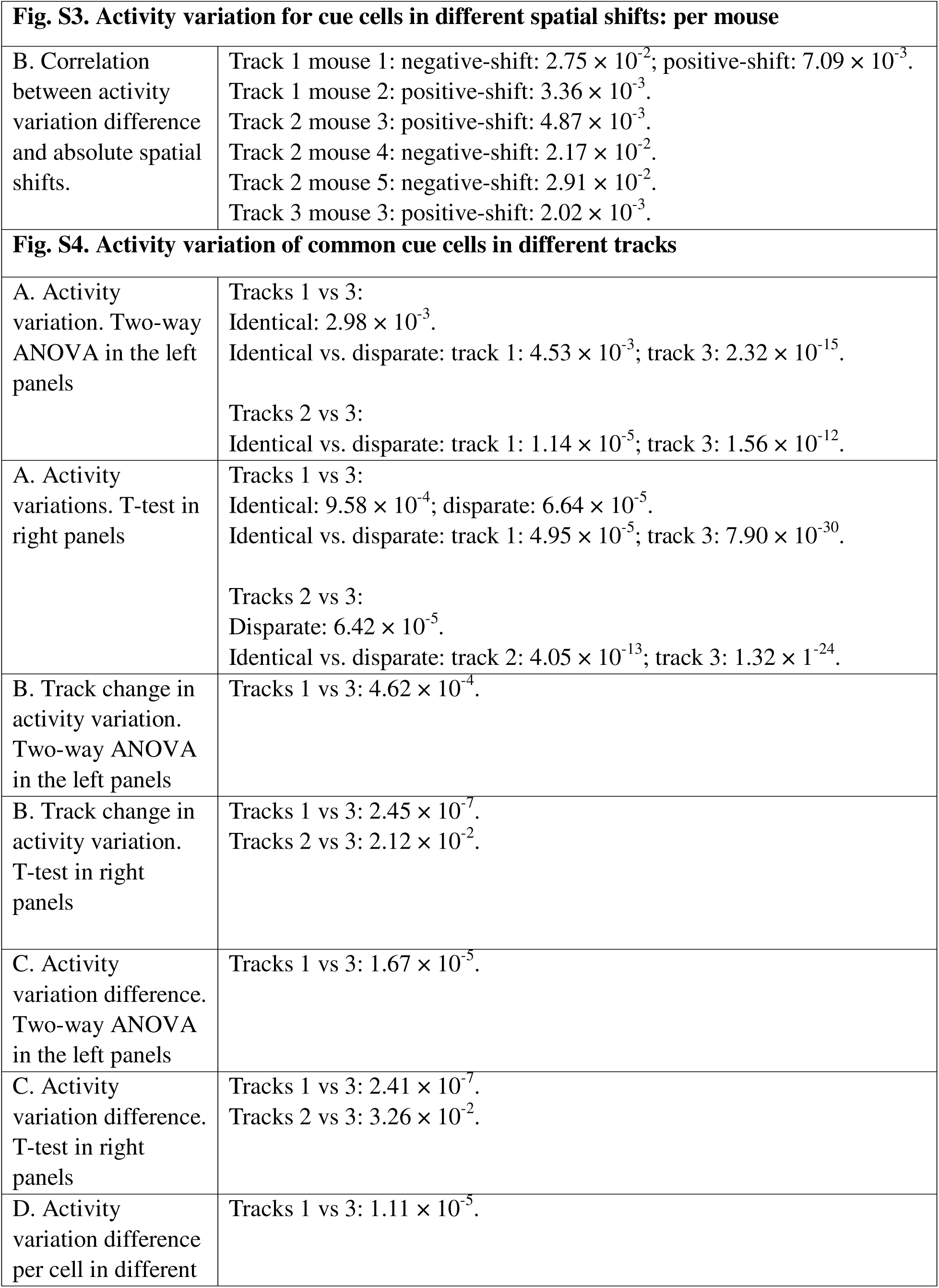
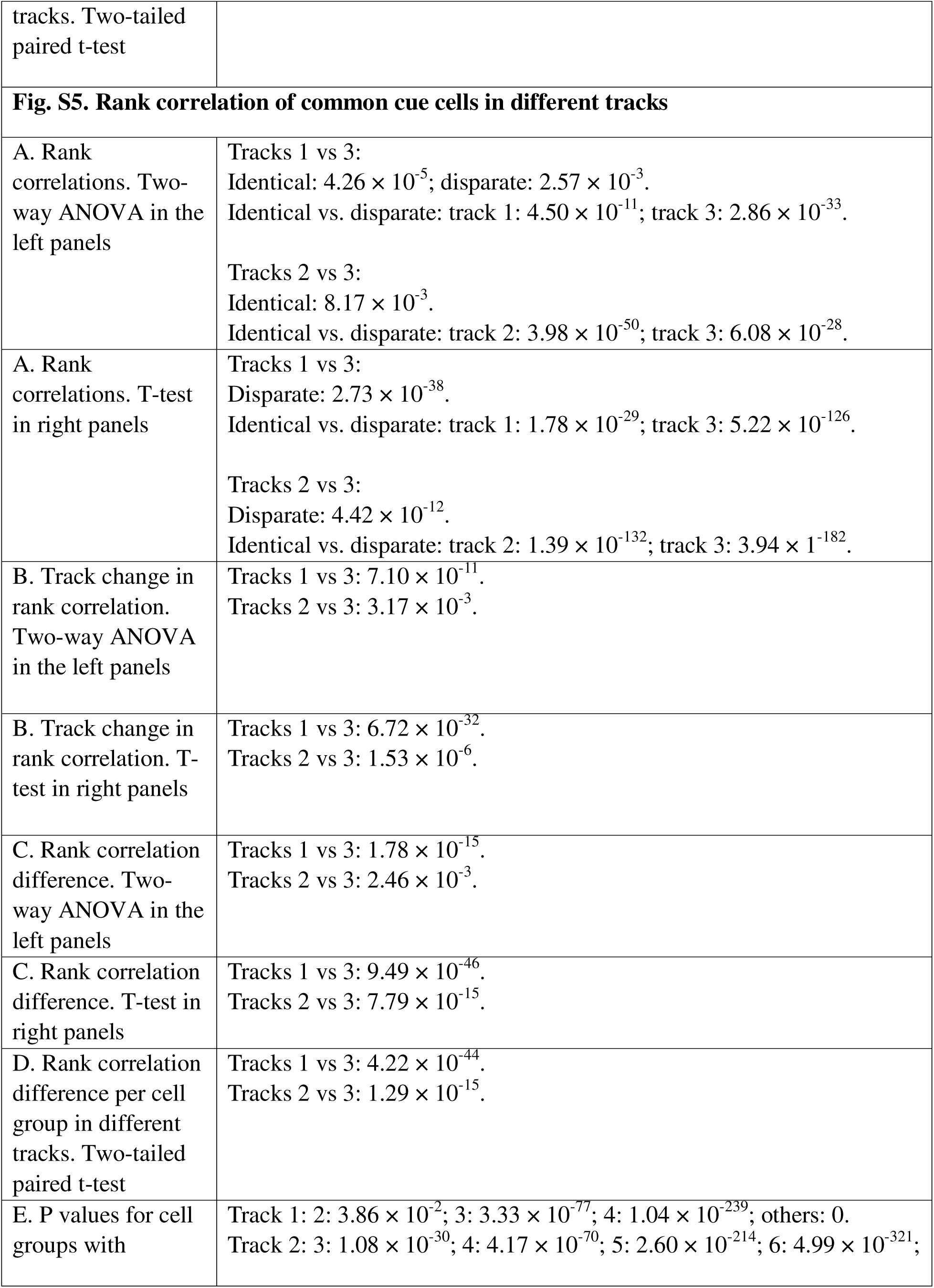
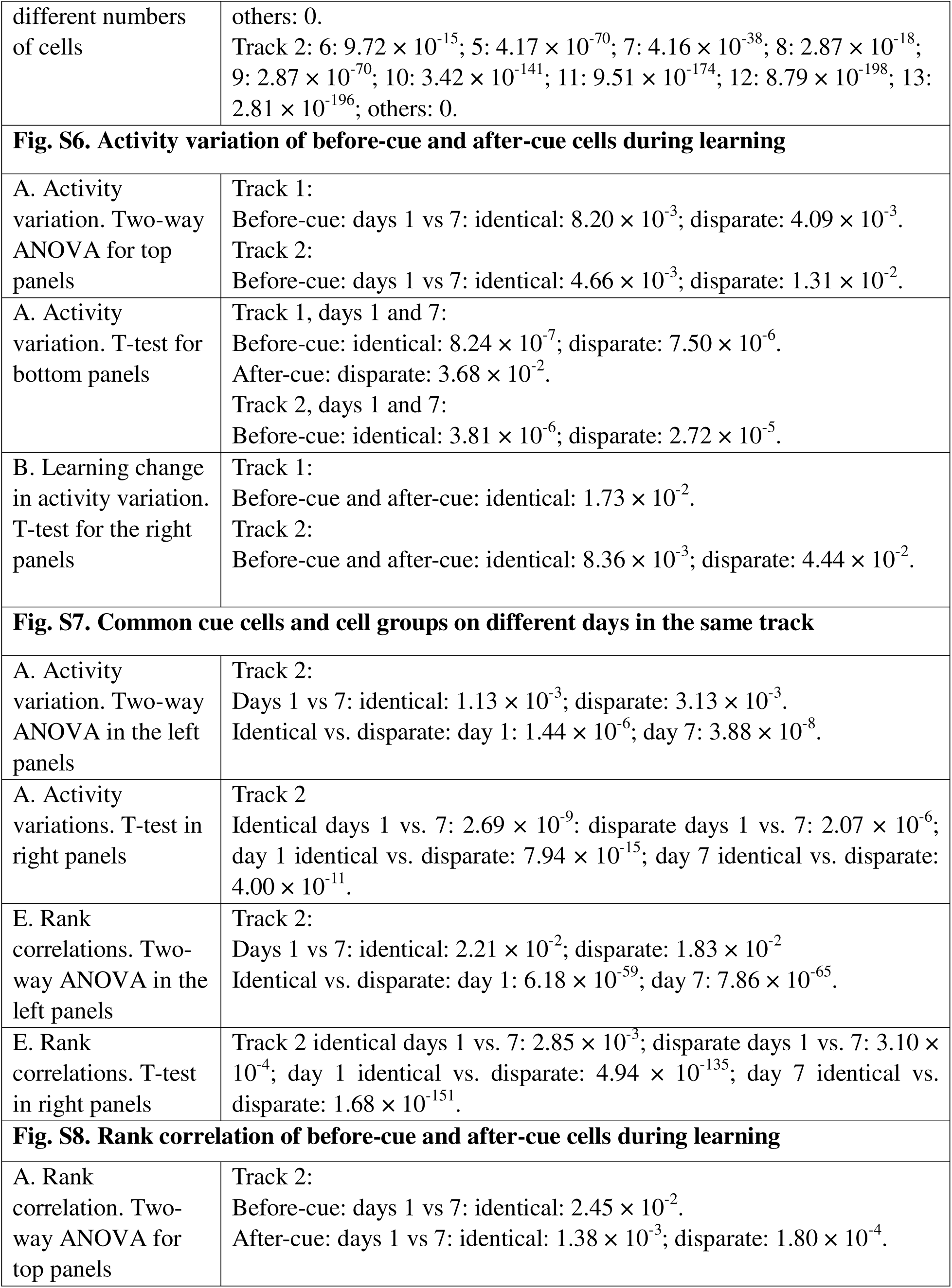
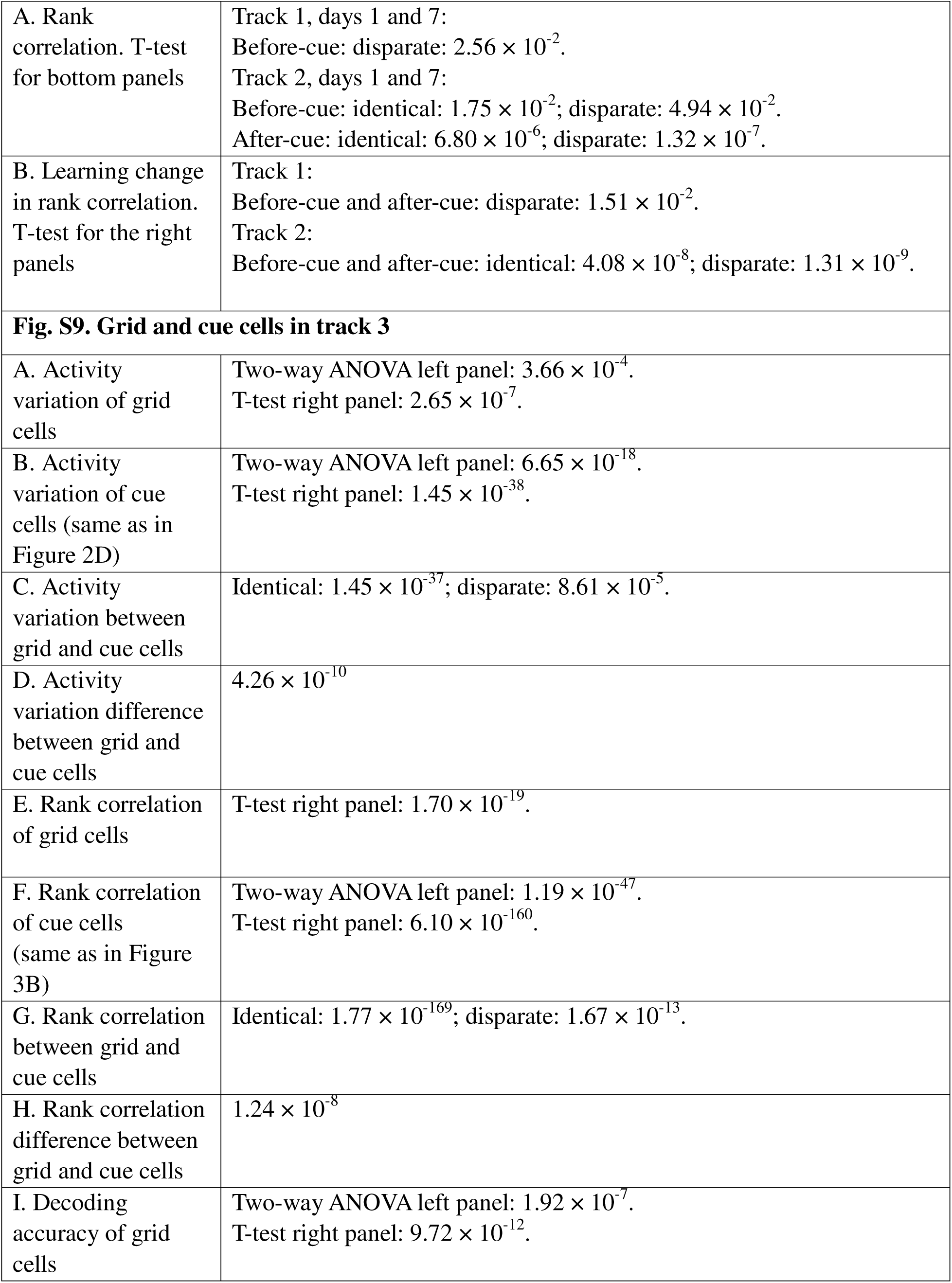
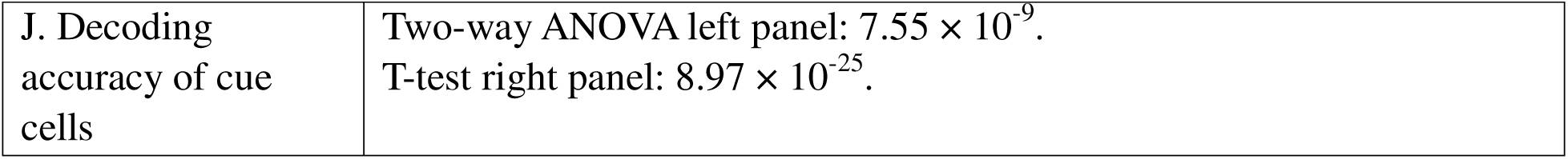
significant p values of all figures.

| **Figure 2. Amplitude variation on day 1: matched distances** |  |
| --- | --- |
| D. Activity variation. Two-way ANOVA in the left panels | Track 1: $1.40 \times 10^{-4}$; track 2: $1.44 \times 10^{-14}$; track 3: $6.65 \times 10^{-18}$. |
| D. Activity variation. T-test in the right panels | Track 1: $3.16 \times 10^{-7}$; track 2: $6.55 \times 10^{-29}$; track 3: $1.45 \times 10^{-38}$. |
| E. Activity variation difference. T-test with zero. | Track 1: $3.16 \times 10^{-7}$; track 2: $6.55 \times 10^{-29}$; track 3: $1.45 \times 10^{-38}$. |
| F. Correlation between averaged activity variation for identical and disparate landmarks of individual cells | Track 1: $7.15 \times 10^{-49}$; track 2: $1.17 \times 10^{-24}$; track 3: $4.68 \times 10^{-37}$. |
| F. Percentages of cells with larger activity variations for identical landmarks or disparate landmarks. Paired t-test | Track 2: $4.42 \times 10^{-3}$; track 3: $2.81 \times 10^{-4}$. |
| **Figure 3. Rank correlation on day 1: matched distances** |  |
| B. Rank correlation. Two-way ANOVA in left panels | Track 1: $2.18 \times 10^{-18}$; track 2: $2.11 \times 10^{-215}$; track 3: $2.46 \times 10^{-85}$. |
| B. Rank correlation. T-test in right panels | Track 1: $5.35 \times 10^{-33}$; track 2: $7.09 \times 10^{-241}$; track 3: $6.10 \times 10^{-160}$. |
| C. Rank correlation difference. T-test with zero. | Track 1: $1.71 \times 10^{-40}$; track 2: $3.39 \times 10^{-233}$; track 3: $4.35 \times 10^{-155}$. |
| D. Correlation between averaged rank correlations for identical and disparate landmarks | Track 1: $2.54 \times 10^{-43}$; track 2: $8.06 \times 10^{-3}$; track 3: $4.91 \times 10^{-18}$. |
| of individual cell groups |  |
| D. Percentages of cell groups with lower rank correlations for identical landmarks or disparate landmarks. Paired t-test | Track 2: $5.74 \times 10^{-7}$; track 3: $1.14 \times 10^{-5}$. |
| **Figure 4. Cue cells with different spatial shifts** |  |
| C. Percentage of before-cue, after-cue, and at-cue cells | Track1: Before vs. After: $6.80 \times 10^{-4}$; After vs. At: $4.94 \times 10^{-6}$; Before vs. At: $1.18 \times 10^{-11}$.<br>Track2: Before vs. After: $1.65 \times 10^{-2}$; After vs. At: $8.75 \times 10^{-9}$; Before vs. At: $8.88 \times 10^{-10}$.<br>Track3: Before vs. After: $2.48 \times 10^{-2}$; After vs. At: $1.49 \times 10^{-7}$; Before vs. At: $4.66 \times 10^{-9}$. |
| D. Before-cue, at-cue, and after-cue cells. Track 1, two-way ANOVA | At-cue: $1.49 \times 10^{-2}$; after-cue: $3.17 \times 10^{-3}$. |
| D. Before-cue, at-cue, and after-cue cells. Track 1, T-test, right panel | Before-cue: $1.19 \times 10^{-2}$; at-cue: $4.24 \times 10^{-3}$; after-cue: $1.02 \times 10^{-4}$. |
| E. Before-cue, at-cue, and after-cue cells. Track 2, two-way ANOVA | Before-cue: $3.11 \times 10^{-17}$; after-cue: $4.87 \times 10^{-2}$. |
| E. Before-cue, at-cue, and after-cue cells. Track 2, T-test, right panel | Before-cue: $2.59 \times 10^{-35}$; at-cue: $3.75 \times 10^{-2}$; after-cue: $3.05 \times 10^{-3}$. |
| F. Before-cue, at-cue, and after-cue cells. Track 3, two-way ANOVA | Before-cue: $4.19 \times 10^{-13}$; at-cue: $1.34 \times 10^{-2}$; after-cue: $3.53 \times 10^{-5}$. |
| F. Before-cue, at-cue, and after-cue cells. Track 3, T- | Before-cue: $4.78 \times 10^{-26}$; at-cue: $8.19 \times 10^{-5}$; after-cue: $1.50 \times 10^{-11}$. |
| test, right panel |  |
| G. Activity variation difference compared with zero | Track 1: before: $6.46 \times 10^{-4}$; at: $9.30 \times 10^{-4}$; after: $1.27 \times 10^{-5}$.<br>Track 2: before: $5.62 \times 10^{-22}$; after: $6.89 \times 10^{-4}$.<br>Track 3: before: $9.68 \times 10^{-12}$; at: $8.49 \times 10^{-4}$; after: $2.87 \times 10^{-13}$. |
| G. Activity variation difference compared with between cell groups | Track 1: before vs at: $4.00 \times 10^{-2}$.<br>Track 2: before vs at: $3.56 \times 10^{-2}$; before vs after: $7.18 \times 10^{-8}$. |
| H. Correlation between activity variation difference and absolute spatial shifts. | Track 1: positive-shift cells: $8.81 \times 10^{-3}$.<br>Track 2: positive-shift cells: $3.07 \times 10^{-3}$; negative-shift cells: $2.45 \times 10^{-2}$.<br>Track 3: positive-shift cells: $3.74 \times 10^{-3}$. |
| **Figure 5. Different tracks** |  |
| B. Correlation in spatial shifts | Tracks 1 vs 3: $2.11 \times 10^{-2}$. Tracks 2 vs 3: $2.36 \times 10^{-2}$. |
| D. Activity variation. Two-way ANOVA in the left panels | Tracks 1 vs 2:<br>Identical: $6.10 \times 10^{-3}$.<br>Identical vs. disparate: track 1: $4.49 \times 10^{-2}$; track 2: $2.44 \times 10^{-8}$. |
| D. Activity variations. T-test in right panels | Tracks 1 vs 2:<br>Identical: $4.43 \times 10^{-5}$.<br>Identical vs. disparate: track 1: $9.45 \times 10^{-4}$; track 2: $3.13 \times 10^{-15}$. |
| E. Track change in activity variation. Two-way ANOVA in the left panels | Tracks 1 vs 2: $2.29 \times 10^{-2}$. |
| E. Track change in activity variation. T-test in right panels | Tracks 1 vs 2: $2.39 \times 10^{-3}$. |
| F. Activity variation difference. Two-way ANOVA in the left panels | Tracks 1 vs 2: $4.95 \times 10^{-4}$. |
| F. Activity variation difference. | Tracks 1 vs 2: $2.03 \times 10^{-4}$. |
| T-test in right panels |  |
| G. Activity variation difference per cell in different tracks. Two-tailed paired t-test | Tracks 1 vs 2: $2.10 \times 10^{-3}$. |
| H. Rank correlations. Two-way ANOVA in the left panels | Tracks 1 vs 2:<br>Identical: $4.25 \times 10^{-9}$; disparate: $1.07 \times 10^{-27}$.<br>Identical vs. disparate: track 1: $4.50 \times 10^{-2}$; track 2: $8.17 \times 10^{-63}$. |
| H. Rank correlations. T-test in right panels | Tracks 1 vs 2:<br>Identical: $7.43 \times 10^{-12}$; disparate: $4.42 \times 10^{-51}$.<br>Identical vs. disparate: track 1: $1.57 \times 10^{-6}$; track 2: $8.99 \times 10^{-3}$. |
| I. Track change in rank correlation. Two-way ANOVA in the left panels | Tracks 1 vs 2: $8.63 \times 10^{39}$. |
| I. Track change in rank correlation. T-test in right panels | Tracks 1 vs 2: $1.13 \times 10^{-66}$. |
| J. Rank correlation difference. Two-way ANOVA in the left panels | Tracks 1 vs 2: $7.19 \times 10^{-49}$. |
| J. Rank correlation difference. T-test in right panels | Tracks 1 vs 2: $2.05 \times 10^{-64}$. |
| K. Rank correlation difference per cell group in different tracks. Two-tailed paired t-test | Tracks 1 vs 2: $3.08 \times 10^{-51}$. |
| **Figure 6. Different days in the same track** |  |
| B. Correlation in spatial shifts | Track 1: $1.05 \times 10^{-25}$; track 2: $4.12 \times 10^{-6}$. |
| C. Correlation in cue scores | Track 1: $1.70 \times 10^{-2}$; track 2: $3.32 \times 10^{-9}$. |
| D. Activity variation. Two-way ANOVA in the left panels | Track 1:<br>Days 1 vs 7: identical: $3.69 \times 10^{-3}$; disparate: $2.95 \times 10^{-3}$.<br>Identical vs. disparate: day 7: $1.62 \times 10^{-2}$. |
| D. Activity variations. T-test in right panels | Track 1<br>Identical days 1 vs. 7: $2.16 \times 10^{-7}$; disparate days 1 vs. 7: $3.24 \times 10^{-7}$;<br>day 1 identical vs. disparate: $2.85 \times 10^{-4}$; day 7 identical vs. disparate: $3.37 \times 10^{-4}$. |
| H. Rank correlations. Two-way ANOVA in the left panels | Track 1:<br>Days 1 vs 7: identical: $2.50 \times 10^{-5}$; disparate: $1.08 \times 10^{-4}$.<br>Identical vs. disparate: day 1: $1.44 \times 10^{-19}$; day 7: $1.07 \times 10^{-31}$. |
| H. Rank correlations. T-test in right panels | Track 1:<br>Identical days 1 vs. 7: $1.11 \times 10^{-7}$; disparate days 1 vs. 7: $1.83 \times 10^{-8}$;<br>day 1 identical vs. disparate: $2.45 \times 10^{-42}$; day 7 identical vs. disparate: $1.45 \times 10^{-54}$. |
| **Figure 7. Landmark identity encoding by spatial activity of grid and cue cells** |  |
| B. Grid cells: Activity variation. T-test in bottom panels | Track 1: day 1 identical vs. disparate: $1.96 \times 10^{-2}$.<br>Track 2: day 1 identical vs. disparate: $9.05 \times 10^{-3}$. |
| C. Grid cells: Rank correlations. Two-way ANOVA in the top panels | Track 1:<br>Days 1 vs 7: disparate: $3.65 \times 10^{-3}$.<br>Identical vs. disparate: day 1: $9.64 \times 10^{-3}$; day 7: $6.59 \times 10^{-3}$.<br>Track 2:<br>Days 1 vs 7: identical: $1.56 \times 10^{-3}$; disparate: $1.16 \times 10^{-6}$<br>Identical vs. disparate: day 1: $1.41 \times 10^{-2}$. |
| C. Grid cells: Rank correlation. T-test in bottom panels | Track 1: identical days 1 vs. 7: $3.97 \times 10^{-2}$; disparate days 1 vs. 7: $1.05 \times 10^{-2}$; day 1 identical vs. disparate: $5.70 \times 10^{-3}$; day 7 identical vs. disparate: $9.76 \times 10^{-4}$.<br>Track 2: identical days 1 vs. 7: $1.96 \times 10^{-3}$; disparate days 1 vs. 7: $4.31 \times 10^{-7}$; day 1 identical vs. disparate: $1.22 \times 10^{-2}$. |
| D. Cue cells: Activity variation. Two-way ANOVA in the top panels (same as Figures | Track 1:<br>Days 1 vs 7: identical: $3.69 \times 10^{-3}$; disparate: $2.95 \times 10^{-3}$.<br>Identical vs. disparate: day 7: $1.62 \times 10^{-2}$.<br>Track 2:<br>Days 1 vs 7: identical: $1.13 \times 10^{-3}$; disparate: $3.13 \times 10^{-3}$. |
| 5K and Extended Data Fig. 5B) | Identical vs. disparate: day 1: $1.44 \times 10^{-6}$; day 7: $3.88 \times 10^{-8}$. |
| D. Cue cells: Activity variations. T-test in the bottom panels (same as Figures 5K and Extended Data Fig. 5B) | Track 1<br>Identical days 1 vs. 7: $2.16 \times 10^{-7}$; disparate days 1 vs. 7: $3.24 \times 10^{-7}$; day 1 identical vs. disparate: $2.85 \times 10^{-4}$; day 7 identical vs. disparate: $3.37 \times 10^{-4}$.<br>Track 2<br>Identical days 1 vs. 7: $2.69 \times 10^{-9}$; disparate days 1 vs. 7: $2.07 \times 10^{-6}$; day 1 identical vs. disparate: $7.94 \times 10^{-15}$; day 7 identical vs. disparate: $4.00 \times 10^{-11}$. |
| E. Cue cells: Rank correlations. Two-way ANOVA in the top panels | Track 1:<br>Days 1 vs 7: disparate: $2.63 \times 10^{-3}$.<br>Identical vs. disparate: day 1: $1.62 \times 10^{-5}$; day 7: $1.41 \times 10^{-9}$.<br>Track 2:<br>Days 1 vs 7: identical: $3.11 \times 10^{-2}$; disparate: $2.31.83 \times 10^{-2}$<br>Identical vs. disparate: day 1: $1.82 \times 10^{-24}$; day 7: $8.60 \times 10^{-25}$. |
| E. Rank correlations. T-test in the bottom panels | Track 1:<br>Identical days 1 vs. 7: $1.59 \times 10^{-2}$; disparate days 1 vs. 7: $2.61 \times 10^{-5}$; day 1 identical vs. disparate: $1.64 \times 10^{-9}$; day 7 identical vs. disparate: $8.90 \times 10^{-16}$.<br>Track 2:<br>Identical days 1 vs. 7: $6.56 \times 10^{-3}$; disparate days 1 vs. 7: $1.26 \times 10^{-3}$; day 1 identical vs. disparate: $9.55 \times 10^{-45}$; day 7 identical vs. disparate: $9.49 \times 10^{-48}$. |
| F. Activity variation of cue and grid cells. T-test | Day 1: identical: $1.24 \times 10^{-27}$; disparate: $1.66 \times 10^{-8}$.<br>Day 7: identical: $5.27 \times 10^{-14}$. |
| G. Rank correlation of cue and grid cells. T-test | Day 1: identical: $2.84 \times 10^{-80}$; disparate: $9.49 \times 10^{-23}$.<br>Day 7: identical: $4.63 \times 10^{-50}$; disparate: $5.01 \times 10^{-4}$. |
| H. Activity variation difference of cue and grid cells. T-test | Day 1: cue vs 0: $4.08 \times 10^{-20}$; cue vs. grid: $1.87 \times 10^{-4}$.<br>Day 7: cue vs 0: $1.26 \times 10^{-15}$; cue vs. grid: $9.08 \times 10^{-6}$. |
| I. Rank correlation difference of cue and grid cells. T-test | Day 1: cue vs 0: $3.86 \times 10^{-52}$; grid vs 0: $7.42 \times 10^{-5}$; cue vs. grid: $5.50 \times 10^{-8}$.<br>Day 7: cue vs 0: $4.00 \times 10^{-61}$; grid vs 0: $1.00 \times 10^{-2}$; $8.41 \times 10^{-14}$. |
| **Figure 8. Temporal activity of grid and cue cells** |  |
| B. Decoding accuracy of grid cells. Two-way ANOVA in the left panels | Track 1:<br>Days 1 vs 7: identical: $4.22 \times 10^{-38}$; disparate: $1.91 \times 10^{-31}$.<br>Identical vs. disparate: day 7: $1.00 \times 10^{-3}$.<br>Track 2:<br>Days 1 vs 7: identical: $1.66 \times 10^{-25}$; disparate: $2.97 \times 10^{-19}$<br>Identical vs. disparate: day 1: $4.91 \times 10^{-10}$; day 7: $2.54 \times 10^{-34}$. |
| B. Decoding accuracy of grid cells. T-test in the right panels | Track 1: identical days 1 vs. 7: $1.06 \times 10^{-86}$; disparate days 1 vs. 7: $1.41 \times 10^{-75}$; day 7 identical vs. disparate: $9.56 \times 10^{-5}$.<br>Track 2: identical days 1 vs. 7: $2.05 \times 10^{-56}$; disparate days 1 vs. 7: $3.60 \times 10^{-45}$; day 1 identical vs. disparate: $8.92 \times 10^{-25}$; day 7 identical vs. disparate: $2.82 \times 10^{-37}$. |
| C. Learning change in decoding accuracy for grid cells. Two-way ANOVA in the left panels | Track 1: $3.74 \times 10^{-2}$. |
| D. Decoding accuracy difference for grid cells. Two-way ANOVA in the left panels | Track 1: $2.91 \times 10^{-2}$. |
| D. Decoding accuracy difference for grid cells. T-test in the right panels | Track 1: day 1 vs. 0: $4.35 \times 10^{-2}$; day 7 vs. 0: $1.35 \times 10^{-5}$; days 1 vs 7: $3.61 \times 10^{-2}$.<br>Track 2: day 1 vs. 0: $2.83 \times 10^{-56}$; day 7 vs. 0: $9.33 \times 10^{-47}$. |
| E. Decoding accuracy of cue cells. Two-way ANOVA in the left panels | Track 1:<br>Identical vs. disparate: day 1: $7.19 \times 10^{-7}$; day 7: $7.90 \times 10^{-7}$.<br>Track 2:<br>Days 1 vs 7: identical: $2.05 \times 10^{-2}$; disparate: $2.36 \times 10^{-6}$.<br>Identical vs. disparate: day 1: $7.19 \times 10^{-7}$; day 7: $7.90 \times 10^{-7}$. |
| E. Decoding accuracy of cue cells. T-test in the right panels | Track 1: day 1 identical vs. disparate: $5.21 \times 10^{-13}$; day 7 identical vs. disparate: $5.67 \times 10^{-4}$.<br>Track 2: identical days 1 vs. 7: $6.04 \times 10^{-5}$; disparate days 1 vs. 7: $2.93 \times 10^{-16}$; day 1 identical vs. disparate: $3.35 \times 10^{-13}$; day 7 identical vs. disparate: $2.09 \times 10^{-26}$. |
| F. Learning change in decoding accuracy for cue cells. Two-way ANOVA in the left panels | Track 2: $1.40 \times 10^{-11}$. |
| F. Learning change in decoding accuracy for cue cells. T-test in the left panels | Track 2: $3.93 \times 10^{-10}$. |
| G. Decoding accuracy difference for cue cells. Two-way ANOVA in the left panels | Track 1: $4.59 \times 10^{-9}$. |
| G. Decoding accuracy difference for cue cells. T-test in the right panels | Track 1: day 1 vs. 0: $6.95 \times 10^{-23}$; day 7 vs. 0: $4.95 \times 10^{-17}$.<br>Track2: day 1 vs. 0: $3.48 \times 10^{-33}$; day 7 vs. 0: $1.06 \times 10^{-69}$; days 1 vs 7: $9.67 \times 10^{-8}$. |
| H. Decoding accuracy of cue and grid cells. T-test | Day 1: identical: $2.35 \times 10^{-2}$.<br>Day 7: identical: $2.16 \times 10^{-17}$; disparate: $3.81 \times 10^{-11}$. |
| I. Decoding accuracy difference of cue and grid cells. T-test | Day 1: cue vs 0: $1.71 \times 10^{-34}$; grid vs 0: $1.75 \times 10^{-13}$; cue vs. grid: $4.23 \times 10^{-4}$.<br>Day 7: cue vs 0: $2.13 \times 10^{-36}$; grid vs 0: $1.02 \times 10^{-26}$; cue vs. grid: $7.43 \times 10^{-5}$. |
| **Fig. S1. Activity variation per FOV and per mouse** |  |
| A. Activity variation per FOV. Two-way ANOVA in the left panels | Track 2: $6.98 \times 10^{-3}$; track 3: $7.73 \times 10^{-4}$. |
| A. Activity variation per FOV. T-test in the right panels | Track 1: $2.13 \times 10^{-4}$; track 2: $1.79 \times 10^{-11}$; track 3: $4.11 \times 10^{-8}$. |
| B. Activity variation difference per FOV. T-test with zero. | Track 1: $2.13 \times 10^{-4}$; track 2: $1.79 \times 10^{-11}$; track 3: $4.11 \times 10^{-8}$. |
| C. Activity variation per mouse. Two-way ANOVA in the left panels | Track 3: $2.45 \times 10^{-2}$. |
| C. Activity variation per mouse. T-test in the right panels | Track 1: $2.11 \times 10^{-3}$; track 2: $3.94 \times 10^{-7}$; track 3: $1.02 \times 10^{-4}$. |
| D. Activity variation difference per mouse. T-test with zero. | Track 1: $2.11 \times 10^{-3}$; track 2: $3.94 \times 10^{-7}$; track 3: $1.02 \times 10^{-4}$. |
| **Fig. S2. Rank correlation per FOV and per mouse** |  |
| A. Rank correlation per FOV. Two-way ANOVA in the left panels | Track 2: $1.42 \times 10^{-5}$; track 3: $1.61 \times 10^{-5}$. |
| A. Rank correlation per FOV. T-test in the right panels | Track 1: $4.86 \times 10^{-2}$; track 2: $3.19 \times 10^{-10}$; track 3: $4.71 \times 10^{-8}$. |
| B. Rank correlation difference per FOV. T-test with zero. | Track 1: $4.86 \times 10^{-2}$; track 2: $3.19 \times 10^{-10}$; track 3: $4.71 \times 10^{-8}$. |
| C. Rank correlation per mouse. Two-way ANOVA in the left panels | Track 2: $2.28 \times 10^{-3}$. Track 3: $1.28 \times 10^{-3}$. |
| C. Rank correlation per mouse. T-test in the right panels | Track 2: $4.73 \times 10^{-6}$; track 3: $5.03 \times 10^{-5}$. |
| D. Rank correlation difference per mouse. T-test with zero. | Track 2: $4.73 \times 10^{-6}$; track 3: $5.03 \times 10^{-5}$. |
| E. Rank correlation difference is always below zero despite the variation in the number of cue cells per group | Track 1: 2: $2.11 \times 10^{-2}$; 3: $2.26 \times 10^{-88}$; 4: $8.59 \times 10^{-170}$; 5: $1.30 \times 10^{-305}$; 2: $6.12 \times 10^{-302}$; others: 0.<br>Track 2: 2: $1.38 \times 10^{-253}$; others: 0.<br>Track 3: 2: $3.25 \times 10^{-256}$; others: 0. |

| **Fig. S3. Activity variation for cue cells in different spatial shifts: per mouse** |  |
| --- | --- |
| B. Correlation between activity variation difference and absolute spatial shifts. | Track 1 mouse 1: negative-shift: $2.75 \times 10^{-2}$; positive-shift: $7.09 \times 10^{-3}$.<br>Track 1 mouse 2: positive-shift: $3.36 \times 10^{-3}$.<br>Track 2 mouse 3: positive-shift: $4.87 \times 10^{-3}$.<br>Track 2 mouse 4: negative-shift: $2.17 \times 10^{-2}$.<br>Track 2 mouse 5: negative-shift: $2.91 \times 10^{-2}$.<br>Track 3 mouse 3: positive-shift: $2.02 \times 10^{-3}$. |
| **Fig. S4. Activity variation of common cue cells in different tracks** |  |
| A. Activity variation. Two-way ANOVA in the left panels | Tracks 1 vs 3:<br>Identical: $2.98 \times 10^{-3}$.<br>Identical vs. disparate: track 1: $4.53 \times 10^{-3}$; track 3: $2.32 \times 10^{-15}$.<br>Tracks 2 vs 3:<br>Identical vs. disparate: track 1: $1.14 \times 10^{-5}$; track 3: $1.56 \times 10^{-12}$. |
| A. Activity variations. T-test in right panels | Tracks 1 vs 3:<br>Identical: $9.58 \times 10^{-4}$; disparate: $6.64 \times 10^{-5}$.<br>Identical vs. disparate: track 1: $4.95 \times 10^{-5}$; track 3: $7.90 \times 10^{-30}$.<br>Tracks 2 vs 3:<br>Disparate: $6.42 \times 10^{-5}$.<br>Identical vs. disparate: track 2: $4.05 \times 10^{-13}$; track 3: $1.32 \times 10^{-24}$. |
| B. Track change in activity variation. Two-way ANOVA in the left panels | Tracks 1 vs 3: $4.62 \times 10^{-4}$. |
| B. Track change in activity variation. T-test in right panels | Tracks 1 vs 3: $2.45 \times 10^{-7}$.<br>Tracks 2 vs 3: $2.12 \times 10^{-2}$. |
| C. Activity variation difference. Two-way ANOVA in the left panels | Tracks 1 vs 3: $1.67 \times 10^{-5}$. |
| C. Activity variation difference. T-test in right panels | Tracks 1 vs 3: $2.41 \times 10^{-7}$.<br>Tracks 2 vs 3: $3.26 \times 10^{-2}$. |
| D. Activity variation difference per cell in different | Tracks 1 vs 3: $1.11 \times 10^{-5}$. |
| tracks. Two-tailed paired t-test |  |
| **Fig. S5. Rank correlation of common cue cells in different tracks** |  |
| A. Rank correlations. Two-way ANOVA in the left panels | Tracks 1 vs 3:<br>Identical: $4.26 \times 10^{-5}$; disparate: $2.57 \times 10^{-3}$.<br>Identical vs. disparate: track 1: $4.50 \times 10^{-11}$; track 3: $2.86 \times 10^{-33}$.<br>Tracks 2 vs 3:<br>Identical: $8.17 \times 10^{-3}$.<br>Identical vs. disparate: track 2: $3.98 \times 10^{-50}$; track 3: $6.08 \times 10^{-28}$. |
| A. Rank correlations. T-test in right panels | Tracks 1 vs 3:<br>Disparate: $2.73 \times 10^{-38}$.<br>Identical vs. disparate: track 1: $1.78 \times 10^{-29}$; track 3: $5.22 \times 10^{-126}$.<br>Tracks 2 vs 3:<br>Disparate: $4.42 \times 10^{-12}$.<br>Identical vs. disparate: track 2: $1.39 \times 10^{-132}$; track 3: $3.94 \times 10^{-182}$. |
| B. Track change in rank correlation. Two-way ANOVA in the left panels | Tracks 1 vs 3: $7.10 \times 10^{-11}$.<br>Tracks 2 vs 3: $3.17 \times 10^{-3}$. |
| B. Track change in rank correlation. T-test in right panels | Tracks 1 vs 3: $6.72 \times 10^{-32}$.<br>Tracks 2 vs 3: $1.53 \times 10^{-6}$. |
| C. Rank correlation difference. Two-way ANOVA in the left panels | Tracks 1 vs 3: $1.78 \times 10^{-15}$.<br>Tracks 2 vs 3: $2.46 \times 10^{-3}$. |
| C. Rank correlation difference. T-test in right panels | Tracks 1 vs 3: $9.49 \times 10^{-46}$.<br>Tracks 2 vs 3: $7.79 \times 10^{-15}$. |
| D. Rank correlation difference per cell group in different tracks. Two-tailed paired t-test | Tracks 1 vs 3: $4.22 \times 10^{-44}$.<br>Tracks 2 vs 3: $1.29 \times 10^{-15}$. |
| E. P values for cell groups with | Track 1: 2: $3.86 \times 10^{-2}$; 3: $3.33 \times 10^{-77}$; 4: $1.04 \times 10^{-239}$; others: 0.<br>Track 2: 3: $1.08 \times 10^{-30}$; 4: $4.17 \times 10^{-70}$; 5: $2.60 \times 10^{-214}$; 6: $4.99 \times 10^{-321}$; |
| different numbers of cells | others: 0.<br>Track 2: 6: $9.72 \times 10^{-15}$; 5: $4.17 \times 10^{-70}$; 7: $4.16 \times 10^{-38}$; 8: $2.87 \times 10^{-18}$; 9: $2.87 \times 10^{-70}$; 10: $3.42 \times 10^{-141}$; 11: $9.51 \times 10^{-174}$; 12: $8.79 \times 10^{-198}$; 13: $2.81 \times 10^{-196}$; others: 0. |
| **Fig. S6. Activity variation of before-cue and after-cue cells during learning** |  |
| A. Activity variation. Two-way ANOVA for top panels | Track 1:<br>Before-cue: days 1 vs 7: identical: $8.20 \times 10^{-3}$; disparate: $4.09 \times 10^{-3}$.<br>Track 2:<br>Before-cue: days 1 vs 7: identical: $4.66 \times 10^{-3}$; disparate: $1.31 \times 10^{-2}$. |
| A. Activity variation. T-test for bottom panels | Track 1, days 1 and 7:<br>Before-cue: identical: $8.24 \times 10^{-7}$; disparate: $7.50 \times 10^{-6}$.<br>After-cue: disparate: $3.68 \times 10^{-2}$.<br>Track 2, days 1 and 7:<br>Before-cue: identical: $3.81 \times 10^{-6}$; disparate: $2.72 \times 10^{-5}$. |
| B. Learning change in activity variation. T-test for the right panels | Track 1:<br>Before-cue and after-cue: identical: $1.73 \times 10^{-2}$.<br>Track 2:<br>Before-cue and after-cue: identical: $8.36 \times 10^{-3}$; disparate: $4.44 \times 10^{-2}$. |
| **Fig. S7. Common cue cells and cell groups on different days in the same track** |  |
| A. Activity variation. Two-way ANOVA in the left panels | Track 2:<br>Days 1 vs 7: identical: $1.13 \times 10^{-3}$; disparate: $3.13 \times 10^{-3}$.<br>Identical vs. disparate: day 1: $1.44 \times 10^{-6}$; day 7: $3.88 \times 10^{-8}$. |
| A. Activity variations. T-test in right panels | Track 2<br>Identical days 1 vs. 7: $2.69 \times 10^{-9}$; disparate days 1 vs. 7: $2.07 \times 10^{-6}$; day 1 identical vs. disparate: $7.94 \times 10^{-15}$; day 7 identical vs. disparate: $4.00 \times 10^{-11}$. |
| E. Rank correlations. Two-way ANOVA in the left panels | Track 2:<br>Days 1 vs 7: identical: $2.21 \times 10^{-2}$; disparate: $1.83 \times 10^{-2}$<br>Identical vs. disparate: day 1: $6.18 \times 10^{-59}$; day 7: $7.86 \times 10^{-65}$. |
| E. Rank correlations. T-test in right panels | Track 2 identical days 1 vs. 7: $2.85 \times 10^{-3}$; disparate days 1 vs. 7: $3.10 \times 10^{-4}$; day 1 identical vs. disparate: $4.94 \times 10^{-135}$; day 7 identical vs. disparate: $1.68 \times 10^{-151}$. |
| **Fig. S8. Rank correlation of before-cue and after-cue cells during learning** |  |
| A. Rank correlation. Two-way ANOVA for top panels | Track 2:<br>Before-cue: days 1 vs 7: identical: $2.45 \times 10^{-2}$.<br>After-cue: days 1 vs 7: identical: $1.38 \times 10^{-3}$; disparate: $1.80 \times 10^{-4}$. |
| A. Rank correlation. T-test for bottom panels | Track 1, days 1 and 7:<br>Before-cue: disparate: $2.56 \times 10^{-2}$.<br>Track 2, days 1 and 7:<br>Before-cue: identical: $1.75 \times 10^{-2}$; disparate: $4.94 \times 10^{-2}$.<br>After-cue: identical: $6.80 \times 10^{-6}$; disparate: $1.32 \times 10^{-7}$. |
| B. Learning change in rank correlation. T-test for the right panels | Track 1:<br>Before-cue and after-cue: disparate: $1.51 \times 10^{-2}$.<br>Track 2:<br>Before-cue and after-cue: identical: $4.08 \times 10^{-8}$; disparate: $1.31 \times 10^{-9}$. |
| **Fig. S9. Grid and cue cells in track 3** |  |
| A. Activity variation of grid cells | Two-way ANOVA left panel: $3.66 \times 10^{-4}$.<br>T-test right panel: $2.65 \times 10^{-7}$. |
| B. Activity variation of cue cells (same as in Figure 2D) | Two-way ANOVA left panel: $6.65 \times 10^{-18}$.<br>T-test right panel: $1.45 \times 10^{-38}$. |
| C. Activity variation between grid and cue cells | Identical: $1.45 \times 10^{-37}$; disparate: $8.61 \times 10^{-5}$. |
| D. Activity variation difference between grid and cue cells | $4.26 \times 10^{-10}$ |
| E. Rank correlation of grid cells | T-test right panel: $1.70 \times 10^{-19}$. |
| F. Rank correlation of cue cells (same as in Figure 3B) | Two-way ANOVA left panel: $1.19 \times 10^{-47}$.<br>T-test right panel: $6.10 \times 10^{-160}$. |
| G. Rank correlation between grid and cue cells | Identical: $1.77 \times 10^{-169}$; disparate: $1.67 \times 10^{-13}$. |
| H. Rank correlation difference between grid and cue cells | $1.24 \times 10^{-8}$ |
| I. Decoding accuracy of grid cells | Two-way ANOVA left panel: $1.92 \times 10^{-7}$.<br>T-test right panel: $9.72 \times 10^{-12}$. |
| J. Decoding<br>accuracy of cue<br>cells | Two-way ANOVA left panel: $7.55 \times 10^{-9}$.<br>T-test right panel: $8.97 \times 10^{-25}$. |

## Materials and Methods

### Animals

All animal procedures were performed in accordance with animal protocol 1524 approved by the Institutional Animal Care and Use Committee (IACUC) at NIH/NINDS. For two-photon imaging experiments, GP5.3 mice (C57BL/6J-Tg (Thy1-GCaMP6f) GP5.3Dkim/J, JAX stock #028280)^41^ were used, including 3 males and 2 females ranging from 4-5 months old at the time when the imaging began. Mice were maintained on a reverse 12-hr on/off light schedule with all experiments being performed in the light off period.

### Surgery

#### Microprism Construction

Microprism construction procedures were similar to those described previously^20,22^. A canula (MicroGroup, 304H11XX) was attached to a circular cover glass (3mm, Warner Instruments, 64-0720). A right angle microprism coated with aluminum on the hypotenuse (1.5mm, OptoSigma, RBP3-1.5-8-550), was attached to the opposite side of the cover glass. All attachments were performed using UV-curing optical adhesive (ThorLabs, NOA81).

#### Microprism Implantation Surgery

Microprism implantation procedures were similar to those described previously^20,22^. Mice were anesthetized using a tabletop laboratory animal anesthesia system (induction: 3% isoflurane, 1L/min oxygen, maintenance: 0.5%-1.5% isoflurane, 0.7L/min oxygen, VetEquip, 901806) and surgery was performed on a stereotaxic alignment system (Kopf Instruments, 1900). A homeothermic pad and monitoring system (Harvard Apparatus, 50-7220F) was used to maintain a body temperature of 37°C. After anesthesia induction, dexamethasone (2mg/kg, VetOne, 13985-533-03) and saline (500µL, 0.9% NaCl, McKesson, 0409-4888-50) were administered by intraperitoneal (IP) injection, and slow-release buprenorphine (1mg/kg, ZooPharm, Buprenorphine SR-LAB) was administered subcutaneously. Enroflox 100 (10mg/mL, VetOne, 13985-948-10) was used as an antimicrobial wash just after the skull was exposed and just prior to sealing the skull. All insertions were performed on the left hemisphere, aligning with previous observations of more favorable vasculature on the left side^20^. A 3mm craniotomy was performed centered at 3.4mm lateral to the midline and 0.75mm posterior to the center of the transverse sinus (approximately 5.4mm posterior to the bregma). A durotomy was then performed over the cerebellum. Mannitol (3g/kg, Millipore Sigma, 63559) was administered by IP prior to the durotomy. A microprism implant, which was assembled as described above, was inserted into the transverse sinus and sealed to the skull with Vetbond (3M, 1469SB). A headplate was then mounted on the skull opposite the craniotomy. Finally, the prism and headplate were adhered to the skull with Metabond (Parkell).

### Virtual Reality

For all visual-behavioral experiments, a customized virtual reality (VR) setup was used, which projects a one-dimensional (1D) virtual environment based on the running of a mouse, similar to that described previously^22^. Mice were head-fixed onto an air-supported polystyrene ball (8” diameter, Smoothfoam) using the mounted headplate. The ball rotated on an axle, allowing only forward and backward rotation. The virtual environment was projected onto a dome screen filling the visual field of mice (270° projection). An optical flow sensor (Paialu, paiModule_10210) with infrared LEDs (DigiKey, 365-1056-ND) was used to measure the rotation of the ball and thereby control the motion of the virtual environment. The optical flow sensor output to an Arduino board (Newark, A000062), which transduced the motion signal to the computer controlling the virtual reality. An approximately 4μl water reward was provided via a lick tube at a fixed location in a given environment using a solenoid. A lick sensor connected to both the lick tube and headplate holder was used to detect mouse licking. A mouse licking the lick tube created a closed circuit between the lick sensor, the lick tube, the mouse (from the tongue to the skull), the headplate (which directly contacts the skull), and the headplate holder. The solenoid and lick sensor were controlled using a Multifunction I/O DAQ (National Instruments, PCI-6229). The virtual environments were generated and projected using ViRMEn software^42^. Imaging and behavior data were synchronized by recording a voltage signal of behavioral parameters from the VR system using the DAQ. ViRMEn environments were updated at 60Hz. The DAQ input/output rate was 1kHz. The synchronization voltage signal was updated at 20kHz. Final behavioral outputs were matched to the imaging frame rate (30Hz, see two-photon Imaging) for synchronization.

Environments were projected through a Wratten filter (Kodak, 53-700) to reduce contamination of the imaging path with projected light. Virtual environments were 1D linear tracks with patterned walls and patterned visual cues at fixed locations. At the end of the track, mice were immediately teleported to the start of the track.

### Behavior

#### Training in VR

Mice were allowed to recover for 5 days after prism implantation surgery and were then placed on water restriction, receiving 1 ml water per day. After approximately 3 days of water restriction, mice were trained daily in VR. The mice were first trained on a 1-meter track to encourage running. They were further trained on a 10-meter track with two water rewards until familiarization, as measured by consistent running (>40 trials per hour) and stable significant anticipation of reward locations for 3 days (less than 5% change in predictive licking, see “Predictive Licking” for quantification). The mice were then switched to a novel environment (track 1, 875 cm) for 7 days of navigation. On day 7 in track 1, the mice conducted a second session to explore the next novel environment (track 2, 875 cm). The mice navigated in track 2 for 7 days and on the last day, they conducted a final session (one day) to explore the third novel environment (track 3, 1000 cm). The mice were imaged daily from the first day in track 1 to the first day in track 3 (a total of 13 days). Each training or imaging session lasted around 45 mins, during which the mice traversed at least 12 runs on the track.

#### Predictive licking

Predictive licking was measured as described previously^23^. It was the percentage of licks that occurred within 20 cm prior to the reward location relative to all other locations (excluding 30 cm after reward).

### Two-photon imaging

Imaging was performed using an Ultima 2Pplus microscope (Bruker) configured with the above VR setup, as described previously^23^. A tunable laser (Coherent, Chameleon Discovery NX) set to a 920 nm excitation wavelength was used. Laser scanning was performed using a resonant-galvo scanner (Cambridge Technology, CRS8K). GCaMP fluorescence was isolated using a bandpass emission filter (525/25 nm) and detected using GaAsP photomultiplier tubes (Hamamatsu, H10770PB). A 16x water-immersion objective (Nikon, MRP07220) was used with ultrasound transmission gel (Sonigel, refractive index: 1.335989; Mettler Electronics, 1844) as the immersion media.

The anterior-posterior (AP) and the medial-lateral (ML) angle of the prism (i.e., the angle of the surface of the prism along to the AP or ML direction of the mouse) relative to the head-fixed position of the mouse were measured prior to the first imaging session. The headplate holder and rotatable objective angles were set before each imaging session to align the objective with the prism in the AP and ML direction, respectively, such that the objective was in parallel with the prism surface. A black rubber tubing was wrapped around the objective and imaging window to prevent light leakage into the objective.

Microscope control and image acquisition were performed using Prairie View software (Bruker, version 5.7). Raw data was converted to images using the Bruker Image-Block Ripping Utility. Dual-plane imaging was performed through a Z-Piezo, which enabled the objective to travel between two planes (30-µm apart) within layer 2. Imaging data in each plane were collected at 7.4 Hz and at 512 × 512 resolution (1.116 µm/pixel). Average beam power at the front of the objective was typically 70-115 mW. Imaging and behavior data were synchronized as described in “Visual Virtual Reality Setup”.

### Image Processing

#### General processing

Motion correction was performed using cross-correlation based, rigid motion correction, as described previusly^23^. Identification of regions of interest (ROIs) and the extraction of their fluorescence time course were performed using Suite2p^43^. The fractional change in fluorescence with respect to baseline (ΔF/F) was calculated as (F(t) – F0(t)) / F0(t)^20^. For each cell, significant calcium transients were identified using amplitude and duration thresholds, such that the false-positive rate of significant transient identification was 1%^27^. A final ΔF/F including only the significant calcium transients was used for all further analysis. The mean ΔF/F (significant transients only) for a cell was calculated as a function of position along the track in 5 cm bins. Data points when the mouse was moving below a speed threshold were excluded from this analysis. The speed threshold was calculated by generating a 100-point histogram of all instantaneous velocities greater than 0 and taking the value twice the center of the first bin (approximately 1% of max positive speed).

#### Cell Alignment

All imaging sessions for a given FOV were aligned pairwise using the alignment toolbox developed previously^44^ to identify common cell pairs between each pair of sessions (e.g., common cells of the sessions on days 1 and 7 in track 1). Pairwise alignments were combined to generate the full set of possible cell alignments for more than two imaging sessions (e.g., common cells on the first days in tracks 1, 2, and 3). All alignments were manually validated for accuracy.

### Data Analysis

#### Cue cells in one session

Cue scores were calculated as previously described^19^. In each track, since visual landmarks were symmetrically arranged on both sides of the track, the calculation used a single landmark template, in which spatial bins (5 cm bin) within and outside of landmark areas (between the front and back edge of each landmark) were set to 1 and 0, respectively. The cross correlation between the landmark template and the spatially binned mean ΔF/F (described in “Image Processing - General processing”) was first calculated (relative shift ≤300 cm). The peak in the cross correlation with the smallest absolute shift from zero was chosen as the best correlation of the mean ΔF/F to the landmark template. The “spatial shift” at which this peak occurred was then used to displace the landmark template to best align with the mean ΔF/F. Spatial shifts were restricted to 5cm increments, as the mean ΔF/F was calculated by every 5cm bins along the track. The correlation was then calculated locally between mean ΔF/F and each landmark within “landmark zone”, which included the landmark region and the region on either side extending by half of the landmark width. Note that the landmark zone of the same landmark could occupy different spatial bins on the mean ΔF/F of individual cells, because the landmark template could have different spatial shifts relative to the mean ΔF/F. The mean of local correlation values across all landmarks was calculated and defined as ‘cue score’ of the cell.

To identify cue cells in a particular track, a cue score threshold was calculated as the 95^th^ percentile of shuffled cue scores, which combined 200 shuffles of each cell from all FOVs of all mice and in all imaging sessions. The cells with cue scores above the threshold were identified as cue cells. Each cue cell was characterized by its “spatial shift” and a set of “landmark zones” for individual landmarks, as described above.

#### Cue cells across multiple sessions

To identify common cue cells across 2 imaging sessions (day 1 in a pair of tracks or days 1 and 7 in the same track), the same cue score calculation was conducted as described above, but the 80^th^ percentile of shuffles was used as a threshold for cue cells. All cells that passed this threshold and were also present in multiple sessions were identified as cue cells across those sessions.

#### Before-cue, at-cue, and after-cue cells in one session

Before-cue and after-cue cells were determined as the cue cells that were active before and after individual landmarks (spatial shift < 0 and > 0), respectively. At-cue cells were the cue cells with zero spatial shift.

#### Before-cue and after-cue cells across two sessions

Before-cue and after-cue cells on days 1 and 7 in the same track were the ones identified as those cells in both sessions based on the above criteria.

#### Grid cells in one session

Grid cells were identified as described previously^22,23^ based their activity features on linear tracks. For the activity of each cell, we first identified its spatial fields, which were defined by comparing the mean ΔF/F value in each 5 cm spatial bin to that of a random distribution created by 1000 bootstrapped shuffled responses^22,25^. For each 5 cm bin, a “pvalue” equaled the percent of shuffled mean ΔF/F that were higher than the real mean ΔF/F. Therefore, 1-pvalue equaled the percent of shuffled mean ΔF/F lower than the real mean ΔF/F. In-field-periods were defined as three or more adjacent bins (except at the beginning and end of the track where two adjacent bins were sufficient) whose 1-pvalue ≥ 0.8 and for which at least 20% of the runs contained significant calcium transients within the period. Out-of-field periods were defined as two or more bins whose 1-pvalue ≤ 0.25. Bins with intermediate mean ΔF/F remained unassigned.

Grid cells were further identified based on a classifier with the following criteria^22,25,26^. (1) A grid cell must have at least two spatial fields on a track. (2) The response of grid cells must have a number of transitions between an in-field and out-of-field period for a track of length L larger than L/(5w), where w is the mean field width of the response. (3) The widest field of the response must be smaller than 5w. (4) At least 30% of the bins must be assigned to either in-field or out-of-field periods. (5) The mean ΔF/F of in-field periods divided by the mean ΔF/F of out-of-field periods must be larger than 2.

Finally, we removed cue cells, which were identified in the same session, from grid cell population. To minimize the effect of cue-cell-like activity on grid cell activity in landmark encoding, we removed “cue cells” with cue scores above 80^th^ percentile of shuffles (the same threshold used in “*Cue cells across multiple sessions*”).

#### Grid cells across two sessions

Grid cells on days 1 and 7 in the same track must be the ones identified as grid cells in both sessions based on the above criteria.

#### Activity variation for a pair of landmarks

For the activity variation of a cue cell at a pair of landmarks, we first averaged the mean spatially binned ΔF/F of the cell within the “landmark zones” for the two landmarks as A1 and A2, and the activity variation was calculated as the absolute difference of A1 and A2 normalized by their sum.

The same calculation was applied to grid cells based on their spatially binned ΔF/F in “landmark zones”, which were generated during the cue score calculation (see “*Cue cells in one session*”).

#### Averaged activity variation per cell

Averaged activity variation was calculated per cell by averaging its activity variations for a certain landmark type (i.e., identical or disparate landmark pairs) at all matched distances.

#### Activity variation difference

Activity variation difference was calculated as the activity variation for disparate landmark pairs subtracted by that for identical landmark pairs. The difference is calculated for landmark pairs at individual matched distances.

#### Activity variation difference per cell

Activity variation difference was calculated per cell by subtracting its averaged activity variation at identical landmarks from that at disparate landmarks.

#### Rank correlation for a pair of landmarks

We calculated rank correlation of simultaneously imaged cue cells and grid cells in the same FOV. Since the variations in the numbers of the cells in individual FOVs could complicate the results, our calculation always included a fixed number (N) of cells randomly selected from each FOV. Only the FOVs with at least N+1 cells were involved in the calculation. Based on the numbers of cells in individual FOVs in each condition, we used a specific N for each condition so that at least half of the imaging FOVs were included in the analysis. N = 15 for cue cells on day 1 of a track (Fig. 3), for common cue cells on pairs of tracks (Fig. 5), for common cue cells on days 1 and 7 of a track (Fig. 6), and for grid and cue cells on day 1 in track 3 (Fig. S9). N = 5 for before-cue or after-cue cells on days 1 and 7 in a track (Fig. S8) and for grid and cue cells on days 1 and 7 of a track (Fig. 7). To further demonstrate that our conclusions were independent of the size of cue cell groups, for the last panels in Fig. S2 and 7, the size of cue cell groups varied from 2 to 20, and for the last panel of Fig. S5, the size of cue cell groups varied from 2 to 19 (20 could not be used because more than half of FOVs contained less than 20 cells).

Individual cell groups are unique, and the number of groups equaled to the number of total cells (M) in the FOV. To achieve this, for each cell A in the FOV, we make one random selection of N number of cells in the remaining cells (the population excluding A). We repeated this step M times.

For a pair of landmarks (landmarks 1 and 2), the rankings of the N cells, R1 and R2, were calculated based on their averaged mean ΔF/F within the “landmark zone” of landmark 1 and landmark 2, respectively. We further calculated the correlation of C1 and C2 as the “rank correlation”.

The same calculation was applied to grid cells based on their spatially binned ΔF/F in “landmark zones”, which were generated during the cue score calculation (see “*Cue cells in one session*”).

#### Averaged rank correlation per cell group

Averaged rank correlation was calculated per cell group by averaging its rank correlations for a certain landmark type (i.e., identical or disparate landmark pairs) at all matched distances.

#### Rank correlation difference

Rank correlation difference was calculated as the rank correlation for disparate landmark pairs subtracted by that for identical landmark pairs. The difference is calculated for landmark pairs at individual matched distances.

#### Rank correlation difference per cell group

Rank correlation difference was calculated per cell group by subtracting its averaged activity variation at identical landmarks from that at disparate landmarks.

#### Track change in activity variation and rank correlation for landmark pairs at different distances

For the track change of activity variations at identical landmark pairs, we used the activity variation (per cell) on one track subtracted by that on another track at individual matched distances. The track change of activity variation at disparate landmark pairs was similarly calculated. The track changes in rank correlation at different distances for identical and disparate landmark pairs were similarly calculated.

#### Learning change in activity variation and rank correlation for landmark pairs at different distances

For the learning change of activity variations at identical landmark pairs, we used the activity variation (per cell) on day 7 subtracted by that on day 1 at individual matched distances. The learning change of activity variation at disparate landmark pairs was similarly calculated. The learning changes in rank correlation at different distances for identical and disparate landmark pairs were similarly calculated.

#### Decoding accuracy for landmark decoding by temporal activity

This analysis was based on simultaneously imaged grid cells or cue cells. 40 cells were randomly picked in each FOV, and 20 random cell groups were drawn per FOV. For each landmark within landmark pair, we first took the cell activity in even runs and generated population vector (v) of at individual spatial bins (5cm per bin) within a zone that covers 50 cm before and after the centers of the landmark. 50 cm was half of the minimal distance (100 cm) between adjacent landmarks on all tracks. The vectors for landmarks 1 and 2 within the landmark pair were represented as v1 and v2, respectively. Therefore, v1 or v2 was a matrix containing 40 rows (cells) and 20 columns (spatial bins). We then compared the activity of the same cell population in odd runs at each time point within the two landmark zones (t1 and t2) with v1 and v2 by calculating the correlation between each vector in t1 and t2 and individual vectors (columns) in v1 and v2. If a temporal activity vector in t1 was best correlated with a vector in v1, this landmark encoding was correct, because the temporal activity at landmark 1 was more consistent with the population activity at the same landmark, rather than at landmark 2. Conversely, if the highest correlation was with a vector in v2, the landmark encoding was incorrect, because the temporal activity at landmark 1 was more similar to the activity at landmark 2 but not at landmark 1.

The decoding accuracy was the percentage of the time points for t1 and t2 that produced correct landmark decoding.

#### Decoding accuracy difference for landmark encoding

Decoding accuracy difference was calculated as the decoding accuracy for disparate landmark pairs subtracted by that for identical landmark pairs. The difference is calculated for landmark pairs at individual matched distances.

#### Learning change in decoding accuracy for landmark pairs at different distances

For the learning change at identical landmark pairs, we used the decoding accuracy (per random 40 cell groups) on day 7 subtracted by that on day 1 at individual matched distances. The learning change at disparate landmark pairs was similarly calculated.

### General analysis statistics

Image processing was performed using previously published MATLAB codes as cited above. The difference between two curves representing activity parameters at matched distances was calculated using two-way ANOVA. The comparison between two pairs of conditions was conducted using two-tailed Student’s t test. Linear correlations were calculated using a two-tailed Pearson’s linear correlation coefficient. P values less than 0.05 were considered significant (* < 0.05, ** <0.01, *** <0.001). All figures show mean and standard error, except where noted.

## References

1. Chan, E., Baumann, O., Bellgrove, M.A., and Mattingley, J.B. (2012). From objects to landmarks: the function of visual location information in spatial navigation. Front Psychol 3, 304. 10.3389/fpsyg.2012.00304.

2. Eichenbaum, H., Yonelinas, A.P., and Ranganath, C. (2007). The medial temporal lobe and recognition memory. Annu Rev Neurosci 30, 123–152. 10.1146/annurev.neuro.30.051606.094328.

3. Mishkin, M.U.L., Macko L. (1983). Object vision and spatial vision: two cortical pathways. Trends in Neurosciences 6, 414–417, aga.

4. Witter, M.P., Doan, T.P., Jacobsen, B., Nilssen, E.S., and Ohara, S. (2017). Architecture of the Entorhinal Cortex A Review of Entorhinal Anatomy in Rodents with Some Comparative Notes. Front Syst Neurosci 11, 46. 10.3389/fnsys.2017.00046.

5. Tennant, S.A., Fischer, L., Garden, D.L.F., Gerlei, K.Z., Martinez-Gonzalez, C., McClure, C., Wood, E.R., and Nolan, M.F. (2018). Stellate Cells in the Medial Entorhinal Cortex Are Required for Spatial Learning. Cell Rep 22, 1313–1324. 10.1016/j.celrep.2018.01.005.

6. Van Cauter, T., Camon, J., Alvernhe, A., Elduayen, C., Sargolini, F., and Save, E. (2013). Distinct roles of medial and lateral entorhinal cortex in spatial cognition. Cereb Cortex 23, 451–459 10.1093/cercor/bhs033.

7. Jacob, P.Y., Gordillo-Salas, M., Facchini, J., Poucet, B., Save, E., and Sargolini, F. (2017). Medial entorhinal cortex and medial septum contribute to self-motion-based linear distance estimation. Brain Struct Funct 222, 2727–2742 10.1007/s00429-017-1368-4.

8. Hales, J.B., Schlesiger, M.I., Leutgeb, J.K., Squire, L.R., Leutgeb, S., and Clark, R.E. (2014). Medial entorhinal cortex lesions only partially disrupt hippocampal place cells and hippocampus-dependent place memory. Cell Rep 9, 893–901. 10.1016/j.celrep.2014.10.009.

9. Ferbinteanu, J., Holsinger, R.M., and McDonald, R.J. (1999). Lesions of the medial or lateral perforant path have different effects on hippocampal contributions to place learning and on fear conditioning to context. Behav Brain Res 101, 65–84. 10.1016/s0166-4328(98)00144-2.

10. Hafting, T., Fyhn, M., Molden, S., Moser, M.B., and Moser, E.I. (2005). Microstructure of a spatial map in the entorhinal cortex. Nature 436, 801–806. 10.1038/nature03721.

11. Sargolini, F., Fyhn, M., Hafting, T., McNaughton, B.L., Witter, M.P., Moser, M.B., and Moser, E.I. (2006). Conjunctive representation of position, direction, and velocity in entorhinal cortex. Science 312, 758–762. 10.1126/science.1125572.

12. Solstad, T., Boccara, C.N., Kropff, E., Moser, M.B., and Moser, E.I. (2008). Representation of geometric borders in the entorhinal cortex. Science 322, 1865–1868. 10.1126/science.1166466.

13. Rodo, C., Sargolini, F., and Save, E. (2017). Processing of spatial and non-spatial information in rats with lesions of the medial and lateral entorhinal cortex: Environmental complexity matters. Behav Brain Res 320, 200–209. 10.1016/j.bbr.2016.12.009.

14. Hunsaker, M.R., Chen, V., Tran, G.T., and Kesner, R.P. (2013). The medial and lateral entorhinal cortex both contribute to contextual and item recognition memory: a test of the binding of items and context model. Hippocampus 23, 380–391. 10.1002/hipo.22097.

15. Beer, Z., Chwiesko, C., Kitsukawa, T., and Sauvage, M.M. (2013). Spatial and stimulus- type tuning in the LEC, MEC, POR, PrC, CA1, and CA3 during spontaneous item recognition memory. Hippocampus *23*, 1425-1438. 10.1002/hipo.22195.

16. Nguyen, D., Wang, G., Wafa, T., Fitzgerald, T., and Gu, Y. (2024). The medial entorhinal cortex encodes multisensory spatial information. Cell Rep 43, 114813. 10.1016/j.celrep.2024.114813.

17. Keene, C.S., Bladon, J., McKenzie, S., Liu, C.D., O’Keefe, J., and Eichenbaum, H. (2016). Complementary Functional Organization of Neuronal Activity Patterns in the Perirhinal, Lateral Entorhinal, and Medial Entorhinal Cortices. J Neurosci 36, 3660–3675. 10.1523/JNEUROSCI.4368-15.2016.

18. Hoydal, O.A., Skytoen, E.R., Andersson, S.O., Moser, M.B., and Moser, E.I. (2019). Object-vector coding in the medial entorhinal cortex. Nature 568, 400–404. 10.1038/s41586-019-1077-7.

19. Kinkhabwala, A.A., Gu, Y., Aronov, D., and Tank, D.W. (2020). Visual cue-related activity of cells in the medial entorhinal cortex during navigation in virtual reality. Elife 9. 10.7554/eLife.43140.

20. Low, R.J., Gu, Y., and Tank, D.W. (2014). Cellular resolution optical access to brain regions in fissures: imaging medial prefrontal cortex and grid cells in entorhinal cortex. Proc Natl Acad Sci U S A 111, 18739–18744. 10.1073/pnas.1421753111.

21. Chen, T.W., Wardill, T.J., Sun, Y., Pulver, S.R., Renninger, S.L., Baohan, A., Schreiter, E.R., Kerr, R.A., Orger, M.B., Jayaraman, V., et al. (2013). Ultrasensitive fluorescent proteins for imaging neuronal activity. Nature 499, 295–300. 10.1038/nature12354.

22. Gu, Y., Lewallen, S., Kinkhabwala, A.A., Domnisoru, C., Yoon, K., Gauthier, J.L., Fiete, I.R., and Tank, D.W. (2018). A Map-like Micro-Organization of Grid Cells in the Medial Entorhinal Cortex. Cell 175, 736–750 e730. 10.1016/j.cell.2018.08.066.

23. Malone, T.J., Tien, N.W., Ma, Y., Cui, L., Lyu, S., Wang, G., Nguyen, D., Zhang, K., Myroshnychenko, M.V., Tyan, J., et al. (2024). A consistent map in the medial entorhinal cortex supports spatial memory. Nat Commun 15, 1457. 10.1038/s41467-024-45853-4.

24. Cohen, J.D., Bolstad, M., and Lee, A.K. (2017). Experience-dependent shaping of hippocampal CA1 intracellular activity in novel and familiar environments. Elife 6. 10.7554/eLife.23040.

25. Domnisoru, C., Kinkhabwala, A.A., and Tank, D.W. (2013). Membrane potential dynamics of grid cells. Nature 495, 199–204. 10.1038/nature11973.

26. Yoon, K., Lewallen, S., Kinkhabwala, A.A., Tank, D.W., and Fiete, I.R. (2016). Grid Cell Responses in 1D Environments Assessed as Slices through a 2D Lattice. Neuron 89, 1086–1099. 10.1016/j.neuron.2016.01.039.

27. Heys, J.G., Rangarajan, K.V., and Dombeck, D.A. (2014). The functional micro-organization of grid cells revealed by cellular-resolution imaging. Neuron 84, 1079–1090. 10.1016/j.neuron.2014.10.048.

28. Deshmukh, S.S., and Knierim, J.J. (2011). Representation of non-spatial and spatial information in the lateral entorhinal cortex. Front Behav Neurosci 5, 69. 10.3389/fnbeh.2011.00069.

29. Huang, X., Schlesiger, M.I., Barriuso-Ortega, I., Leibold, C., MacLaren, D.A.A., Bieber, N., and Monyer, H. (2023). Distinct spatial maps and multiple object codes in the lateral entorhinal cortex. Neuron 111, 3068–3083 e3067. 10.1016/j.neuron.2023.06.020.

30. Weible, A.P., Rowland, D.C., Monaghan, C.K., Wolfgang, N.T., and Kentros, C.G. (2012). Neural correlates of long-term object memory in the mouse anterior cingulate cortex. J Neurosci 32, 5598–5608. 10.1523/JNEUROSCI.5265-11.2012.

31. Deshmukh, S.S., Johnson, J.L., and Knierim, J.J. (2012). Perirhinal cortex represents nonspatial, but not spatial, information in rats foraging in the presence of objects: comparison with lateral entorhinal cortex. Hippocampus 22, 2045–2058. 10.1002/hipo.22046.

32. Deshmukh, S.S., and Knierim, J.J. (2013). Influence of local objects on hippocampal representations: Landmark vectors and memory. Hippocampus 23, 253–267. 10.1002/hipo.22101.

33. Geiller, T., Fattahi, M., Choi, J.S., and Royer, S. (2017). Place cells are more strongly tied to landmarks in deep than in superficial CA1. Nat Commun 8, 14531. 10.1038/ncomms14531.

34. Lever, C., Burton, S., Jeewajee, A., O’Keefe, J., and Burgess, N. (2009). Boundary vector cells in the subiculum of the hippocampal formation. J Neurosci 29, 9771–9777. 10.1523/JNEUROSCI.1319-09.2009.

35. Tsao, A., Moser, M.B., and Moser, E.I. (2013). Traces of experience in the lateral entorhinal cortex. Curr Biol 23, 399–405. 10.1016/j.cub.2013.01.036.

36. Doan, T.P., Lagartos-Donate, M.J., Nilssen, E.S., Ohara, S., and Witter, M.P. (2019). Convergent Projections from Perirhinal and Postrhinal Cortices Suggest a Multisensory Nature of Lateral, but Not Medial, Entorhinal Cortex. Cell Rep 29, 617–627 e617. 10.1016/j.celrep.2019.09.005.

37. Wilson, D.I., Watanabe, S., Milner, H., and Ainge, J.A. (2013). Lateral entorhinal cortex is necessary for associative but not nonassociative recognition memory. Hippocampus 23, 1280–1290. 10.1002/hipo.22165.

38. Wilson, D.I., Langston, R.F., Schlesiger, M.I., Wagner, M., Watanabe, S., and Ainge, J.A. (2013). Lateral entorhinal cortex is critical for novel object-context recognition. Hippocampus 23, 352–366. 10.1002/hipo.22095.

39. Vandrey, B., Garden, D.L.F., Ambrozova, V., McClure, C., Nolan, M.F., and Ainge, J.A. (2020). Fan Cells in Layer 2 of the Lateral Entorhinal Cortex Are Critical for Episodic-like Memory. Curr Biol 30, 169–175 e165. 10.1016/j.cub.2019.11.027.

40. Yoganarasimha, D., Rao, G., and Knierim, J.J. (2011). Lateral entorhinal neurons are not spatially selective in cue-rich environments. Hippocampus 21, 1363–1374. 10.1002/hipo.20839.

41. Dana, H., Chen, T.W., Hu, A., Shields, B.C., Guo, C., Looger, L.L., Kim, D.S., and Svoboda, K. (2014). Thy1-GCaMP6 transgenic mice for neuronal population imaging in vivo. PLoS One 9, e108697. 10.1371/journal.pone.0108697.

42. Aronov, D., Nevers, R., and Tank, D.W. (2017). Mapping of a non-spatial dimension by the hippocampal-entorhinal circuit. Nature 543, 719–722. 10.1038/nature21692.

43. Pachitariu, M., Stringer, C., Dipoppa, M., Schroder, S., Rossi, L. F., Dalgleish, H., Caradini, M., Harris, K. D. (2017). Suite2p: beyond 10,000 neurons with standard two-photon microscopy. bioRxiv.

44. Sheintuch, L., Rubin, A., Brande-Eilat, N., Geva, N., Sadeh, N., Pinchasof, O., and Ziv, Y. (2017). Tracking the Same Neurons across Multiple Days in Ca(2+) Imaging Data. Cell Rep 21, 1102–1115. 10.1016/j.celrep.2017.10.013.

